# Discovery of a functionally selective serotonin 5-HT_1A_ receptor agonist for the treatment of pain

**DOI:** 10.1101/2023.09.11.557127

**Authors:** Annika Ullrich, Johannes Schneider, João M. Braz, Eduard Neu, Nico Staffen, Markus Stanek, Jana Bláhová, Tamsanqa Hove, Tamara Albert, Anni Allikalt, Stefan Löber, Karnika Bhardwaj, Sian Rodriguez-Rosado, Elissa Fink, Tim Rasmussen, Harald Hübner, Asuka Inoue, Brian K. Shoichet, Allan J. Basbaum, Bettina Böttcher, Dorothee Weikert, Peter Gmeiner

**Affiliations:** Department of Chemistry and Pharmacy, Friedrich-Alexander-Universität Erlangen-Nürnberg, 91058 Erlangen, Germany; FAU NeW, 91058 Erlangen, Germany; Department of Anatomy, University of California, San Francisco, CA, USA; Rudolf-Virchow-Center, Julius-Maximilians-Universität, 97080 Würzburg, Germany; Department of Pharmaceutical Chemistry, University of California, San Francisco, CA, USA; Graduate School of Pharmaceutical Sciences, Tohoku University, Sendai, 980-8578, Japan

## Abstract

The G protein-coupled serotonin receptor 5-HT_1A_R mediates antinociception and may serve as a valuable target for the treatment of pain. Starting from a chemical library, ST171, a bitopic chemotype activating 5-HT_1A_R was evolved. *In vitro* pharmacological investigations of ST171 revealed potent and selective G_i_ activation (EC_50_ = 0.3 nM), with marginal G_s_ and β-arrestin recruitment. Preclinical studies in mice showed that ST171 was effective in acute and chronic (inflammatory and neuropathic) pain models, without causing sedation. Comparison of cryo-EM structures of a 5-HT_1A_R-G_i_ complex bound to the functionally biased agonist ST171, with a structure bound to the functionally balanced agonist befiradol, showed that both ligands bind to the same orthosteric site, but address different exo-sites. The individual poses are associated with ligand-specific helical dispositions and rearrangements of microdomains. Complementation of these studies with molecular dynamics simulations allowed us to derive structural features associated with ST171’s functional selectivity, a phenomenon that may be crucial to the discovery of more effective and safe GPCR drugs.

## Introduction

The treatment of pain by opioids is difficult because they induce adverse effects, including addiction, constipation, and respiratory depression (*1, 2*). Efforts have been dedicated to the discovery of novel opioids, including opioid receptor partial agonist (*3*), and functionally selective ligands that are devoid of these side effects (*4, 5*). Other approaches sought analgesics that target non-opoiod sensitive links in pain processing circuitry.

Of interest in this approach is the serotonin 5-HT_1A_ receptor (5-HT_1A_R), which is a valuable target for the treatment of anxiety and depressive disorders (*6*). Importantly, the 5-HT_1A_R, is also highly expressed and can inhibit nociceptive processing circuits at different levels of the neuraxis (*7, 8*). For example an action at the 5-HT_1A_R results in analgesia with inverse tolerance (*9*), a decrease of opioid-induced rewarding effects (*10*) and reduced respiratory depression (*11*). Of particular interest is befiradol, a selective 5-HT_1A_R agonist that has analgesic properties comparable to clinically used opioids (*12, 13*). Although befiradol can also reverse opioid-induced respiratory depression in rats, this reversal often occurs with concurrent hyperalgesia (*11*) and sedation (*14*). Those adverse side effects led to termination of clinical trials for the treatment of neuropathic and cancer pain. To leverage the potential of the 5-HT_1A_R as an antinociceptive target, here we sought novel chemotypes for the 5-HT_1A_R, with functional selectivity for G_i_ signaling. Our objective is to discover novel, efficacious non-opioid analgesic drugs without the hyperalgesic and sedative effects of befiradol.

## Results

### Discovery of ST171

We have recently shown that the discovery of new chemotypes is a very powerful strategy to find receptor ligands with preferential G_i_ signaling and promising *in vivo* analgesic activity (*15*). Hence, we could generate new lead compounds for effective and safe therapeutics.

To identify new chemotypes for the 5-HT_1A_R, we inspected our in-house library with approximately 10,000 compounds for potential binders at the 5-HT_1A_R (Figure 1B). The library included 2,000 FDA-approved drugs for addressing targets beyond GPCRs and bioactive compounds generated in our recent GPCR projects. In a screening of the FDA-approved drugs and a sub-library that includes a set of structurally diverse aminergic receptor ligands, the 5-HT_1A_R affinity of ST162, a compound that was part of an earlier dopamine receptor program, attracted our attention. Evaluation of 60 structural analogs of ST162 for their ability to displace a radiolabeled ligand ([³H]WAY100,635) from the 5-HT_1A_R, led us to ST171, as it displaced more than 80 % radioligand at 10 nM and, even at 1 nM, specific binding was observed. Compared to serotonin and befiradol, ST171 showed superior binding affinity for the 5-HT_1A_R in concentration-response assays with a 680-fold subtype selectivity over 5-HT_2A_R as a representative of the G_q_-coupled subtypes and an 8,000-fold selectivity over 5-HT_6_R as a G_s_-coupled family member (Figure 1C; 5-HT_1A_R: *K_i_* [ST171] = 0.41 ± 0.06 nM, *K_i_* [serotonin] = 220 ± 52 nM, *K_i_* [befiradol] = 15 ± 3.4 nM; 5-HT_2A_R: *K_i_* [ST171] = 280 ± 30 nM, *K_i_* [serotonin] = 130 ± 22 nM, *K_i_* [befiradol] = 15,000 ± 3,700 nM; 5-HT_6_R: *K_i_* [ST171] = 3,300 ± 300 nM, *K_i_* [serotonin] = 140 ± 37 nM, *K_i_* [befiradol] = >50,000 nM). (Supplementary Table 1)

**Figure 1:**
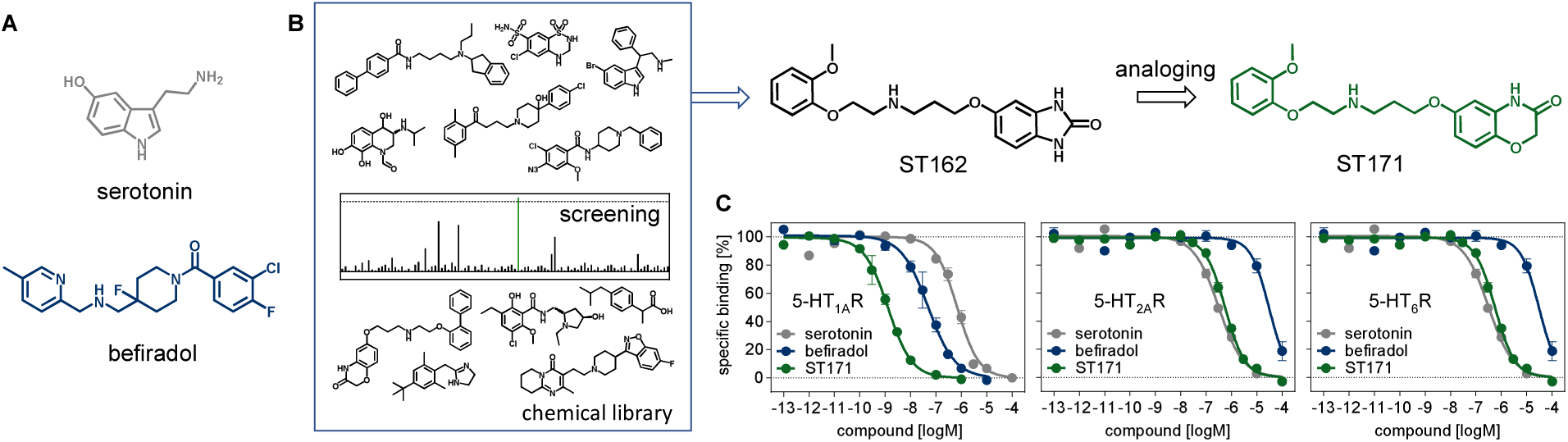
Discovery of ST171. (A) Chemical structures of the reference agents serotonin and befiradol. (B) In-house library investigation of binding affinity at the 5-HT_1A_R using radioligand binding studies. (C) Comparison of the binding affinities of serotonin, befiradol, and ST171 at 5-HT_1A_R (displacement of [^3^H]WAY100635), 5-HT_2A_R (displacement of [^3^H]ketanserin), and 5-HT_6_R (displacement of [^3^H]LSD). Data is shown with ± SEM of 4-11 independent experiments, each performed as triplicates.

### ST171 is a functionally selective agonist of the 5-HT_1A_R

*In vitro* functional studies were initiated to explore the signaling profile of ST171 compared to serotonin and befiradol resulting from interaction with the predominantly G_i/o_-coupled 5-HT_1A_R (*16*). BRET-based sensing of cAMP (CAMYEL biosensor) in CHO-K1 cells showed inhibition of cAMP formation in response to treatment with the agonists serotonin and befiradol. Interestingly, we observed bell-shaped concentration-response curves indicating reduced inhibition of adenylyl cyclase at higher concentrations of these agonists, and thus suggesting G_s_- in competition with the predominant G_i/o_-coupling (Figure 2A; serotonin: EC_50_^left^ = 3.6 ± 0.6 nM, EC_50_^right^ = 240 ± 70 nM, E_max_ = 100 %, E_sat_ = 64 ± 3 %; befiradol: EC_50_^left^ = 1.8 ± 0.3 nM, EC_50_^right^ = 110 ± 1 nM, E_max_ = 91 ± 3 %, E_sat_ = 65 ± 4 %). Bell-shaped concentration-response curves were also obtained in a cAMP GloSensor assay in HEK293A, HEK293T and HEK β-arrestin knockout (Δβarr) cells (*17*) (Supplementary Figure 1A-C). In the presence of pertussis toxin (PTX) inhibiting the interaction of G_i/o_ proteins with GPCRs (*18, 19*), stimulation with serotonin or befiradol increased intracellular cAMP levels with a sigmoidal concentration-response curve (Figure 2A; serotonin: EC_50_ = 52 ± 6 nM, befiradol: EC_50_ = 25 ± 11 nM). Serotonin and befiradol inhibited cAMP accumulation (GloSensor) in HEK293A cells deficient of Gα_s_ proteins (*20*), while the compounds stimulated cAMP accumulation in HEK293A cells deficient of Gα_i/o_ (*21*) (Figure 2B and C; serotonin: EC_50_[ΔG_i_] = 240 ± 62 nM, EC_50_[ΔG_s_] = 2.0 ± 0.5 nM; befiradol: EC_50_[ΔG_i_] = 270 ± 65 nM, E_max_[ΔG_i_] = 70 ± 2 %; EC_50_[ΔG_s_] = 2.3 ± 0.6 nM, E_max_[ΔG_s_] = 85 ± 5 %), confirming that G_s_ and G_i/o_ mediate the accumulation and inhibition, respectively, of the bell-shape response upon stimulation with serotonin or befiradol. Co-transfection of the different Gα_i/o_ subunits rescued Gα_i/o_-mediated inhibition of cAMP accumulation in the HEK293AΔG_i_ cells (Supplementary Figure 1E). In agreement with the activation of different G proteins, serotonin and befiradol also promoted a G_i_ and G_s_ protein-mediated formation of intracellular inositol phosphate via hybrid Gα_qi_ and Gα_qs_ proteins, respectively (Supplementary Figure 1D; serotonin: EC_50_[G_qi_] = 58 ± 9.4 nM, EC_50_[G_qs_] = 420 ± 70 nM; befiradol: EC_50_[G_qi_] = 12 ± 1.3 nM, E_max_[G_qi_] = 92 ± 3 %; EC_50_[G_qs_] = 53 ± 3.7 nM, E_max_[G_qs_] = 83 ± 6 %).

**Figure 2:**
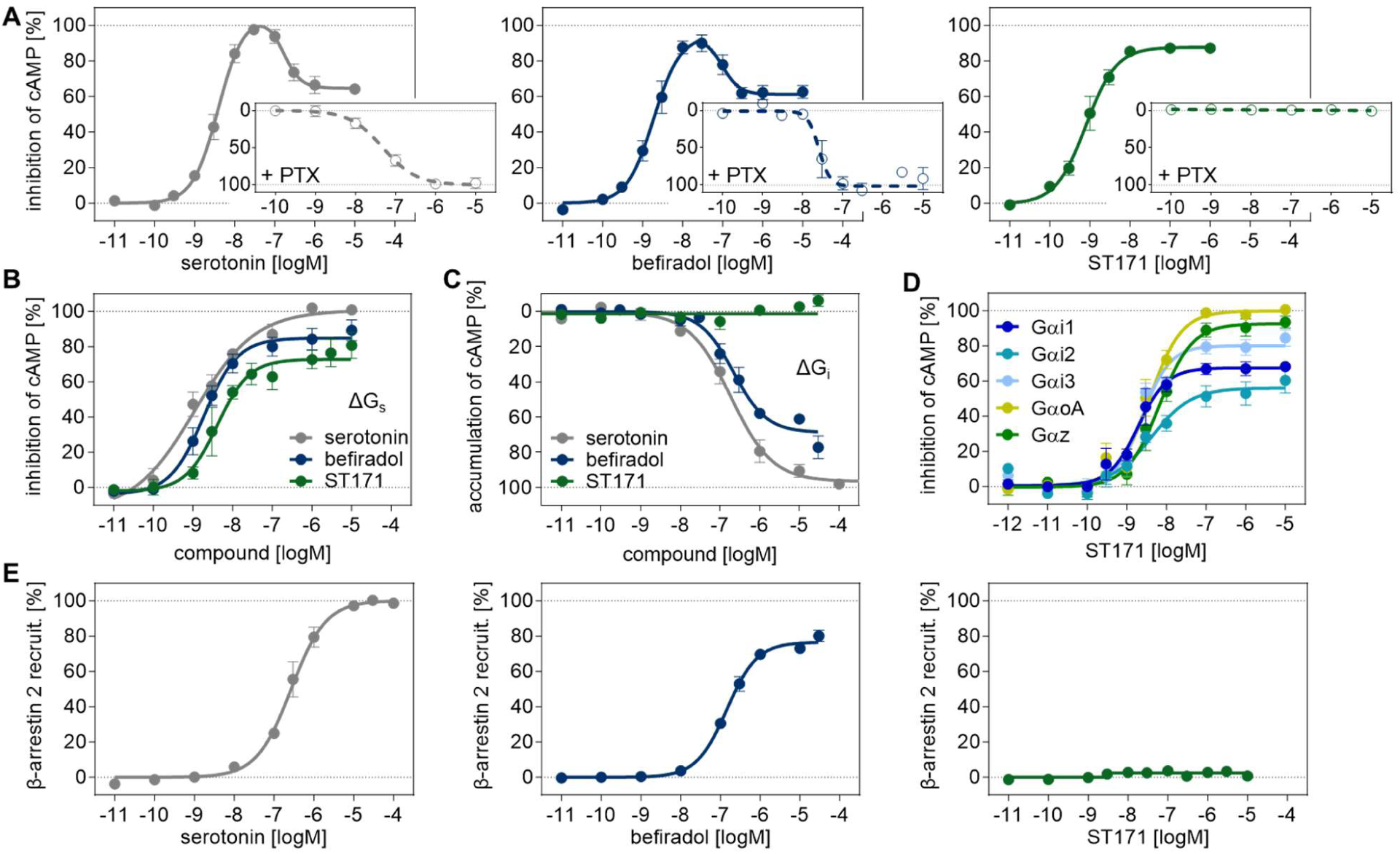
Functional characterization of serotonin, befiradol and ST171. (A) Inhibition of cAMP accumulation by serotonin, befiradol, and ST171 in the presence and absence of PTX. Data was obtained with a CAMYEL assay in CHO-K1 cells and is normalized to the maximal response of serotonin. Data is shown with ± SEM of 3-5 independent experiments, each performed as triplicates. (B) Inhibition of cAMP accumulation by serotonin, befiradol, and ST171 in (B) HEK293A cells deficient of G_s_ proteins and (C) accumulation of cAMP by serotonin, befiradol, and ST171 in HEK293A cells deficient of G_i/o_ proteins. Data was obtained with a GloSensor, normalized to the maximal response of serotonin and is shown with ± SEM of 6 experiments, each performed as triplicates. (D) Inhibition of cAMP accumulation by ST171 in HEK293AΔG_i_ cells co-transfected with Gα_i_ subunits (Gα_i1_, Gα_i2_, Gα_i3_, Gα_z_, Gα_oA_). Data was obtained with the GloSensor, normalized to the maximal response of serotonin and is shown with ± SEM of 5 experiments, each performed as triplicates. (E) β-arrestin 2 recruitment monitored by bystander BRET in HEK293T cells with elevated GRK2 levels. Data is normalized to the maximal response of serotonin and shown with ± SEM of 5-6 independent experiments, each performed as duplicates.

ST171 inhibited cAMP accumulation with high efficacy and subnanomolar potency (Figure 2A; EC_50_ = 0.88 ± 0.3 nM, E_max_ = 87 ± 1 % in CHO-K1 cells and Supplementary Figure 1B and 1C, Supplementary Table 2). Notably, unlike serotonin and befiradol, the concentration-response curve was sigmoidal and PTX addition completely blocked the effect indicating selective signaling through G_i/o_ proteins over G_s_ (Figure 2A, E_max_ < 5 %). ST171 did not elicit an effect in HEK293A cells lacking Gα_i/o_, while knockout of G_s_ had no effect on ligand efficacy (Figure 2B and C; E_max_[ΔG_i_] < 5 %; EC_50_[ΔG_s_] = 6.4 ± 1.4 nM, E_max_[ΔG_s_] = 73 ± 4 %) confirming functional selectivity over G_s_. Furthermore, intracellular IP accumulation in the presence of the hybrid Gα_qs_ protein could not be observed (Supplementary Figure 1D, E_max_[G_qs_] < 5 %). Overexpression of the different Gα_i/o_ subtypes Gα_i1_, Gα_i2_, Gα_i3_, Gα_z_, or Gα_oA_ rescued the inhibition of cAMP formation by ST171 in HEK293AΔG_i_ cells and interestingly revealed a preference for the activation of Gα_oA_ (E_max_ 100 ± 3 %) and Gα_z_ (E_max_ 93 ± 4 %) over the other Gα_i_ subtypes (E_max_ 57-80 %, Figure 2D, Supplementary Table 3). In HEK β-arrestin knockout cells (cAMP Glosensor, Supplementary Figure 1A, E_max_ = 60 ± 3 %) and the IP accumulation assay in the presence of Gα_qi_ (Supplementary Figure 1D, E_max_ = 43 ± 2%) ST171 showed partial agonist activity. We confirmed the partial agonist effect of ST171 in a BRET biosensor-based Gα_i1_ activation assay at different receptor expression levels. While serotonin and befiradol showed almost identical efficacies and only a minor reduction in potency at low receptor levels, the intrinsic activity of ST171 diminished gradually, from 77 % to about 37 % with decreasing 5-HT_1A_R expression (Supplementary Figure 1G, Supplementary Table 4).

Stimulation of the 5-HT_1A_R with serotonin and befiradol promoted the recruitment of β-arrestin 2, which was determined by enhanced bystander BRET using a *Renilla* Luciferase tagged β-arrestin 2 and a membrane-anchored *Renilla* GFP (*22*) with co-transfected GRK2 (Figure 2E; serotonin: EC_50_ = 320 ± 32 nM; befiradol: 150 ± 11 nM, E_max_ = 77 ± 1 %). The importance of GRK2 for the β-arrestin 2 recruitment was determined with a highly sensitive enzyme complementation-based assay (PathHunter) with native and transiently coexpressed GRK2, showing that elevated GRK2 results in a 9-fold (serotonin) and 4-fold (befiradol) increase of potency (Supplementary Figure 2; serotonin: EC_50_[native GRK2] = 620 ± 35 nM, EC_50_[elevated GRK2] = 67 ± 4.8 nM; befiradol: EC_50_[native GRK2] = 690 ± 120 nM, EC_50_[elevated GRK2] = 180 ± 47 nM, E_max_[native GRK2] = 67 ± 3 %, E_max_[elevated GRK2] = 96 ± 3 %).

In contrast to serotonin and befiradol, ST171 was not able to stimulate the recruitment of β-arrestin 2 in both the bystander BRET (Figure 2E; E_max_ < 5 %) and the complementation-based assay with native GRK2 and only showed minor recruitment with elevated GRK2 levels (Supplementary Figure 2; E_max_[native GRK2] < 5 %, EC_50_[elevated GRK2] = 5.5 ± 1.4 nM, E_max_[elevated GRK2] = 15 ± 1 %).

Taken together, we identified ST171 as a highly potent 5-HT_1A_R agonist with functional selectivity for G_i/o_ proteins over G_s_ and β-arrestin 2.

ST171 showed low to moderate affinity for a set of 23 other class A GPCRs, but high affinity for the dopamine D4 and the α_1A_-adrenergic receptor (Supplementary Figure 4, Supplementary Table 6). As α_1_AR agonism is associated with hypotension and bradycardia (*15, 23*), we investigated ligand efficacy of ST171 in an IP accumulation assay (α_1_ARs). The data indicated that ST171 was devoid of G_q_-promoted intrinsic activity at α_1_AR while partial agonist activity was measured for D_4_ in a BRET Gα_i1_ dissociation assay (EC_50_ = 1.6 ± 0.22 nM, E_max_ = 42 ± 3%, Supplementary Figure 4A-C, Supplementary Table 6). Using the BRET Gα_i1_ dissociation assay we also demonstrated absence of agonist activity of ST171 at α_2A_AR, which is an important alternative target for pain alleviation (*15*), or for the related α_2B_AR and α_2C_AR subtypes (Suppelmentary Figure 4D-F, Supplementary Table 6).

### ST171 reduces hypersensitivity in mouse models of chronic (neuropathic and inflammatory) pain

Although serotonin (5-HT) can modulate a wide variety of pain states, from migraine to inflammatory and neuropathic pain, because 5-HT exerts a bidirectional action, its overall contribution to nociceptive processing and/or analgesia is not straightforward (*8, 24, 25*). Not only can 5-HT exert both pro- (*26, 27*) and anti-inflammatory effects (*28*), but both pro- (*29, 30*) antinociceptive actions have also been been reported (*31–33*). Mediating these opposing actions are at least 7 subtypes of 5-HT receptors, which are expressed throughout the CNS and PNS (*34*). Comparable paradoxical actions at the G_i/o_-coupled 5-HT_1A_R subtype have also been reported (*35–38*), possibly due to differences in pain assays (*39, 40*) or mode of drug administration (*37, 41, 42*). The latter influence is significant as the 5-HT_1A_R is widely expressed (*43, 44*), including primary sensory neurons of the dorsal root ganglia (DRG) and in spinal cord (*45, 46*).

Here we tested the analgesic properties of our novel and selective 5-HT_1A_R agonist, ST171, in both acute and chronic (neuropathic and inflammatory) pain conditions. In these studies, we used the 10 mg/kg intraperitoneal (IP) dose of ST171, which showed favorable pharmacokinetics. We compared the results to the action of befiradol, also at 10 mg/kg (*38*). We first focused on acute mechanical and thermal pain assays. Figure 3 illustrates that a systemic injection of both ST171 and befiradol were analgesic in both assays, significantly increasing mechanical thresholds (Fig 3A) as well as the response latency to thermal stimuli in 3 different tests of “heat pain” (Fig 3C-E). Importantly, when tested for sedative effects using an accelerating rotorod, mice injected with 10 mg/kg ST171 did not differ from control mice injected with vehicle (Fig. 3B). Only at a 15 mg kg dose of ST171 did we record a sedative effect on the rotarod. In contrast, the 10 mg/kg dose of befiradol was clearly sedating, which suggests that the motor deficits may have contributed to its antinociceptive effects. Importantly, biferadol administered at 5 mg/kg was still sedating and interestingly, a non-sedating dose (1 mg/kg) was only antinociceptive in acute heat pain tests. There was no reduction of the response to noxious mechanical stimulation. Therapeutic window is clearly significantly greater for ST171.

**Figure 3:**
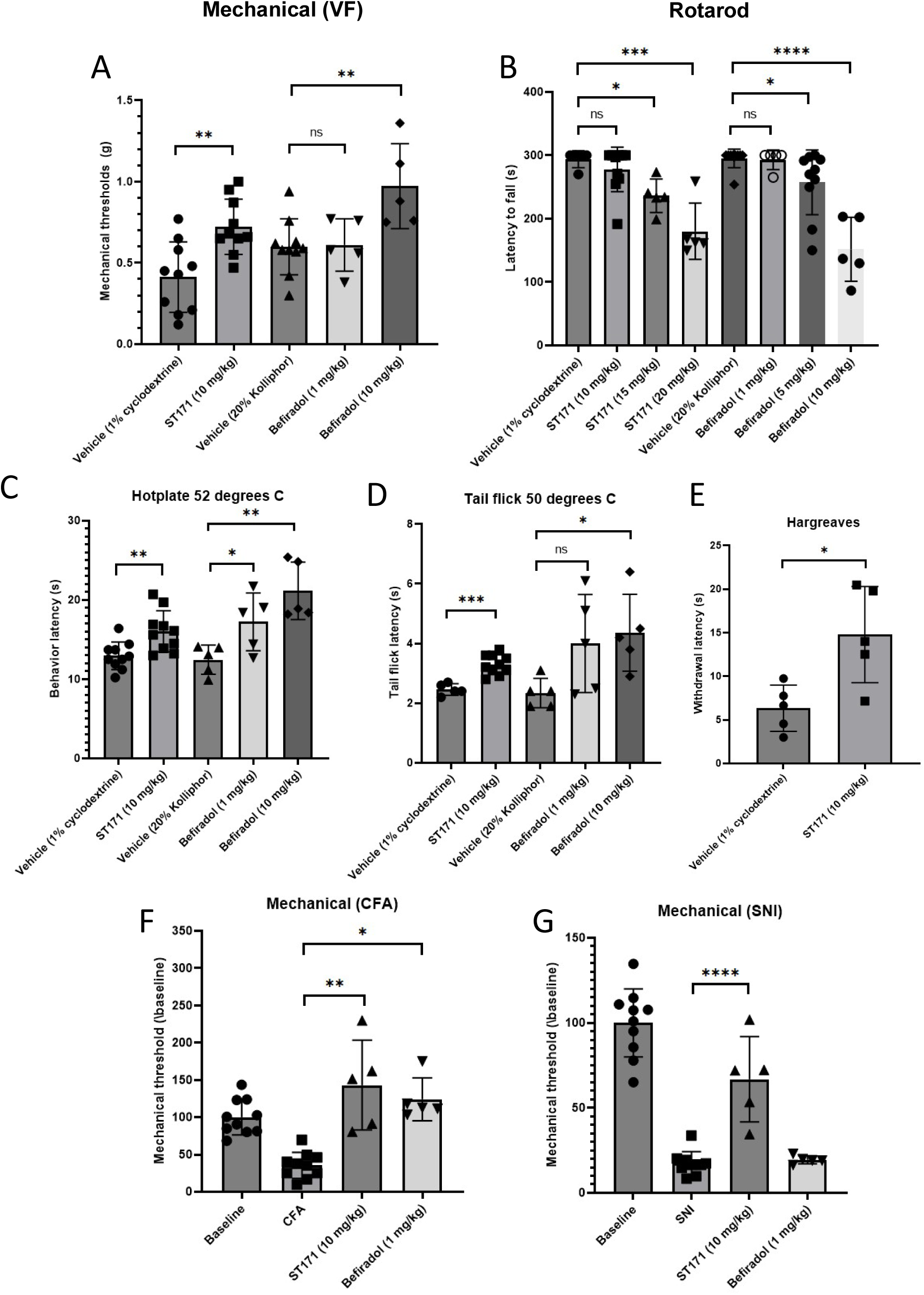
ST171 is analgesic in acute pain tests and anti-hyperalgesic in mouse models of chronic (neuropathic and inflammatory) pain. (**A**) *In vivo* efficacy of ST171 and befiradol in the von Frey assay. (**B**) Effect of increasing doses of ST171 and befiradol on motor performances in the rotarod assay. (**C-E**) Analgesic effects of ST171 and befiradol in different thermal assays: 52°C hot plate (**C**), 50°C tail flick (**D**) and Hargreaves (**E**). **(F)** Anti-hyperalgesic effects of ST171 and befiradol in the Complete Freund’s adjuvant (CFA)-induced inflammatory pain model. (**G**) Anti-allodynic effects of ST171 and befiradol in the spared nerve injury (SNI) model of neuropathic pain. All data represent Mean ± SEM of 5-10 animals. The effects of ST171 and befiradol were compared to their respective vehicle using a Student’s t-test *p<0.05; **p<0.01; ***p<0.005; ****p<0.001.

**Figure 4:**
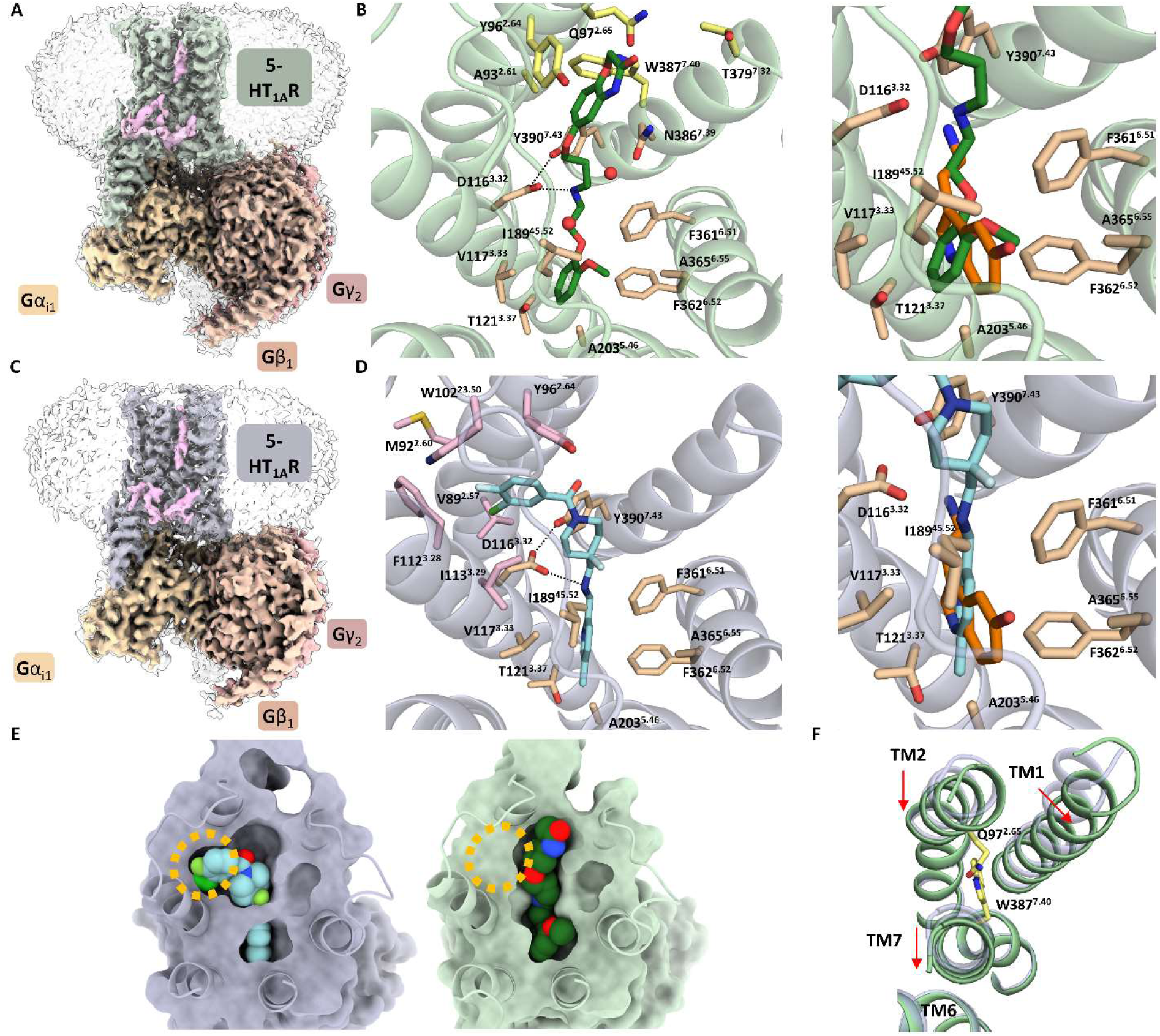
Cryo-EM structure of the ST171 and befiradol-bound 5-HT_1A_R-G_i_ Protein complex. (A) Cryo-EM map of the ST171-bound 5-HT_1A_R-G_i_ protein complex with cholesterols and PI_4_P (light pink). (B) Top view of ST171 binding pose and close up comparison to the serotonin binding mode (PDB:7E2Y, orange). (C) Cryo-EM map of the befiradol-bound 5-HT_1A_R-G_i_ Protein complex with cholesterol and PI_4_P (light pink). (D) Top view of befiradol binding pose and close up comparison to the serotonin binding mode (PDB:7E2Y, orange). (E) Surface representation of befiradol (violet) and ST171 (light green) bound 5-HT_1A_R. (F) Inward shift of TM2 and outward shift of TM1 and TM7 of ST171-bound 5-HT_1A_R structure (light green) in comparison to the befiradol-bound structure (light blue) (extracellular view).

We next assessed the efficacy of ST171 and befiradol in two models of chronic pain, namely the spared nerve injury (SNI) model (*47*) of neuropathic pain and the complete Freund’s adjuvant (CFA)-induced inflammatory pain model (*15*). Figure 3 illustrates that the 10 mg/kg ip dose of ST171 significantly increased the mechanical thresholds of the injured hindpaw, in both the inflammatory (Figure 3F) and neuropathic (Figure 3G) models. In fact, mechanical thresholds returned to baseline levels in the CFA, but not the SNI model. In contrast, the non-sedating dose of befiradol (1 mg/kg) was only antinociceptive in the CFA-induced pain model. Taken together, our results indicate that a systemic administration of our novel 5-HT_1A_R agonist has broad therapeutical potential across multiple modalities of acute and chronic pain and is clearly superior from a therapeutic window perspective, to befiradol.

### Structural studies on 5-HT_1A_R-G_i1_ complexes bound to ST171 or befiradol Overall structures

We determined cryo-EM structures of 5-HT_1A_R – G_i1_ protein complexes bound to ST171 and befiradol at nominal resolutions of 2.4 Å and 2.9 Å, respectively, (Figure 3) and performed unbiased MD simulations (in total 20 µs for each ligand-bound protein complex), mutagenesis and SAR studies. The cryo-EM maps enabled modelling of highly resolved receptor-G_i_ protein complexes and unambiguous placement of the ligands in the binding pocket (Figure 3 A-C and E-G). We were able to detect two cholesterols around the TM bundle in the ST171-bound structure and one cholesterol in the befiradol-bound structure (*48*). Compared to the inactive methiothepin-bound 5-HT_1B_R (PDB: 5V54) (*49*), our agonist-bound ternary complexes showed downward movement of the toggle switch residue W358^6.48^, movement of I124^3.40^ and an outward swing of F354^6.44^ in the PIF motif as well as a break of the DRY motif formed by R134^3.50^ and D133^3.49^ by a rotation of R134^3.50^. Additional structural changes include a rotation of the NPxxY motif at the cytoplasmic end of TM7 inducing an interaction of Y400^7.53^ with R134^3.50^ which is further stabilized by a phosphatidylinositol-4-phosphate (PI4P) (*50*). All these conformational changes contributed to an inward shift of TM7 by 3.3 Å (ST171) and 3.2 Å (befiradol) and an outward movement of TM6 by 9.3 Å (ST171) and 8.9 Å (befiradol) to facilitate G protein binding. In conclusion, the two structures show typical properties of active state G_i_-coupled GPCRs. The serotonin-bound 5-HT_4_R-G_s_ complex (PDB: 7XT9) (*51*) exhibits an outward shift of the extracellular TM2 by 1.4 Å, along with an inward shift of the intratracellular region of TM7 by 3.2 Å, and a significant kink of TM5 towards TM6 (Supplementary Figure 5).

Interestingly, an assignment of the location and orientation of the extracellular parts of the ST171-bound receptor compared to its befiradol-bound structure reveals a reduced TM2 and TM7 distance, a higher distance between TM1 and TM2 and a disposition of TM7 away from TM6 (Figure 3F). This rearrangement appears to be stabilized by an interhelical hydrogen bond between the amide group of Q97^2.65^ and the NH of W387^7.40^.

Compared to the LSD-bound 5-HT_2B_ receptor in complex with β-arrestin 1 (PDB: 7SRS) (*52*), our complexes exhibit an outward shift of TM6 and a shift of TM5 towards TM6. Additionally, the residues in the NPxxY motif of our complexes display a clockwise rotation when viewed from the extracellular side, in contrast to the β-arrestin 1 complex. Finally, the angle formed between TM7 and TM8 is smaller in our complexes compared to the β-arrestin 1 complex. These structural characteristics may serve as the basis for the functional selectivity of ST171.

### Binding modes and receptor-ligand interactions

We observed several differences in the binding poses of befiradol and ST171, which may explain their distinct pharmacological profiles. Both agonists show a bitopic binding mode occupying the conserved orthosteric pocket but addressing two different allosteric sites of the 5-HT_1A_R (Figure 4E).

The methoxyphenyl-substituted aminoethoxy group of ST171 and the methylpyridinyl-substituted aminomethyl moiety of befiradol occupy a similar space in the orthosteric binding pocket and mainly interact with V117^3.33^, I167^4.56^, I189^45.52^, A203^5.46^, F361^6.51^, F362^6.52^ and A365^6.55^ through hydrophobic interactions without engaging T121^3.37^, a residue that has been observed to interact with the indole NH of serotonin (Figure 4B) (*50*). In the vicinity of the ether group of the methoxyphenol ring of ST171, a water molecule was detected in the cryo-EM structure, which is likely involved in a water-mediated hydrogen bond with D116^3.23^. In accordance, an SAR study including a replacement of ST171’s ether function with a thioether unit, a weaker H-bond acceptor (*53*), showed a considerable reduction of potency and efficacy (compound TA48, Supplementary Figure 6, Supplementary Table 3) indicating a contribution of the methoxy group to the binding energy. One hydrophilic key interaction in this pocket is with the highly conserved D116^3.32^, to which the secondary amines of ST171 and befiradol form a charge-reinforced hydrogen bond, an interaction that is commonly seen in aminergic GPCRs including the serotonin-bound 5-HT_1A_R (*50*). Additionally, the cryo-EM structure revealed the presence of a water-mediated hydrogen bond between the secondary amine of ST171 and the backbone carbonyl of N386^7.39^. MD simulations were performed using the ligand-bound 5-HT_1A_R with a restrained G protein interface. The studies showed that water occupied the identified position in most frames, thereby indicating robust and highly conserved water-mediated hydrogen bonds. The distances between the secondary amine of befiradol or ST171 and D116^3.32^ displayed notable discrepancies, measuring 3.4 Å for ST171 and 4.0 Å for befiradol (distance between the secondary amine of the ligand and Cγ of D116^3.32^). This disparity is particularly intriguing, given that D116^3.32^ serves as a critical anchor for basic moieties of aminergic ligands (*54*). MD simulations substantiated the persistence of this distinction (average distances: 3.1 Å and 3.8 Å for ST171 and befiradol, respectively, measured between the secondary amine of the ligand and the center of mass of the carboxyl function of D116^3.32^) and showed higher fluctuation for the befiradol head group (Supplementary Figure 7). Insertion of a CH_2_ unit between the head group and the secondary amine of ST171 leads to an attenuation of binding and receptor activation (TA12, Supplementary Figure 6, Supplementary Table 3) confirming the importance of the charge-reinforced hydrogen bond.

Although both ligands overlap in the orthosteric binding pocket, befiradol and ST171 address different extended binding sites of the 5-HT_1A_R. The benzoxazinone moiety of ST171 reaches a cleft formed by TM2 and TM7 where it interacts with A93^2.61^ and Y96^2.64^ through hydrophobic interactions, with Q97^2.65^ and W387^7.40^ through polar interactions, and with N386^7.39^ through water-mediated interactions (Figure 4B, 3D and 3E). Notably, Y96^2.64^ adopts a conformation in which its phenyl moiety aligns nearly parallel to TM2, facilitating π-π interaction between the benzoxazinone ring and Y96^2.64^. This conformation deviates from those observed in all other available 5-HT_1A_R structures, where the plane of the phenyl moiety of Y96^2.64^ adopts a horizontal orientation. The stability of this unique conformation in the ST171 complex was corroborated by MD simulations, which indicated only minimal rotation of the phenyl moiety throughout the simulation. In contrast, simulations of befiradol- and serotonin-bound receptors exhibited a preferred conformation with an orthogonal orientation of the phenyl moiety of Y96^2.64^ with respect to TM2 (Supplementary Figure 8). We suggest that the π-π interaction observed in the ST171-bound complex is crucial for the 1.2 Å shift of the extracellular portion of TM2 relative to its orientation in the befiradol- bound complex. This shift, initially observed in the cryo-EM structure, persisted prominently throughout the MD simulations (Supplementary Figure 9). The binding pose of the benzoxazinone moiety that enables π-π stacking may stabilize an interhelical hydrogen bond that we observed between the amide group of Q97^2.65^ and the NH group of W387^7.40^ (Supplementary Figure 10). This hydrogen bond could not be observed in the befiradol-bound structure where the distance between the respective side chains was too high (4.3 Å). MD simulations of the befiradol-bound receptor indicate an equilibrium between a closed and an open state in the ternary complex model (restraint G protein interface) while the open state is more favored in the binary complex model (unrestraint G protein interface). In contrast, MD simulations using the ST171-bound receptor confirm a high stability of the closed state in both the ternary and the binary complex model. We propose that this interaction substantially contributes to the G_i_-biased signaling of ST171. SARs confirm the crucial role of the benzoxazinone moiety. Hence, modifying the position of the benzoxazinone by shortening or extending the linker unit results in a significant decrease in both potency and efficacy (TA13 and TA14, Supplementary Figure 6, Supplementary Table 3).

The allosteric appendage of befiradol, a 3-chloro-4-fluorobenzene moiety occupies a hydrophobic binding site between TM2, TM3 and ECL1 built from V89^2.57^, M92^2.60^, Y96^2.64^, W102^23.50^, F112^3.28^, I113^3.29^. This cavity is not open for ligand binding in the ST171-bound and the previous 5-HT_1A_R structures and becomes accessible for befiradol upon transmission into a conformer in which the sides chains of F112^3.28^ and M92^2.60^ move outward the TM bundle. Interestingly, spontaneous formation of this hydrophobic pocket could not be observed by MD simulations of both the ternary (restrained G protein interface) and the binary complex models (unrestrained G protein interface) of the non-liganded 5-HT_1A_ receptor. Starting from the befiradol- bound 5-HT_1A_R-G_i_ complex, the hydrophobic pocket closed within 10 ns by a conformational change of F112^3.28^ and M92^2.60^. This data suggests that the hydrophobic pocket is formed by an induced fit mechanism in which the 3-chloro-4-fluorobenzene moiety forces entry into the newly formed hydrophobic pocket rather than by conformational selection. This observation has been confirmed by metadynamics simulations indicating that an energy barrier of approximately 7.5 kcal/mol has to be overcome to facilitate the opening of the hydrophobic pocket (Supplementary Figure 11). Instead of F112^3.28^, this cavity is closed by a larger tryptophan residue in more than 70 % of the aminergic GPCRs. Mutation of F112^3.28^ to tryptophan resulted in a 16-fold decrease in the binding affinity of befiradol (5-HT_1A_R: *K*_i_ = 15 ± 3.4 nM; 5-HT_1A_R-F112^3.28^W: *K*_i_ = 240 ± 63 nM) whereas binding of serotonin and ST171 were hardly affected (serotonin: *K*_i_ [5-HT_1A_R] = 220 ± 52 nM; *K*_i_ [5-HT_1A_R-F112^3.28^W] = 140 ± 29 nM; ST171: *K*_i_ [5-HT_1A_R] = 0.41 ± 0.06 nM; *K*_i_ [5-HT_1A_R-F112^3.28^W] = 0.16 ± 0.05 nM). The effect of the residue substitution underlines the importance of the gate-keeper function of F112^3.28^ for the 5-HT_1A_R selectivity of befiradol.

## Discussion

Analysis of our in-house library of about 10,000 compounds identified the benzoxazinone ST171, which binds the 5-HT_1A_R with an affinity in the sub-nanomolar range. Functional and structural investigation of this ligand in complex with the 5-HT_1A_R led to some unanticipated, but yet highly interesting findings.

First, ST171 is a strong partial agonist for the activation of Gα_i/o_, causing inhibition of cAMP accumulation, but only minimal effect on the recruitment of β-arrestin 2. Intriguingly, 5-HT_1A_R activation by ST171 is functionally selective for the Gα_i/o_ family, and in particular for Gα_oA_ and Gα_z_. In crontrast the former drug candidate, befiradol, and the endogenous agonist serotonin, do induce accumulation of cAMP at higher ligand concentrations, suggesting alternative Gα_s_–coupling of the 5-HT_1A_R. Interestingly, preferred coupling to Gα_oA_ and Gα_z_ was also described for the promising analgesic agents, mitragynine pseudoindoxyl (MP) and ‘9087, which bind the μ-opiod or the α_2A_-adrenergic receptor, respectively (*15, 55*). This observation suggests that a preferred activation of Gα_oA_ and Gα_z_ may be of particular relevance to developing of both safe and effective pain management drugs.

Second, our *in vivo* behavioural analyses revealed that ST171 has broad therapeutical potential, both in acute and chronic pain mouse model. ST171 is also superior from a therapeutic window perspective to other 5-HT_1A_R biased agonists, such as befiradol (*56*). Targeting the 5-HT_1A_R for the treatment of pain has a strong therapeutic rationale, particularly via an act at the spinal cord, where its activation is antinociceptive (*40, 42, 57*). Problematic, of course, is that because of widespread expression of the receptor, generating highly selective 5-HT_1A_R agonists devoid of side effects has been challenging. Indeed, our analysis suggests that motor impairment contributed to befiradol’s potent antinociceptive effects at higher doses. At lower non-sedating doses, befiradol although still efficacious against heat pain, was ineffective at reversing the mechanical allodynia in a neuropathic pain model. This conclusion is consistent with previous studies showing poor efficacy of befiradol against the mechanical hypersensitivity that develops in diabetic or oxaliplatin-induced neuropathy (*38, 58*). In contrast, we showed that non sedating doses of ST171 retained strong analgesic effects across multiple modalities (heat and mechanical) of both inflammatory (CFA) and neuropathic (SNI) pain models. We conclude that selective and potent 5-HT_1A_R agonists to treat pain conditions with different etiologies and an excellent therapeutic window, can be developed.

Third, structural studies of the ligand-receptor complexes in the presence of Gα_i_ by cryo-EM reveal that ST171 and befiradol engage the 5-HT_1A_R in different ways, which likely accounts for the distinct pharmacology observed for the two ligands. While both ligands fill the orthosteric binding pocket similar to the endogenous ligand serotonin, they extend further into different exo-sites of the receptor. For befiradol we observe an induced-fit into a cleft between TMs 2, 3 and ECL1. This cavity is not available in any of the other 5-HT_1A_R structures available to date and may explain the high 5-HT_1A_R selectivity of befiradol. In contrast, the benzoxazinone moiety of ST171 extends towards TMs 2 and 7, anchoring TM2 in a more inward conformation. We propose that π-π interaction observed in the exo-site of the ST171-bound complex is crucial for a 1.2 Å shift of TM2 relative to its orientation in the befiradol-bound complex. The binding pose of the benzoxazinone moiety that enables π-π stacking confers ligand-specific stabilization of an interhelical hydrogen bond between the amide group of Q97^2.65^ and the NH group of W387^7.40^ which could not be observed in the befiradol-bound structure, where the distance between the respective side chains was too high (4.3 Å). These observations may explain how ST171 binding stabilizes the receptor in a conformation that exhibits high selectivity for G_i/o_ protein binding with preference to Gα_i/o_ while showing negligible affinity for G_s_ protein and minimal recruitment of β-arrestin 2. To learn how befiradol and serotonin stabilize ternary complexes of the 5-HT_1A_R with Gα_s_ and β-arrestin in detail, further structural studies will be necessary.

While ST171 has high affinity and subtype selectivity for the 5-HT_1A_R, it is not entirely selective, as it also engages α_1A_R and receptors of the dopamine D_4_ subtype affinity in the subnanomolar range. Future ligand optimization will enable the reduction of off-target effects and, thus, leverage the development of a safe and efficacious antinociceptive drug candidate.

## Contributions of the authors

A.U. and J.S. cloned, expressed and purified the 5-HT1AR-G_i1_ complexes with the cooperation of A.A. B.B., T.R., and T.H. prepared the grids, collected and processed the cryo-EM data. J.S. built the complex models and performed structure refinement assisted by A.U. A.U., H.H. and J.B. characterized the pharmacological properties by radioligand binding and functional *in vitro* assays with assistance from D.W. J.M.B., K.B and S.R-R carried out the *in vivo* experiments. M.S. and T.A. performed the chemical synthesis and analytics of the compounds under the supervision of S.L. N.S. and E.N. carried out and analyzed MD simulations. E.F. and B.K.S. coordinated pK studies. A.I. provided the knockout cell lines and contributed to the data interpretation. The manuscript was written by A.U., J.S., N.S., E.N., D.W, and P.G. with contributions from all authors. P.G., B.B., A.J.B. and D.W. coordinated the experiments. P.G. conceived the project and supervised the overall research.

## Acknowledgments

This project was supported by the DFG grant GRK 1910 (P.G, A.U., A.A.), the Elitenetwork of Bavaria (J.S.), the Verband der Chemischen Industrie (Kekulé fellowship, E.N.) and the Studienstiftung des Deutschen Volkes (E.N.). This work benefited from access to EMBL-IC generously supported by the Boehringer Ingelheim Foundation and has been supported by iNEXT-Discovery, project numbers 20311 and 24759, funded by the Horizon 2020 program of the European Commission. Electron Cryo Microscopy for sample evaluation was carried out in the cryo EM-facility of the Julius-Maximilians-Universität Würzburg funded by the Deutsche Forschungsgemeinschaft (DFG - 359471283, 456578072, 525040890. MD calculations were performed at the NHR cluster at FAU (grant: b132dc). We thank Simon Fromm and Christian Kraft for their technical support in obtaining the Cryo-EM structures. Michel Bouvier (Université de Montréal) generously provided the constructs for enhanced bystander BRET and the Gα_i1_-BRET assay. A.I. was funded KAKENHI JP21H04791, JP21H05113, JP21H05037 and JPJSBP120213501 from Japan by Society for the Promotion of Science (JSPS); JPMJFR215T, JPMJMS2023 and 22714181 from Japan Science and Technology Agency (JST); JP22ama121038 and JP22zf0127007 from the Japan Agency for Medical Research and Development (AMED).

## Materials and Methods

### Compounds

#### Chemical Synthesis

The synthesis routes towards ST162, ST171 and analogs thereof are depicted in the Supplementary Figures 14 and 15, respectively. A detailed description of the chemical synthesis and the analytical data are provided as Supplementary text.

#### Test compounds

Serotonin (catalog no.: H9523) and befiradol (catalog no.: SML2324) were purchased as hydrochlorides from Sigma-Aldrich (Steinheim, Germany). The Discovery Probe FDA-approved Drug Library (APExBio, 1971 compounds, 10 mM) was purchased from BioTrend, Cologne, Germany. Stock solutions for befiradol, ST171, its structural analogs and the in-house library were prepared in DMSO at a concentration of 10 mM. The stock solution of serotonin was prepared in water with a concentration of 10 mM. Dilutions were prepared in the buffers indicated for each assay.

#### Plasmids and mutagenesis

If not noted otherwise, the human 5-HT_1A_R and human isoforms of the other GPCR in pcDNA3.1 (cDNA.org) was used for in vitro experiments. The 5-HT_1A_R-F112^3.28^W mutant was generated by PCR employing the Quikchange method. Sequence identity was verified by DNA sequencing (Eurofins Genomcis).

### Radioligand binding

#### High-throughput screening with radioligand displacement

Starting point of the screening for new 5-HT_1A_R chemotypes was our in-house library of approximately 10,000 compounds consisting of bioactive compounds generated in our recent GPCR projects and a sub-library of 2,000 FDA-approved drugs for addressing targets beyond GPCRs. To create a screening library that comprises a broad variety of different chemotypes, we carefully inspected our in-house library of about 8,000 GPCR ligands to select compounds representing a set of great structural diversity. This sub-library comprised of 100 compounds of structurally diverse ligands. The second test library constituted of 1,971 samples of the DiscoveryProbe™ FDA-approved drug library (L1021-100, APExBIO, USA purchased from Biotrend, Cologne, Germany) consisting of FDA-approved drugs with known bioactivity and safety data in humans. These test libraries were supplied to an affinity screening for 5-HT_1A_R binding applying a radioligand displacement assay.

In detail, the library samples were provided as 10 mM stocks in DMSO and were diluted to a 10-fold working solution with binding buffer (50 mM TRIS, 5 mM MgCl_2_, 1 mM EDTA, 100 µg/mL bacitracin at pH 7.4). Membranes from HEK293T cells transiently transfected with den cDNA of the human 5-HT_1A_R (cDNA Center, Bloomsberg, PA) and expressing the receptor with density of B_max_ = 2300 fmol/mg protein and with a K_D_ = 0.070 nM were incubated at 3 µg protein per well with 0.2 nM of the radioligand [^3^H]WAY600,135 (specific activity: 83 Ci/mmol, Novandi, Södertäljie, Sweden) and the test compounds at a final concentration of 10 nM for 60 min at 37°C. Receptor bound radioactivity was isolated by filtration and subsequently measured in a scintillation counter as described (*59*). Radioligand displacement induced by each compound was determined in two independent single point measurements. Specific binding of the radioligand was calculated by normalization of the radioactive counts to total binding in the presence of buffer (100 %) and unspecific binding (0 %) evaluated in presence of 5 µM of unlabeled WAY600,135. Mean displacement of the radioligand as parameter for the binding affinity of the test compounds was calculated by applying the equation [displacement [%] = 100 – specific binding [%]].

Screening was continued with 60 analogs of ST162 involving the dose-dependent investigation of ligand binding and the determination of *K*_i_-values for 5-HT_1A_. ST171 was identified because of its superior binding affinity.

#### Radioligand binding

To determine the binding affinities of ST171 and the references serotonin and befiradol to the serotonin receptor subtypes 5-HT_1A_, 5-HT_2A_, and 5-HT_6_ dose-response curves were measured using membranes from HEK293T cells transiently transfected with the appropriate cDNA (all purchased from cDNA Center, Bloomsberg, PA) and the corresponding radioligand (Supplementary Table 5) as described previously (*5, 59*). In brief, membranes were incubated in binding buffer at final protein concentrations of 2-10 µg/well with radioligand (at a concentration similar to the *K*_D_) and varying concentrations of the competing ligands for 60 minutes at 37 °C. Nonspecific binding was determined in the presence of unlabeled ligand (for 5- HT_1A_, 5-HT_2A_) or serotonin (5-HT_6_) at 10 µM. Protein concentration was measured using the method of Lowry (*60*). The resulting competition curves were analyzed by nonlinear regression using the algorithms implemented in Prism 9.0 (GraphPad Software, San Diego, CA) to provide IC_50_ values, which were subsequently transformed into a K_i_ values applying the equation of Cheng and Prusoff (*61*). Mean *K*_i_ values in [nM±SD] were derived from 2-11 experiments each performed in triplicates. Binding affinities to GPCRs of the dopamine, adrenergic, muscarinergic, opioid, orexin, and neurotensin receptor family were evaluated similarly and according to the details listed in Supplementary Table 5.

#### Radioligand saturation

The evaluation of the receptor density in 5-HT_1A_R expressing HEK293T cells was performed with radioligand saturation experiments using whole cells. Cells were transfected as described for the functional experiments (see: *BRET-based Gα_i1_ activation and β-arrestin 2 recruitment*). After 24 h cells were split to be transferred to microplates for functional testing or maintaining cell growth in culture dishes for saturation binding experiments. Cells were harvested at the same time as for the functional tests and resuspended and diluted in binding buffer (see above) to get 30,000 cells per well. Radioligand saturation binding was performed by incubating the cells with [^3^H]WAY600,135 at a concentration in the range of 1-2 nM, which represents the saturation concentration for 5-HT_1A_R binding, and was worked up as described above. Receptor density was calculated by transforming the values for specific binding, number of cells and specific activity of the radioligand to B_max_ in [receptors/cell]. Mean values were derived from 4 to 8 single experiments each done in triplicates.

### *In vitro* functional studies

#### BRET-based cAMP assay (CAMYEL biosensor)

G_i/s_ Protein activation was determined by the detection of intracellular cAMP levels with the BRET based CAMYEL biosensor (*62*). CHO-K1 cells were grown in culture medium (DMEM/F12 medium supplemented with 10 % fetal bovine serum, 100 µg/mL penicillin, 100 µg/mL streptomycin, and 0.5 mg/mL L-glutamine) at 37 °C and 5 % CO_2_ and transiently transfected with 2 µg 5-HT_1A_R and 1.0 µg biosensor per 10 cm culture dish using Mirus TransIT-2020 (Peqlab, Erlangen, Germany) as transfection reagent. After 24 h, cells were transferred into white 96-well half area or 384-well plates (Greiner) in culture medium at a density of 15,000 cells / well (96 well) or 10,000 cells / well (384 well), and maintained at 37 °C and 5 % CO_2_ for further 24 h. The medium was exchanged with 20 µL DPBS in which cells were incubated for 30 min, before the buffer was exchanged with fresh DPBS followed by an additional incubation for 30 min at 37 °C. 10 µL coelenterazine h (Promega, 5 µM final concentration) was added. Following incubation for 15 min, ligand dilution row and forskolin (3 µM final concentration) were added. After additional 15 min, BRET ratio was determined from light emission at 475 and 530 nm with the Clariostar plate reader (BMG, Ortenberg, Germany) . All responses were normalized to serotonin and analyzed using the algorithms for nonlinear regression (4 parameters) in PRISM 6.0. For bell-shaped curves, two nonlinear regressions were fit from the global minimum to the global maximum and from the global maximum to the local minimum at high ligand concentrations. To investigate the solely G_s_ Protein mediated response, pertussis toxin (25 ng/mL, Sigma-Aldrich, Taufkirchen, Germany) was added during cell seeding on assay plates.

#### GloSensor cAMP assay

G_i_ and G_s_ activation by 5-HT_1A_R and the Gα_i_ subunit-coupling profile was investigated using the GloSensor cAMP assay and knock-out cell lines HEK293AΔG_s_, HEK293AΔG_i_, HEK293AΔARRB1/B2 (*17, 20, 21*) and HEK293T and HEK293A cells. Cells were cultured in culture medium (DMEM/F12 medium supplemented with 10 % fetal bovine serum, 100 µg/mL penicillin, 100 µg/mL streptomycin, and 0.5 mg/mL L-glutamine) at 37 °C and 5 % CO_2_ on 10 cm culture dishes. For HEK293AΔGi, dishes were coated with type I bovine collagen (Corning). Cells with confluency of 60-80 % (10 cm culture dish) were transfected with 0.5 µg (HEK293A, HEK293T, HEK293AΔARRB1/B2) or 1 µg 5-HT1_A_R plasmid DNA (HEK293AΔG_s_, HEK293AΔG_i_), 0.5 µg (HEK293A, HEK293T, HEK293AΔARRB1/B2) or 2 µg (HEK293AΔG_s_, HEKS293AΔG_i_) Glo-22F cAMP plasmid DNA (Promega) and 2 µg MOCK DNA (HEK293A, HEK293T, HEK293AΔARRB1/B2). For the investigation of the Gα_i_ subunit-coupling profile HEK293AΔG_i_ cells were co-transfected with 0.5 µg of the plasmid encoding the respectiveGα_i_ subunit (Gα_i1_, Gα_i2_, Gα_i3_, Gα_z_, or Gα_oA_, www.cdna.org). All transfections were carried out using using Mirus TransIT-293 as transfection reagent in a 3:1 reagent to DNA ratio. After incubation for 24 h, cells were detached and transferred into CP4 medium (Eurofins DiscoverX Products) at density of 15,000 cells/well (HEK293T) or 10,000 cells/well (HEK293A, HEK293AΔG_s_, HEK293AΔG_i_, HEK293AΔARRB1/B2) into white bottom 384-well plates (Greiner) coated with poly-D-lysine (Sigma-Aldrich, for HEK293T, HEK293A, HEK293AΔG_s_, HEK293AΔARRB1/B2) or type I bovine collagen (HEK293AΔG_i_). Plates were incubated at 37 °C and 5 % CO_2_ for further 24 h. After incubation, CP4 medium was exchanged with Hanks’Balanced Salt Solution (with added glucose 1 g/l, NaHCO_3_ 0.35 g/l) containing GloSensor cAMP Reagent (3 %, Promega) and plates were incubated for 60 min. Afterward, ligand dilutions were added, and plates were incubated for 15 min (HEK293T, HEK293A, HEK293AΔG_i_, with or without cotransfected Gα_i_ subunits, and HEK293AΔARRB1/B2) or 10 min (HEK293AΔG_s_) at room temperature in the dark. Forskolin was added (10 µM final concentration HEK293T, HEK293A, HEK293AΔARRB1/B2, and HEK293AΔG_i_ co-transfected with Gα_i_ subunits; 5 µM final concentration with additional phosphodiesterase inhibitor RO-20-1724, 100 µM final concentration HEK293AΔG_s_) and the plates were incubated for further 15 min (HEK293T, HEK293A, HEK293AΔARRB1/B2, and HEK293AΔG_i_ co-transfected with Gα_i_ subunits) or 20 min (HEK293AΔG_s_) at room temperature in the dark. Luminescence was determined with the Clariostar plate reader (BMG LabTech) or the luminescence was directly measured after incubation with ligand dilutions (HEK293AΔG_i_). All responses were normalized to the minimal and maximal effect of serotonin and analyzed using nonlinear regression in PRISM 6.0. For bell-shaped curves, two nonlinear regressions were fit from the global minimum to the global maximum and from the global maximum to the local minimum at high ligand concentrations.

#### IP-One accumulation

Determination of Gα_i_ and Gα_s_ selective signaling at the 5-HT_1A_R was performed applying an IP accumulation assay (IP-One HTRF^®^, PerkinElmer, Rodgau, Germany) according to the manufacturer’s protocol and in analogy to previously described protocols (*15, 63*). HEK293T cells were transiently co-transfected with the cDNA for 5-HT_1A_R and the hybrid G-protein Gα_qi_ or Gα_qs_ (Gα_q_ proteins with the last five amino acids at the C-terminus replaced by the corresponding sequence of Gα_i_ or Gα_s_) (gift from The J. David Gladstone Institutes, San Francisco, CA), respectively in a ratio of 1:2. On the next day cells were transferred to 384 well micro plates (Greiner, Frickenhausen, Germany) and incubated for further 24 h. On the day of the experiment test compounds with concentrations from 0.01 to 100 µM were added to the cells and incubated for 90 min. Accumulation of the second messenger was stopped by addition of the detection reagents (IP1-d2 conjugate and Anti-IP1cryptate TB conjugate) for 60 min. To measure Gα_q_ mediated signaling of the adrenergic α_1A_ and α_1B_ receptor HEK293T were transfected with the cDNA for human α_1A_ and α_1B_ receptor (cDNA Center, Bloomsberg, PA) and worked up as described above. TR-FRET was monitored with a Clariostar plate reader. FRET-ratios were calculated as the ratio of emission intensity of the FRET acceptor (665/10 nm) divided by the FRET donor intensity (620/10 nm). Raw FRET-ratios were normalized to buffer conditions (0%) and the maximum effect of norepinephrine or serotonin (100%). Data analysis was performed using the equation for sigmoid concentration-response curves (four-parameter) implemented in GraphPad Prism 9.3 (GraphPad Software, La Jolla, USA) to derive the maximum effect (E_max_) and the ligand potency (EC_50_). Three to eight repeats in duplicate were performed for each test compound.

#### BRET-based Gα_i1_ activation and β-arrestin 2 recruitment

For the determination of G-protein signaling relative to receptor density of 5-HT_1A_R a BRET biosensor based assay using Gα_i1-RLucII_ together with Gβ_1_ and Gγ_2-GFP10_ (*15, 64*) was performed. HEK293T cells were co-transfected with 400 ng, 200 ng, 100 ng, 40 ng, or 10 ng of cDNA for 5-HT_1A_R and plasmids for G protein activation (100 ng Gα_i1_, 200 ng Gβ_1_, 800 ng Gγ_2_) using the Mirus TransIT-293 reagent (Peqlab). DNA was complemented to a total amount of 2 µg DNA per dish with ssDNA (Sigma Aldrich). After 24 h cells were transferred in 96 well plates (Greiner) and grown for further 24 h. For the experiment the cell medium was exchanged with PBS (phosphate buffered saline) and cells were stimulated with ligands at 37°C for 10 min. Coelenterazine 400a (abcr GmbH, Karlsruhe, Germany) at a final concentration of 2.5 µM was added 5 min before measurement. Gα_i_ mediated signaling by the adrenergic α_2A_, α_2B_, and α_2C_ receptors as well as by the dopamine D_4_ receptor was monitored with cells co-transfected with 200 µg of the appropriate receptor DNA (α_2A_ was provided by Yang Du, University of Hong Kong, Shenzhen, CA; α_2B_, α_2C_, and D_4_ were purchased from cDNA Center, Bloomsberg, PA) and the plasmids for G protein activation (the ratio of receptor/Gα/Gβ/Gγ was 4/1/2/8) using linear polyethyleneimine (PEI, Polysciences, 3:1 PEI:DNA ratio) (*65*) according to the assay protocol described above. 5-HT_1A_R mediated β-arrestin 2 recruitment was measured applying the enhanced bystander BRET when co-transfecting 100 ng of 5-HT_1A_R together with 20 ng of β-arrestin_RLucII_, 300 ng of the bystander protein CAAX_rGFP_ and 100 ng of GRK2 with linear polyethyleneimine in HEK293T cells (*15, 22*). The assay conditions were identical to the protocol for monitoring Gα_i_ signaling. BRET was monitored on a Clariostar plate reader with the appropriate filter sets (donor 410/80 nm, acceptor 515/30 nm) and was calculated as the ratio of acceptor emission to donor emission. BRET ratio was normalized to the effect of buffer (0%) and the maximum effect (100%) of serotonin for 5-HT_1A_R, norepinephrine for the α_2_ adrenoceptors, and quinpirole for D_4_. For each compound 4-17 individual experiments were performed when investigating receptor density dependent 5-HT_1A_R activation, 3-14 for α_2_AR activation, 7-8 repeats for D_4_R activation, and 5-6 experiments when measuring arrestin recruitment by 5-HT_1A_R stimulation, each done in duplicates.

#### β-arrestin 2 recruitment assay with the PathHunter assay

For the enzyme fragment complementation-based arrestin recruitment assay, the ARMS2-PK2 sequence was C-terminally fused to the human wild-type 5-HT_1A_R using the Gibson-assembly method and a NheI-HF and HindIII-HF digested pCMV-ARMS2-PK2 vector (Eurofins DiscoverX Products). Sequence identity was confirmed using DNA sequencing (Eurofins Genomics). HEK 293 β-arrestin-2-EA cells were cultured in culture medium (DMEM/F12 medium supplemented with 10 % fetal bovine serum, 100 µg/mL penicillin, 100 µg/mL streptomycin, 0.5 mg/mL L-glutamine, and 150 µg/mL hygromycin B) at 37 °C and 5 % CO_2_ and transiently transfected with 5-HT_1A_R-ARMS2-PK2 (2 µg/10 cm cell culture plate) using Mirus TransIT-293. After 24 h, cells were transferred into white 384-well plates (Greiner) in CP4 assay medium at a density of 5,000 cells / well, and maintained at 37 °C and 5 % CO_2_ for further 24 h. On the day of the assay, 5 µL/well of the compounds diluted in PBS were added and plates were incubated at 37 °C for 180 min. After adding 10 µL/well detection mix (3.5 % substrate reagent 2, 18 % substrate reagent 1, and 78.5 % cell assay buffer), plates were incubated at room temperature in the dark for one hour. Chemiluminescence was detected with the CLARIOstar plate reader (BMG LabTech). All responses were normalized to the minimal and maximal effect of serotonin and analyzed using the algorithms for nonlinear regression in PRISM 6.0.

#### *In vivo* behavioral studies

Animal experiments were approved by the UCSF Institutional Animal Care and Use Committee and were conducted in accordance with the NIH Guide for the Care and Use of Laboratory animals (protocol #AN195657). Adult (8-10 weeks old) male C56BL/6 mice (strain # 664) were purchased from the Jackson Laboratory. Mice were housed in cages on a standard 12:12 hour light/dark cycle with food and water ad libitum. Sample sizes were modelled on our previous studies and on studies using a similar approach, which were able to detect significant changes. For all behavioral tests, the experimenter was always blind to treatment. Animals were habituated for 60 min in Plexiglas cylinders and then tested 30 min after intraperitoneal (IP) injection of the compounds. ST171 was dissolved in 2HPβCD-saline (1%:99%) and Befiradol in Kolliphor HS15-saline-water (20%:40%:40%). For 10 mg/kg doses, compounds were prepared in 2.5 mg/ml solutions of which 100 µL were administered to mice (average weight 25 g). Hindpaw mechanical thresholds were determined with von Frey filaments using the up-down method (*66*). Hindpaw thermal sensitivity was measured with a radiant heat source (Hargreaves) or by placing the mice on a 52°C hotplate. For the tail flick assay, sensitivity was measured by immersing the tail into a 50°C water bath. For the ambulatory (rotarod) test, mice were first trained on an accelerating rotating rod, 3 times for 5 minutes, before testing with any compound. On the test day, latency to fall from the rod was measured 30 min after injection of the compound. The cutoff was 300 s. Inflammation-induced hyperalgesia was generated by a unilateral intraplantar injection of complete Freund’s adjuvant (CFA; 20 μL of of 50% solution in saline; Sigma) and tested 3 days post-CFA. Peripheral nerve injury-induced allodynia was generated by ligating and transecting two of the three branches of the sciatic nerve, leaving the sural nerve intact (*47*). Hypersensitivity was tested 7 days after injury. Statistical tests were performed with GraphPad Prism 9.0 (GraphPad Software Inc., San Diego).

### Sample preparation for cryo-EM and structure determination

#### Protein engineering

DNA coding for the human 5-HT_1A_R with an N-terminal hemagglutinin signal sequence (HA), a FLAG tag (DYKDDDDA), an AAA-linker, followed by a 10 × histidine tag, and a tobacco etch virus protease cleavage site was codon optimized for expression in insect cells and synthesized by Eurofins Genomics Germany. A L125W^3.41^ point mutation was introduced to increase the expression and stability of the protein. A dominant-negative human Gα_i1_ (*67*) (dnGα_i1_ with S47N, G203A, E245A, A326S) was used to improve the stability of the nucleotide free G protein, which stabilizes the active state of the receptor.

#### Expression and purification of the 5-HT_1A_R – G Protein complex

Recombinant baculoviruses encoding the 5-HT_1A_R construct, dnG_αi1_, human 8×HIS-GSSG-G_β1_, and human G_γ2_ were generated with the Bac-to-Bac^TM^ baculovirus expression system (Invitrogen). *Spodoptera frugiperda* Sf9 cells (Invitrogen) at a density of ∼ 4 × 10^6^ cells / mL were coinfected with all complex components (receptor construct 1:100, each G protein subunit 1:200 virus to cell culture volume) in the presence of 1 µM ST171 or 1 µM befiradol, respectively. To increase the receptor expression 32 mL / L cell culture saturated cholesterol in 5 % methyl-β-cyclodextrin was added after 24 h. Cells were harvested by centrifugation 48 h post infection and stored at -80 °C. If not mentioned otherwise, the following steps were performed at 4 °C. Cell pellets from 2 L cell culture were thawed in hypotonic buffer (20 mM HEPES, 10 mM MgCl_2_, 20 mM KCl, 10 µM ST171 or befiradol, supplemented with a protease inhibitor cocktail (Roche)). After homogenization with a potter, cells were centrifuged for 20 min at 25,000 g. Cell pellets were resuspended in hypertonic buffer (20 mM HEPES, 10 mM MgCl_2_, 20 mM KCl, 100 mM NaCl, 5 mM CaCl_2_, 10 µM ST171 or befiradol, supplemented with a protease inhibitor cocktail (Roche)) and incubated with 25 mU / mL apyrase for 1 h at RT. The complex was solubilized with 0.5 % lauryl-maltose-neopentyl- glycol (LMNG, Anatrace) and 0.01 % cholesteryl hemisuccinat (CHS, Anatrace) for 2 h. Unsolubilized material was removed by centrifugation for 20 min at 25,000 g, before the complex was immobilized by on column binding to an Anti-FLAG antibody packed resin (Sigma). After washing the resin with 9 column volumes wash buffer (20 mM HEPES, 5 mM MgCl_2_, 100 mM NaCl, 0.01 % LMNG, 0.0025 % CHS, 10 µM ST171 or befiradol), the complex was eluted with FLAG peptide (200 µg / mL) in wash buffer. The complex was further purified by size-exclusion chromatography at RT using an Äkta go FPLC with a Superdex 200 Increase 10 / 300 GL column (Cytiva) in SEC buffer (20 mM HEPES, 100 mM NaCl, 0.002 % LMNG, 0.0005 % CHS, 10 µM ST171 or befiradol). Eluted fractions were analyzed by SDS-PAGE. Fractions containing the complex were pooled and concentrated with a VivaSpin 2 (Cytiva, MWCO = 100 kDa), before the sample was aliquoted and flash frozen in liquid nitrogen for storage at -80 °C.

#### Cryo-EM sample preparation

Quantifoil grids (UltraAuFoil R 0.6/1, Au 300 mesh, Quantifoil MicroTools, Jena, Germany) were treated by plasma cleaning (Harrick Plasma, model PDC-002,Ithaca, NY, USA) at an air pressure of 29 Pa for 2 min at medium-power. For ST171, plasma cleaned grids were washed with sample buffer to enhance the distribution of particle orientations. The samples were vitrified with a Vitrobot Mark IV (FEI): the chamber of the vitrobot was equilibrated to a temperature of 4 °C with 100% humidity and 3.5 µl of the sample (2- 6 mg/mL) were applied to the plasma-cleaned grid mounted in the chamber. This was followed by blotting off excess liquid for 5 s with an uncalibrated blot force of -20 before plunging the grid into liquid ethane. For each sample, batches with 4-6 grids were vitrified.

#### Cryo-EM imaging and Image processing

1-2 grids from each batch were evaluated in the cryo EM-facility of the Julius-Maximilians-Universität Würzburg (Titan-Krios G3, X-FEG, 300 kV, Falcon III direct electron detector): Movies were acquired with EPU at a magnification of 75k× (pixel size at specimen level 1.06 Å) in counting mode. The total exposure was 80 e^-^/Å² fractionated over 40 frames (exposure time 80 s). The movie-frames were motion-corrected, dose-weighted and averaged with MotionCor2 (*68*) as part of the pre-processing during the data acquisition. The averages were imported to CryoSPARC v3.5-4.1 (*69*) and further processed to obtain 2D- class averages, 3D-maps and variability components. The samples were selected for full-data acquisition based on particle density, 2D-class averages showing secondary structure elements and low heterogeneity in the variability analysis. The full data set was obtained from a different grid of the same batch with a Titan Krios3 G4 with C-FEG, Selectris X energy filter and Falcon IVi direct electron detector at the EMBL Imaging Center in Heidelberg (Germany): Movies were recorded in counting mode at a magnification of 165kx (pixel size at specimen level 0.73 Å) with zero-loss imaging (energy filter slit width=10 eV) and a total exposure of 80 e^-^/Å² (exposure time 3.7 s). Semi-automated data acquisition was done with SerialEM with 1 movie per hole and up to 9 holes per stage position with a targeted underfocus between 0.6 µm and 1.6 µm. The movies were recorded in EER-format and were motion corrected, dose-weighted and averaged with MotionCor2 parallel to the image acquisition with the “relion_it.py” script of Relion 4 (*70*). The movie averages were imported to cryoSparc 4.0-4.1 for further processing as outlined in the Supplementary Figures 12, 13. The final maps were auto-sharpened in Phenix (*71*) for subsequent model building and structural refinement.

#### Structure determination and refinement

The initial model (PDB 7E2Z) (*50*) was rigid body fit into cryo-EM maps using the Chimera (*72*) software. The 5-HT_1A_R – ST171 model was manually adjusted in COOT (*73*) and paired with iterative real-space refinement in Phenix (*74*). Model quality was evaluated using MolProbity. Figures of the structure and model were prepared in Chimera (*72*) and Pymol (https://pymol.org/2/).

### Molecular Dynamics Simulations

The simulations are based on the here reported cryo-EM structures of the 5-HT_1A_R in complex with ST171 and befiradol or based on the 5-HT_1A_R in complex with serotonin (PDB: 7E2Y) (*50*). Coordinates were prepared by removing the Gα/β/γ subunits of the heterotrimeric G protein. The cholesterol between TM1 and TM7 was retained for all complexes since previous studies indicated its effect inf the regulations of serotonin receptors (*50*). Due to the importance of water-mediated interactions in the binding mode of ST171 waters determine in the cryo-EM structure of ST171-bound complex were retained in the preparation of the ST171-bound system. Missing residues in the extracellular loop 2 (ECL2) were modeled using MODELLER (*75*). The long and flexible intracellular loop 3 (ICL3) was modeled in the binary complex using the first 5 amino acid residues after TM5 and the first 5 amino acid residues before TM6. For the ternary complex ICL3 was not modeled. The open termini, as well as the terminal residues at the intracellular ends of TM5 and 6 for the ternary complexes, were capped with an acetyl group or N-methylamid respectively. Using the h++ web-server (*76*) and manual inspection, all titratable residues were left in their dominant protonation state at pH 7.0 except D82^2.50^. Prior investigations have indicated that this residue is protonated in the active state (*77*). Parameter topology and coordinate files were generated using the tleap module of the AMBER22 program package (*78*). The created GPCR models were energy minimized using the PMEMD module of AMBER22 by applying 500 steps of steepest decent followed by 4500 steps of conjugated gradient. All files were converted to GROMACS input files using ParmEd (*79*). The protein structures were then aligned to the Orientation of Proteins in Membranes (OPM) (*80*) structure of the 5-HT_1A_R in complex with serotonin (PDB: 7E2Y). Each complex was inserted into a solvated and pre-equilibrated membrane of dioleoyl-phosphatidylcholine (DOPC) lipids using the GROMACS tool g_membed (*81*). Water molecules were replaced by sodium and chloride ions to result in neutral and physiological systems with 0.15 M NaCl. Final dimensions of the simulation systems were about 80 × 80 × 100 Å^3^ containing ∼65,000 atoms, including ∼154 DOPC molecules, ∼13,200 waters, ∼58 sodium, and ∼70 chloride ions. For all simulations the general AMBER force field (GAFF2) (*82*) was used for the ligands, the lipid 14 force field (*83*) for the DOPC and cholesterol molecules and ff14SB (*84*) for the protein residues. The SPC/E water model (*85*) was applied. Parameters for befiradol, ST171 and serotonin were assigned using antechamber (*78*). The structures of all ligands were optimized by Gaussian 16 using the B3LYP functional and the 6-31G(d) basis set. Charges were calculated using the HF functional and the 6-31G(d) basis set. Subsequently atom point charges were assigned according to the RESP procedure (*86*). A formal charge of +1 was defined for all ligands. Simulation were performed using GROMACS 2023.0 (*87*). Each simulation system was energy minimized and equilibrated in the NVT ensemble at 310 K for 1 ns followed by the NPT ensemble at 310 K and 1 atm for 1 ns with harmonic restraints of 10.0 kcal·mol^-1^ on the protein and ligands. In the NVT ensemble, the velocity-rescale thermostat was used. In the NPT ensemble the Berendsen barostat, the velocity-rescale thermostat, a surface tension of 22 dyn·cm^-1^, and a compressibility of 4.5 × 10^-5^ bar^-1^ was applied. Subsequently, the system was further equilibrated in the NPT ensemble at 310 K and 1 atm for 25 ns with restraints on protein backbone and ligand atoms in which the restraints were reduced every 5 ns in a stepwise fashion to be 10.0, 5.0, 1.0, 0.5 and 0.1 kcal·mol^-1^, respectively. Production simulations were performed for a duration of 2 µs. Five replicas were initiated for each ligand in both the binary and ternary complexes. Bond lengths to hydrogen atoms were constrained using LINCS (*88*) algorithm. Periodic boundary conditions were applied. A cutoff of 12.0 Å was used for Lennard-Jones interactions and the short-range electrostatic interactions. Long-range electrostatics were computed using the particle mesh Ewald (PME) (*89*) method with a fourth-order interpolation scheme and fast Fourier transform (FFT) grid spacing of 1.6 Å. A continuum model correction for energy and pressure was applied to long-range van der Waals interactions. The equations of motion were integrated with a time step of 2 fs. During production simulation of the ternary complexes, all residues within 5 Å of the G protein interface were restrained to the initial structure using 5.0 kcal·mol^-1^·Å^-2^ harmonic restraints applied to non-hydrogen atoms. Using such restraints instead of the intracellular binding partner reduces the overall system size, enabling faster simulations, while ensuring that the receptor maintains a conformation corresponding to the ternary complex throughout the simulation.

Multiple walker metadynamics simulations were conducted using the GROMACS 2021.4 software patched with PLUMED (*90*). Starting frames for the 32 individual walkers were taken from the trajectories of three independent 2 µs runs of unbiased MD simulations of the befiradol-bound binary complex model after dismission of its ligand. The dihedral angle between the N, C_α_, C_β_ and the C_γ_ atoms of the F112^3.28^ side chain was selected as a collective variable and was subject to a bias potential which enabled a reconstruction of the free energy surface. Gaussian hills with a height of 0.239 kcal·mol^-1^ were applied every 1.0 ps. Hill width was set to 0.1 rad. Rescaling of the gaussian function was done with a bias factor of 5.

Analysis of the trajectories was performed using Visual Molecular Dynamics (VMD) (*91*) and CPPTRAJ (*92*). Plots were created using Matplotlib 3.6.3 (*93*) and Seaborn 0.12 (*94*).

### Data and materials availability

The data of these findings are available in the main text and the supplementary materials. The structure model of ST171-5-HT_1A_R and befiradol-5-HT_1A_R ares deposited in the Protein Data Bank (PDB) under the accession codes PDB: 8PJK (ST171), and 8PKM (befiradol) and the 3D cryo-EM density maps in the Electron Microscopy Data Bank (EMDB) with the accession codes EMD-17712 (ST171) and EMD-17747 (befiradol), respectively.

## Supplementary text

**Chemical synthesis of ST162, ST171 and analogs thereof** (also see Supplementary Figures 14 and 15).

### General

Reagents and dry solvents were purchased in the highest available purity grade and used without further purification. MS was run on a BRUKER ESQUIRE 2000 ion-trap mass spectrometer or on a BRUKER amaZon SL mass spectrometer using ESI ionization source. HR-MS was performed on a Bruker Daltonic microTOF II focus TOF-MS spectrometer using ESI as ionization source. NMR spectra were obtained on a Bruker Avance 360 or a Bruker Avance 600 spectrometer at 300 K. ^1^H and ^13^C chemical shifts are given in ppm (δ) relative to TMS in the solvents indicated. IR spectra were performed on a Jasco FT/IR 4100 spectrometer using a film of substance on a NaCl plate. Purification by flash column chromatography was conducted using silica gel 60 (40-63 µm mesh) from Merck, Germany and eluents as binary mixtures with the volume ratios indicated. Purification with preparative HPLC were conducted on an Agilent 1100 preparative series HPLC system combined with a VWD detector. As HPLC column, a MACHEREY-NAGEL Varioprep VP 250/10 Nucleodur C18 HTec (10 x 250 mm, 5 µm) was used (flow rate 4 mL/min, λ = 254 nm). TLC analyses were performed using Merck 60 F_254_ aluminium sheets and analyzed by UV light (254 nm). Additionally, ninhydrin spray reagent was used for detection of aminergic compounds. Analytical HPLC was performed on an AGILENT 1200 series HPLC system employing a DAD detector. As HPLC column, a ZORBAX ECLIPSE XDB-C8 (4.6 x 150 mm, 5 µm) was used. HPLC purity was measured using the following binary solvent systems:

System 1: MeOH/H2O + 0.1 % HCOOH, flow 0.5 ml/min; gradient: 10 % MeOH for 3 min, 10 % - 100 % MeOH in 15 min, 100 % MeOH for 6 min, 100 % - 10 % MeOH in 3 min, 10 % MeOH for 3 min

System 2: ACN/H2O + 0.1 % TFA, flow 0.5 ml/min; gradient: 10 % ACN for 3 min, 10 % - 90 % ACN in 15 min, 90 % ACN for 6 min, 90 % - 10 % ACN in 3 min, 10 % ACN for 3 min The purity of all test compounds was determined to be >96%.

### 4-Methoxybenzene-1,2-diamine (**2**)

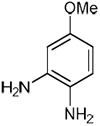

A suspension of 4-methoxy-2-nitroaniline (1.00 g, 5.95 mmol) and palladium on charcoal 10% wt. (100 mg) in methanol (10 mL) was stirred for 24 h under hydrogen atmosphere with light protection at room temperature. The suspension was filtered through celite and the solvent was evaporated. The crude product was purified by flash chromatography (CH_2_Cl_2_/methanol 99:1 + 0.2% NH_3_) to give **2** as brown solid (645 mg, 78% yield). ESI *m/z* 138.9 [M+1]^+^; IR (NaCl) ν (cm^-1^): 3338, 3288, 2954, 2925, 1628, 1515, 1451, 1295, 1248, 1202, 1166, 1032, 955; ^1^H-NMR (CDCl_3_, 600 MHz) δ (ppm): 3.24 (s(b), 4H), 3.73 (s, 3H), 6.26 (dd, *J* = 8.4, 2.8 Hz, 1H), 6.32 (d, *J* = 2.8 Hz, 1H), 6.64 (d, *J* = 8.4 Hz, 1H); ^13^C-NMR (CDCl_3_, 360 MHz) δ (ppm): 55.53, 102.93, 104.05, 118.33, 127.3, 137.07, 154.6.

### 5-Methoxy-1,3-dihydro-2H-benzo[d]imidazol-2-one (**3**)

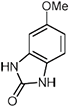

A solution of **2** (400 mg, 2.89 mmol) and 1,1’-carbonyldiimidazole (516 mg, 3.18 mmol) in DMF (11.4 mL) was stirred at 80 °C for 3 h. After the reaction has cooled to room temperature, the solution was concentrated in vacuo, 1 N HCl solution was added and the aqueous phase was extracted with ethyl acetate. The combined organic layers were washed with brine, dried over Na_2_SO_4_ and evaporated. The crude product was purified by flash chromatography (CH_2_CL_2_/methanol 98:2) to give **3** as a light brown solid (325 mg, 68% yield). ESI-MS *m/z* 164.9 [M+1]^+^; IR (NaCl) ν (cm^-1^): 3397, 3005, 2863, 1759, 1616, 1506, 1366, 1195, 1156, 1106, 1022, 887, 776; ^1^H-NMR (DMSO-*d_6_*, 600 MHz) δ (ppm): 3.69 (s, 3H), 6.49 – 6.52 (m, 2H) 6.8 (d, *J* = 8 Hz, 1H), 10.36 (s, 1H), 10.5 (s, 1H); ^13^C-NMR (DMSO-*d_6_*, 600 MHz) δ (ppm): 55.39, 95.25, 106.04, 108.67, 123.54, 130.48, 154.31, 155.61.

### 5-Hydroxy-1,3-dihydro-2H-benzo[d]imidazol-2-one (**4**)

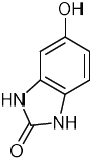

A solution of **3** (211 mg, 1.29 mmol) in CH_2_Cl_2_ (5.3 mL) was cooled to – 78 °C and put under argon atmosphere. 1 M BBr_3_ solution in CH_2_Cl_2_ (3.85 mL, 3.85 mmol) was added dropwise over 20 minutes. The mixture was kept at – 78 °C for 1 h, then allowed to warm to room temperature and stirred for another 2 h. The reaction was quenched by slowly adding satured NaHCO_3_ solution at -10 °C and pH was kept < 7. The mixture was extracted with ethyl acetate. The combined organic layers were washed with brine, dried over Na_2_SO_4_ and evaporated to give **4** as a reddish solid (113 mg, 59% yield). ESI-MS *m/z* 150.9 [M+1]^+^; IR (NaCl) ν (cm^-1^): 3353, 3096, 2998, 1727, 1598, 1479, 1366, 1241, 1023, 883, 815, 760, 611; ^1^H-NMR (DMSO-*d_6_*, 600 MHz) δ (ppm): 6.32 (dd, *J* = 8.3, 2.3 Hz, 1H), 6.37 (d, *J* = 2.3 Hz, 1H), 6.67 (d, *J* = 8.3 Hz, 1H), 8.85 (s, 1H), 10.19 (s, 1H), 10.30 (s, 1H); ^13^C-NMR (DMSO-*d_6_*, 600 MHz) δ (ppm): 96.39, 107.07, 108.67, 122.21, 130.42, 151.92, 155.55. *5-(3-Bromopropoxy)-1,3-dihydro-2H-benzo[d]imidazol-2-one (**5**):*

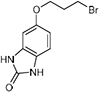

To a suspension of **4** (106 mg, 0.71 mmol) and anhydrous K_2_CO_3_ (117 mg, 0.85 mmol) in ethanol (1 mL) was added 1,3-dibromopropane (0.18 mL, 1.77 mmol) dropwise and the resulting mixture was refluxed for 3.5 h. After the reaction has cooled to room temperature, water was added and the aqueous phase was extracted with ethyl acetate. The combined organic layers were washed with brine, dried over Na_2_SO_4_ and evaporated. The crude product was purified by flash chromatography (CH_2_Cl_2_/methanol 96:4 + 0.2% NH_3_) to give 5 as a light yellow solid (57 mg, 30% yield). ESI-MS *m/z* 271.2 [M+1]^+^; IR (NaCl) ν (cm^-1^): 3434, 3235, 3024, 2925, 1754, 1708, 1642, 1510, 1468, 1254, 1170, 1025, 767; ^1^H-NMR (DMSO-*d_6_*, 600 MHz) δ (ppm): 2.18 – 2.23 (m, 2H), 3.66 (t, *J* = 6.6 Hz, 2H), 4.01 (t, *J* = 6 Hz, 2H), 6.51 – 6.55 (m, 2H), 6.79 (d, *J* = 8.4 Hz, 1H), 10.36 (s, 1H), 10.49 (s, 1H); ^13^C-NMR (DMSO-*d_6_*, 600 MHz) δ (ppm): 31.38, 31.93, 65.93, 96.16, 106.92, 108.68, 123.8, 130.45, 153.32, 155.58.

### 5-(3-((2-(2-Methoxyphenoxy)ethyl)amino)propoxy)-1,3-dihydro-2H-benzo[d]imidazol-2-one x TFA (**ST162**)

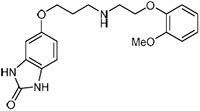

A solution of **5** (44 mg, 0.16 mmol), 2-methoxyphenoxyethylamine (*96*) (69 mg, 0.41 mmol) and KI (27 mg, 0.16 mmol) in CH_3_CN (2 mL) was refluxed under a nitrogen atmosphere for 4.5 h. After the reaction has cooled to room temperature, water was added and the aqueous phase was extracted with CH_2_Cl_2_. The combined organic layers were washed with brine, dried over Na_2_SO_4_ and evaporated. The crude product was purified by preparative HPLC (10-70% CH_3_CN/ 0.1% aq. trifluoroacetic acid) to give **ST 162** as a white solid (24 mg, 31% yield). ESI-MS *m/z* 358.5 [M+1]^+^; HR-MS calculated: 358.17613, found: 358.17578; IR (NaCl) ν (cm^-1^): 3411, 2962, 2921, 2848, 1740, 1636, 1503, 1369, 1256, 1205, 1027, 809, 613; ^1^H-NMR (DMSO-*d_6_*, 360 MHz) δ (ppm): 2.04 – 2.14 (m, 2H), 3.15 – 3.27 (m, 2H), 3.76 (s, 3H, OCH_3_), 4.01 (t, *J* = 6 Hz, 2H), 4.24 (t, *J* = 4.8 Hz, 2H), 6.49 – 6.57 (m, 2H), 6.81 (d, *J* = 8.3 Hz, 1H) 6.86 – 7.06 (m, 4H), 8.99 (s(b), 2H), 10.39 (s, 1H), 10.54 (s, 1H); ^13^C-NMR (DMSO-*d_6_*, 360 MHz) δ (ppm): 25.62, 44.96, 46.1, 55.46, 65.02, 65.4, 96.14, 106.89, 108.71, 112.38, 115.03, 120.73, 122.36, 123.83, 130.47, 147.07, 149.48, 153.26, 155.65; HPLC System 1: t_R_ = 14.9 min, purity: 99% (UV detection at λ = 254 nm), System 2: t_R_ = 9.7 min, purity: 99% (UV detection at λ = 254 nm).

### 6- Hydroxy-2H-benzo[b][1,4]oxazin-3(4H)-one (**7**)

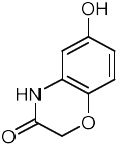

To a refluxing solution of 2*H*-benzo[*b*][1,4]oxazin-3(4*H*)-one (200 mg, 1.34 mmol) in trifluoroacetic acid (3.6 mL) was added a solution of [bis(trifluoroacetoxy)iodo]benzene (692 mg, 1.61 mmol) in trifluoroacetic acid (7.2 mL) via a syringe at once and the mixture was stirred for 10 min. After the reaction has cooled to room temperature, it was transferred slowly in an ice cold saturated NaHCO_3_ solution. The pH was adjusted to 4 – 5 with 1 N HCl and the aqueous phase was extracted with ethyl acetate. Combined organic layers were washed with brine, dried over Na_2_SO_4_ and evaporated. The crude product was purified by flash chromatography (hexane/ethyl acetate 4:1) to give 7 as an off white solid (100 mg, 45% yield). ESI-MS *m/z* 165.8 [M+1]^+^; IR (NaCl) ν (cm^-1^): 3152, 3055, 2915, 1683, 1620, 1498, 1416, 1309, 1203, 1047, 847, 818, 725; ^1^H-NMR (DMSO-*d_6_*, 600 MHz) δ (ppm): 4.42 (s, 2H), 6.27 (dd, *J* = 8.7, 2.8 Hz, 1H), 6.36 (d, *J* = 2.8 Hz, 1H), 6.72 (d, *J* = 8.7 Hz, 1H), 9.13 (s, 1H, benzo[*b*]oxazinone-OH), 10.52 (s, 1H).

### 6-(2-Bromoethoxy)-2H-benzo[b][1,4]oxazin-3(4H)-one (**8**)

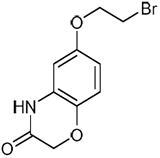

Compound **8** was synthesized according to the protocol of 5 by adding 1,2-dibromoethane (90 µl, 1.0 mmol) to a mixture of 7 (68 mg, 0.41 mmol) and K_2_CO_3_ (57 mg, 0.41 mmol) in ethanol (5 mL) and let it stir under reflux conditions for 2 h. After the reaction has cooled to room temperature water was added and the mixture was extracted with ethyl acetate. The combined organic layers were washed with brine, dried over Na_2_SO_4_ and evaporated. The crude product was purified by flash chromatography (*n*-hexane/ethyl acetate 15:3) to give 8 as white solid (71 mg, 63 % yield). ESI-MS *m/z* 272.4; 274.4 [M+H]^+^; ^1^H-NMR (400 MHz, DMSO-*d_6_*) δ 10.64 (s, 1H), 6.87 (d, *J* = 8.6 Hz, 1H), 6.52 (dd, *J* = 8.6, 2.9 Hz, 1H), 6.49 (d, *J* = 2.7 Hz, 1H), 4.49 (s, 2H), 4.24 – 4.20 (m, 2H), 3.78 – 3.74 (m, 2H); ^13^C-NMR (101 MHz, DMSO-*d_6_*) δ 165.76, 154.06, 137.86, 128.27, 117.05, 108.69, 103.17, 68.74, 67.32, 32.01; HPLC System 1: t_R_ = 18.8 min (UV detection at λ = 254 nm), System 2: t_R_ = 16.8 min (UV detection at λ = 254 nm).

### 6-(3-Bromopropoxy)-2H-benzo[b][1,4]oxazin-3(4H)-one (**9**)

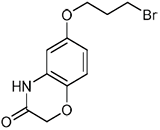

Compound **9** was prepared according to the protocol of **5** by refluxing a suspension of 7 (65 mg, 0.39 mmol), anhydrous K_2_CO_3_ (65 mg, 0.47 mmol) and 1,3-dibromopropane (0.10 mL, 0.98 mmol) in ethanol (0.8 mL) for 2 h. The crude product was purified by flash chromatography (hexane/ethyl acetate 2:1) to give 9 as a white wax (67 mg, 59% yield). ESI-MS *m/z* 285.9 [M+1]^+^; IR (NaCl) ν (cm^-1^): 3214, 3062, 2956, 1695, 1609, 1516, 1387, 1316, 1265, 1211, 1049, 898, 758; ^1^H-NMR (DMSO-*d_6_*, 600 MHz) δ (ppm): 2.18 – 2.23 (m, 2H), 3.64 (t, *J* = 6.5 Hz, 2H), 3.98 (t, *J* = 6 Hz, 2H), 4.48 (s, 2H), 6.48 (d, *J* = 2.9 Hz, 1H), 6.5 (dd, *J* = 8.7, 2.9 Hz, 1H), 6.85 (d, *J* = 8.7 Hz, 1H), 10.6 (s, 1H); ^13^C-NMR (DMSO-*d_6_*, 600 MHz) δ (ppm): 31.25, 31.76, 65.72, 66.82, 102.46, 108.20, 116.55, 128.02, 137.33, 153.63, 165.21.

### 6-(4-Bromobutoxy)-2H-benzo[b][1,4]oxazin-3(4H)-one (**10**)

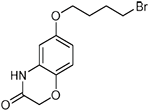

To a suspension of **7** (68 mg, 0.41 mmol) and anhydrous K_2_CO_3_ (68 mg, 0.49 mmol) in ethanol (5 mL) was added 1,4-dibromobutane (0.12 mL, 1.0 mmol) dropwise and the resulting mixture was refluxed for 2 h. After cooling to room temperature water was added and the aqueous phase was extracted with ethyl acetate. The organic layers were combined and washed with brine. After drying over Na_2_SO_4_ the organic solvent was evaporated and the crude product was purified by flash chromatography (*n*-hexane/ethyl acetate 4:1) to give 10 as an off-white solid (66 mg, 53 % yield). ESI-MS *m/z* 300.9; 302.9 [M+H]^+^; ^1^H-NMR (400 MHz, DMSO-*d_6_*) δ 10.61 (s, 1H), 6.85 (d, *J* = 8.6 Hz, 1H), 6.48 (dd, *J* = 8.6, 2.9 Hz, 1H), 6.46 (d, *J* = 2.7 Hz, 1H), 4.48 (s, 2H), 3.91 (t, *J* = 6.2 Hz, 2H), 3.59 (t, *J* = 6.7 Hz, 2H), 1.98 – 1.90 (m, 2H), 1.84 – 1.73 (m, 2H); ^13^C-NMR (101 MHz, DMSO) δ 165.28, 153.84, 137.21, 128.02, 116.54, 108.13, 102.47, 67.03, 66.86, 34.87, 29.08, 27.42; HPLC System 1: t_R_ = 19.7 min (UV detection at λ = 254 nm), System 2: t_R_ = 17.4 min (UV detection at λ = 254 nm).

### 6-(3-((2-(2-Methoxyphenoxy)ethyl)amino)propoxy)-2H-benzo[b][1,4]oxazin-3(4H)-one xTFA (**ST171**)

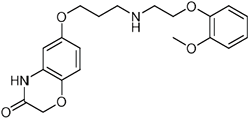

A solution of **9** (66 mg, 0.23 mmol), 2-methoxyphenoxyethylamine (*96*) (96 mg, 0.58 mmol) and KI (58 mg, 0.35 mmol) in CH_3_CN (3.3 mL) was refluxed under nitrogen atmosphere for 5 h. After the reaction has cooled to room temperature, water was added and the aqueous phase was extracted with CH_2_Cl_2_. The combined organic layers were washed with brine, dried over Na_2_SO_4_ and evaporated. The crude product was purified by preparative HPLC (10-100% methanol/ 0.1% aq. trifluoroacetic acid) to give **ST171** as a white solid (64 mg, 74% yield). ESI-MS *m/z* 373.1 [M+1]^+^; HR-MS calculated: 373.17580, found: 373.17540; IR (NaCl) ν (cm^-1^): 3431, 3065, 2922, 1685, 1610, 1507, 1396, 1326, 1256, 1203, 1126, 1054, 797; ^1^H-NMR (CDCl_3_, 600 MHz) δ (ppm): 2.24 – 2.31 (m, 2H), 3.41 (t, *J* = 5.8 Hz, 2H), 3.49 (t, *J* = 4.7 Hz, 2H), 3.80 (s, 3H), 4.08 (t, *J* = 5.2 Hz, 2H), 4.29 – 4.34 (m, 4H), 6.40 – 6.45 (m, 2H), 6.76 (d, *J* = 8.6 Hz, 1H), 6.88 – 6.97 (m, 3H), 7.01 – 7.06 (m, 1H), 8.73 (s, 1H), 9.85 (s(b), 2H); ^13^C-NMR (CDCl_3_, 600 MHz) δ (ppm): 25.76, 46.53, 47.55, 55.75, 65.72, 66.43, 67.24, 103.21, 108.8, 112.3 116.79, 116.96, 121.47, 123.8, 127.01, 138.04, 146.66, 149.91, 153.46, 165.6; HPLC System 1: t_R_ = 16.5 min, purity: 98% (UV detection at λ = 254 nm), System 2: t_R_ = 11.6 min, purity: 98% (UV detection at λ = 254 nm).

### 3-(2-Methoxyphenoxy)propyl-1-amine (**11**)

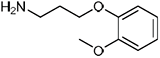

A mixture of 2-methoxyphenol (2.98 mL, 26.9 mmol), acrylonitrile (17.9 mL, 270 mmol), K_2_CO_3_ (187 mg, 1.35 mmol) and *tert*-butanol (0.24 mL, 2.7 mmol) was heated to reflux under argon atmosphere. After 4 h K_2_CO_3_ (187 g, 1.35 mmol) was added, and the reaction was stirred for further 48 h under reflux conditions. After cooling to room temperature phosphoric acid 85 % (0.12 mL, 2.2 mmol) was added dropwise and the reaction was stirred for further 30 min. Subsequently ethyl acetate was added, and the mixture was washed with 1 N NaOH solution. The resulting aqueous phase was extracted with ethyl acetate again and the combined organic layers were washed with 1 N NaOH and 10 % phosphoric acid solution, dried over Na_2_SO_4_ and evaporated to give 3-(2-methoxyphenoxy)propanenitrile as light brown solid (2.49 g, 52 % yield). ESI-MS *m/z* 177.8 [M+H]^+^; ^1^H-NMR (400 MHz, CDCl_3_) δ 7.05 – 6.97 (m, 1H), 6.95 – 6.87 (m, 3H), 4.25 (t, *J* = 6.7 Hz, 2H), 3.87 (s, 3H), 2.85 (t, *J* = 6.7 Hz, 2H).

A solution of the obtained 3-(2-methoxyphenoxy)propanenitrile (1.40 g, 7.90 mmol) in anhydrous THF (6 mL) was added dropwise a 1 M solution of borane tetrahydrofuran complex in THF (8.75 mL, 8.75 mmol). The reaction mixture was heated to 65°C for 15 h under argon atmosphere. After cooling to 0°C a 5 M NaOH solution (9.7 mL) was added slowly, and the reaction was heated up to 65°C again for further 4 h. After the reaction has cooled to room temperature ethyl acetate was added and the organic phase was washed with water and brine. After drying over Na_2_SO_4_ the organic solvent was evaporated in vacuo to give 11 as yellowish oil (874 mg, 61 % yield). ESI-MS *m/z* 181.8 [M+H]^+^; ^1^H-NMR (400 MHz, CDCl_3_) δ 6.95 – 6.83 (m, 4H), 4.11 (t, *J* = 6.2 Hz, 2H), 3.86 (s, 3H), 2.93 (t, *J* = 6.7 Hz, 2H), 2.02 – 1.93 (m, 2H), 1.65 (s, 2H); ^13^C-NMR (151 MHz, CDCl_3_) δ 149.47, 148.33, 121.25, 120.91, 113.38, 111.72, 67.58, 55.90, 39.59, 29.40; HPLC System 1: t_R_ = 11.0 min (UV detection at λ = 254 nm), System 2: t_R_ = 11.7 min (UV detection at λ = 254 nm).

### N-(2-(2-(Methylthio)phenoxy)ethyl)acetamide (**12**)

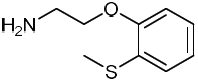

A mixture of 2-(methylthio)phenol (0.26 mL, 2.1 mmol) and 2-methyl-2-oxazoline (0.18 mL, 2.1 mmol) was stirred in a sealed tube at 160°C overnight under argon atmosphere. The crude product was purified by flash chromatography (isohexane/ ethyl acetate 3:1) to yield *N*-(2-(2-(methylthio)phenoxy)ethyl)acetamide as yellow oil (0.29 g, 61% yield). ESI-MS *m/z* 225.8 [M+H]^+^; ^1^H-NMR (400 MHz, DMSO-d_6_) δ 8.04 (s, 1H), 7.18 – 7.08 (m, 2H), 7.00 – 6.94 (m, 2H), 4.01 (t, *J* = 6.0 Hz, 2H), 3.40 (q, *J* = 5.9 Hz, 2H), 2.37 (s, 3H), 1.83 (s, 3H).

A solution of the obtained *N*-(2-(2-(methylthio)phenoxy)ethyl)acetamide (200 mg, 887 µmol) in MeOH (5.0 mL) was added carefully HCl_conc_ (5.0 mL). The resulting mixture was refluxed for 24 h. After the solution has cooled to room temperature 5 N NaOH solution was added to adjust the pH to 9 - 10 and the resulting aqueous phase was extracted with ethyl acetate. The organic layers were combined, dried over Na_2_SO_4_ and evaporated in vacuo to yield 12 as yellowish oil (154 mg, 95 % yield). ESI-MS *m/z* 183.9 [M+H]^+^; ^1^H-NMR (400 MHz, DMSO-*d_6_*) δ 7.16 – 7.08 (m, 2H), 7.00 – 6.91 (m, 2H), 3.97 (t, J = 5.6 Hz, 2H), 2.90 – 2.85 (m, 2H), 2.37 (s, 3H); ^13^C-NMR (101 MHz, DMSO-*d_6_*) δ 155.21, 127.57, 125.86, 125.35, 121.68, 112.11, 71.35, 41.40, 13.87; HPLC System 1: t_R_= 12.4 min (UV detection at λ = 254 nm), System 2: t_R_= 12.1 min (UV detection at λ = 254 nm).

### 6-(3-((3-(2-Methoxyphenoxy)propyl)amino)propoxy)-2H-benzo[b][1,4]oxazin-3(4H)-one (**TA12**)

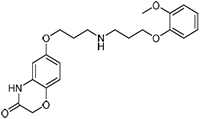

A suspension of 9 (15 mg, 52 µmol), **11** (24 mg, 0.13 mmol) and KI (13 mg, 79 µmol) in acetonitrile (3.3 mL) was heated in a sealed tube to reflux for 5 h under argon atmosphere. After the reaction has cooled to room temperature water was added and the aqueous phase was extracted with CH_2_Cl_2_. The combined organic layers were washed with brine, dried over Na_2_SO_4_ and evaporated under vacuo. The crude product was purified by preparative HPLC (10 - 90 % CH_3_CN/ 0.1 % aq. trifluoroacetic acid) to give TA12 as white TFA salt (15 mg, 57 % yield). ESI-MS *m/z* 387.2 [M+H]^+^; HR-MS calculated: 387.1914 [M+H]^+^, found: 387.1912 [M+H]^+^; ^1^H-NMR (600 MHz, CDCl3) δ 9.34 (s, 2H), 8.88 (s, 1H), 7.01 – 6.96 (m, 1H), 6.92 – 6.85 (m, 3H), 6.77 (d, J = 8.8 Hz, 1H), 6.44 (d, J = 2.7 Hz, 1H), 6.40 (dd, J = 8.8, 2.8 Hz, 1H), 4.35 (s, 2H), 4.17 (t, J = 5.5 Hz, 2H), 4.06 (t, J = 5.6 Hz, 2H), 3.84 (s, 3H), 3.41 – 3.38 (m, 2H), 3.35 – 3.33 (m, 2H), 2.34 – 2.25 (m, 4H); ^13^C-NMR (151 MHz, CDCl3) δ 165.96, 153.72, 149.34, 147.20, 137.81, 126.92, 122.77, 121.18, 116.99, 114.56, 111.76, 109.51, 102.84, 69.00, 67.17, 65.31, 55.83, 47.53, 45.42, 26.05, 24.98; HPLC System 1: t_R_ = 15.3 min, purity: 97 % (UV detection at λ = 254 nm), System 2: t_R_ = 14.3 min, purity: 98 % (UV detection at λ = 254 nm).

### 6-(4-((2-(2-Methoxyphenoxy)ethyl)amino)butoxy)-2H-benzo[b][1,4]oxazin-3(4H)-one (**TA13**)

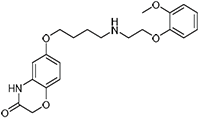

Compound **TA13** was prepared according to the protocol of **TA12** by refluxing a suspension of 10 (15 mg, 50 µmol), 2-(2-methoxyphenoxy)ethan-1-amine (*96*) (21 mg, 0.12 mmol) and KI (12 mg, 75 µmol) in acetonitrile (3.3 mL) under argon atmosphere for 2.5 h. After cooling to room temperature water was added and the aqueous phase was extracted with CH_2_Cl_2_. The combined organic layers were washed with brine, dried over Na_2_SO_4_ and evaporated. Purification of the crude product by preparative HPLC (10 - 90 % CH_3_CN/ 0.1 % aq. trifluoroacetic acid) yielded **TA13** as colorless TFA salt (13 mg, 52 % yield). ESI-MS *m/z* 387.2 [M+H]^+^; HR-MS calculated: 387.1915 [M+H]^+^, found: 387.1917 [M+H]^+^; ^1^H-NMR (600 MHz, DMSO-*d_6_*) δ 10.65 (s, 1H), 8.82 (s, 2H), 7.05 – 7.01 (m, 2H), 7.00 – 6.97 (m, 1H), 6.92 – 6.89 (m, 1H), 6.88 – 6.84 (m, 1H), 6.50 – 6.48 (m, 2H), 4.48 (s, 2H), 4.22 (t, J = 5.2 Hz, 2H), 3.91 (t, J = 5.8 Hz, 2H), 3.78 (s, 3H), 3.37 – 3.32 (m, 2H), 3.14 – 3.07 (m, 2H), 1.88 – 1.71 (m, 4H); ^13^C-NMR (151 MHz, DMSO-*d_6_*) δ 165.68, 154.23, 149.89, 147.51, 137.66, 128.49, 122.75, 121.15, 116.95, 115.40, 112.81, 108.51, 102.92, 67.61, 67.29, 65.41, 55.90, 47.47, 46.34, 26.24, 22.80; HPLC System 1: t_R_ = 16.1 min, purity: 98 % (UV detection at λ = 254 nm), System 2: t_R_ = 15.2 min, purity: 99 % (UV detection at λ = 254 nm).

### 6-(2-((2-(2-Methoxyphenoxy)ethyl)amino)ethoxy)-2H-benzo[b][1,4]oxazin-3(4H)-one (**TA14**)

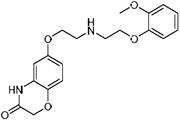

Compound **TA14** was prepared as described for **TA12**, by stirring a suspension of 8 (39 mg, 0.14 mmol), 2-(2-methoxyphenoxy)ethan-1-amine (*96*) (60 mg, 0.36 mmol) and KI (36 mg, 0.21 mmol) in acetonitrile (3.3 mL) in a sealed tube under argon atmosphere and reflux conditions for 2.5 h. After the reaction has cooled to room temperature water was added and the resulting aqueous phase was extracted with CH_2_Cl_2_. The combined organic layers were washed with brine, dried over Na_2_SO_4_ and evaporated in vacuo. The crude product was purified by preparative HPLC (10 - 90 % CH_3_CN/ 0.1 % aq. trifluoroacetic acid) to give **TA14** as white TFA salt (32 mg, 48 % yield). ESI-MS *m/z* 359.1 [M+H]^+^; HR-MS calculated: 359.1602 [M+H]^+^, found: 359.1599 [M+H]^+^; ^1^H-NMR (400 MHz, DMSO-*d_6_*) δ 10.75 (s, 1H), 8.86 (s, 2H), 7.07 – 7.03 (m, 1H), 7.02 – 6.97 (m, 2H), 6.93 – 6.92 (m, 1H), 6.91 – 6.88 (m, 1H), 6.60 – 6.53 (m, 2H), 4.51 (s, 2H), 4.25 (t, J = 5.1 Hz, 2H), 4.19 (t, J = 5.0 Hz, 2H), 3.78 (s, 3H), 3.48 (t, J = 4.9 Hz, 2H), 3.43 (t, J = 5.1 Hz, 2H); ^13^C-NMR (101 MHz, DMSO-*d_6_*) δ 165.28, 153.03, 149.43, 147.02, 137.74, 128.11, 122.37, 120.72, 116.60, 114.96, 112.32, 108.12, 102.89, 66.82, 64.94, 63.80, 55.46, 46.51, 46.31; HPLC System 1: t_R_ = 15.3 min, purity: 98 % (UV detection at λ = 254 nm), System 2: t_R_ = 14.4 min, purity: 97 % (UV detection at λ = 254 nm).

### 6-(3-((2-(2-(Methylthio)phenoxy)ethyl)amino)propoxy)-2H-benzo[b][1,4]oxazin-3(4H)-one (**TA48**)

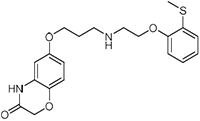

Compound **TA48** was prepared as described for **TA13** by refluxing a mixture of 9 (70 mg, 0.32 mmol), 12 (0.14 g, 0.79 mmol) and KI (78 mg, 0.47 mmol) in acetonitrile (3.8 mL) under argon atmosphere for 4 h. After cooling to room temperature water was added and the aqueous phase was extracted with ethyl acetate. The organic layers were combined, washed with brine, dried over MgSO_4_ and evaporated in vacuo. The crude product was purified by preparative HPLC (10 - 90 % CH_3_CN/ 0.1 % aq. trifluoroacetic acid) to give **TA48** as white TFA salt (62 mg, 39 % yield). ESI-MS *m/z* 389.0 [M+H]^+^; HR-MS calculated: 389.1530 [M+H]^+^, found: 389.1531 [M+H]^+^; ^1^H-NMR (600 MHz, DMSO-*d_6_*) δ 10.68 (s, 1H), 8.72 (s, 2H), 7.21 – 7.15 (m, 2H), 7.07 – 7.03 (m, 2H), 6.88 (d, J = 8.7 Hz, 1H), 6.50 (dd, J = 8.7, 2.9 Hz, 1H), 6.48 (d, J = 2.8 Hz, 1H), 4.50 (s, 2H), 4.30 (t, J = 5.1 Hz, 2H), 3.99 (t, J = 5.9 Hz, 2H), 3.46 – 3.41 (m, 2H), 3.31 – 3.25 (m, 2H), 2.38 (s, 3H), 2.11 – 2.06 (m, 2H); ^13^C-NMR (151 MHz, DMSO-*d_6_*) δ 165.12, 153.74, 153.41, 137.22, 127.89, 127.28, 125.32, 125.05, 122.14, 116.38, 112.10, 107.90, 102.36, 66.69, 64.94, 64.62, 45.85, 45.16, 25.54, 13.34; HPLC System 1: t_R_= 15.8 min, purity: 98 % (UV detection at λ = 254 nm), System 2: t_R_= 14.6 min, purity: 99 % (UV detection at λ = 254 nm).

---

**Supplementary Figure 1:**
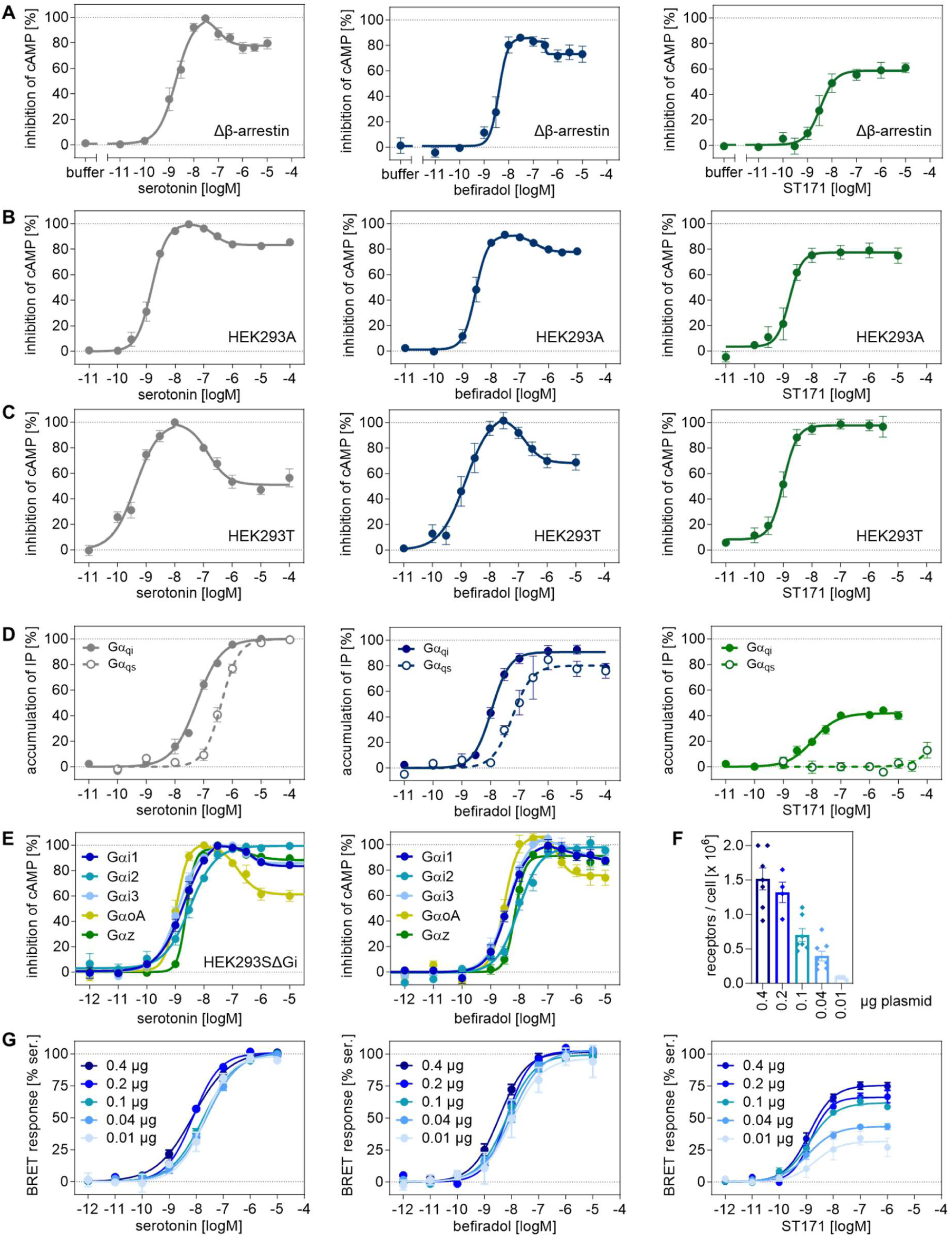
Functional profiles of serotonin, befiradol, and ST171 in different cell lines (A-C) Inhibition of cAMP accumulation by serotonin, befiradol, and ST171 in HEK293A cells deficient of β-arrestins, HEK293A cells and HEK293T cells. Data was obtained with the GloSensor and is normalized to the maximal response of serotonin. Data is shown with ± SEM of 5-7 independent experiments, performed as triplicates (for A, B) or as duplicates (C). (D) Gα_i_ and Gα_s_ selective signaling determined with the intracellular IP accumulation assay with hybrid Gα_qi_ and Gα_qs_ proteins. Data is shown with ± SEM of 6-8 independent experiments, each performed as duplicates. (E) Inhibition of cAMP accumulation by serotonin and befiradol in HEK293AΔG_i_ co transfected with Gαi subunits (Gα_i1_, Gα_i2_, Gα_i3_, Gα_z_, Gα_oA_). Data was obtained with the GloSensor, normalized to the maximal response of serotonin and is shown with ± SEM of 5 experiments, each performed as triplicates. (F) 5-HT_1A_ receptor densities in HEK cells. The numbers of receptors per cell were determined in radioligand saturation experiments with [^3^H]WAY100,635 in parallel to the BRET Gα_i_ activation assay. Data was derived from 4-8 single experiments each performed in triplicate. (G) 5-HT_1A_R mediated Gα_i_ activation by serotonin, befiradol and ST171 in HEK293T cell expressing different 5-HT_1A_R densities according to the transfection of 0.01-0.4 µg plasmid/culture plate. Data was determined with a Gα_i1_ BRET biosensor and was normalized to the maxium effect of serotonin. Results are displayed as mean ± SEM of 5-17 experiments each conducted in duplicates.

**Supplementary Figure 2:**
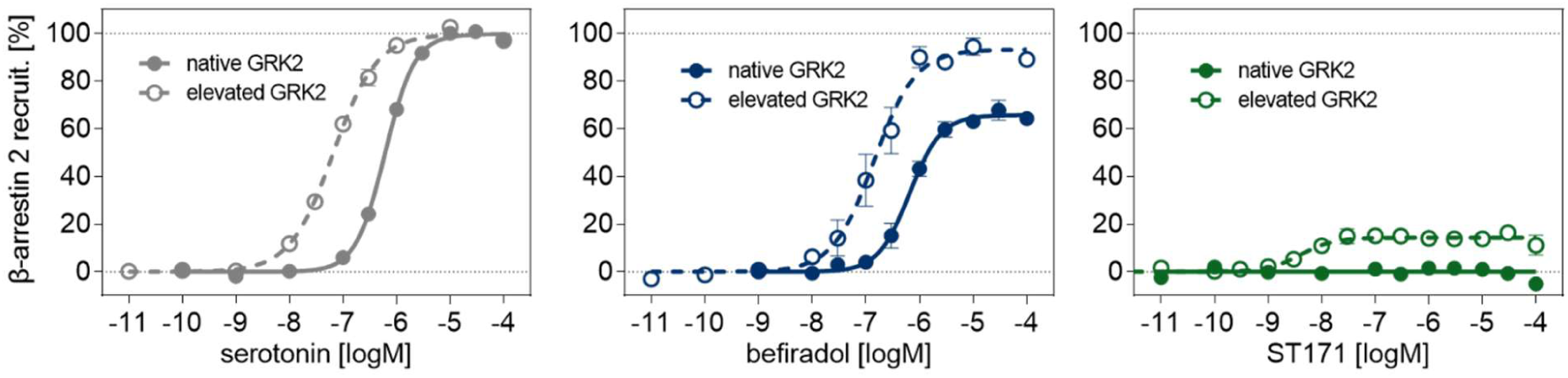
Arrestin recruitment to the 5-HT_1A_R for serotonin, befiradol, and ST171 with native GRK2 levels (- GRK) and with elevated GRK2 levels (+GRK) determined with the PathHunter β-arrestin 2 recruitment assay. Data is shown with ± SEM of 7-12 independent experiments, each performed as duplicates.

**Supplementary Figure 3:**
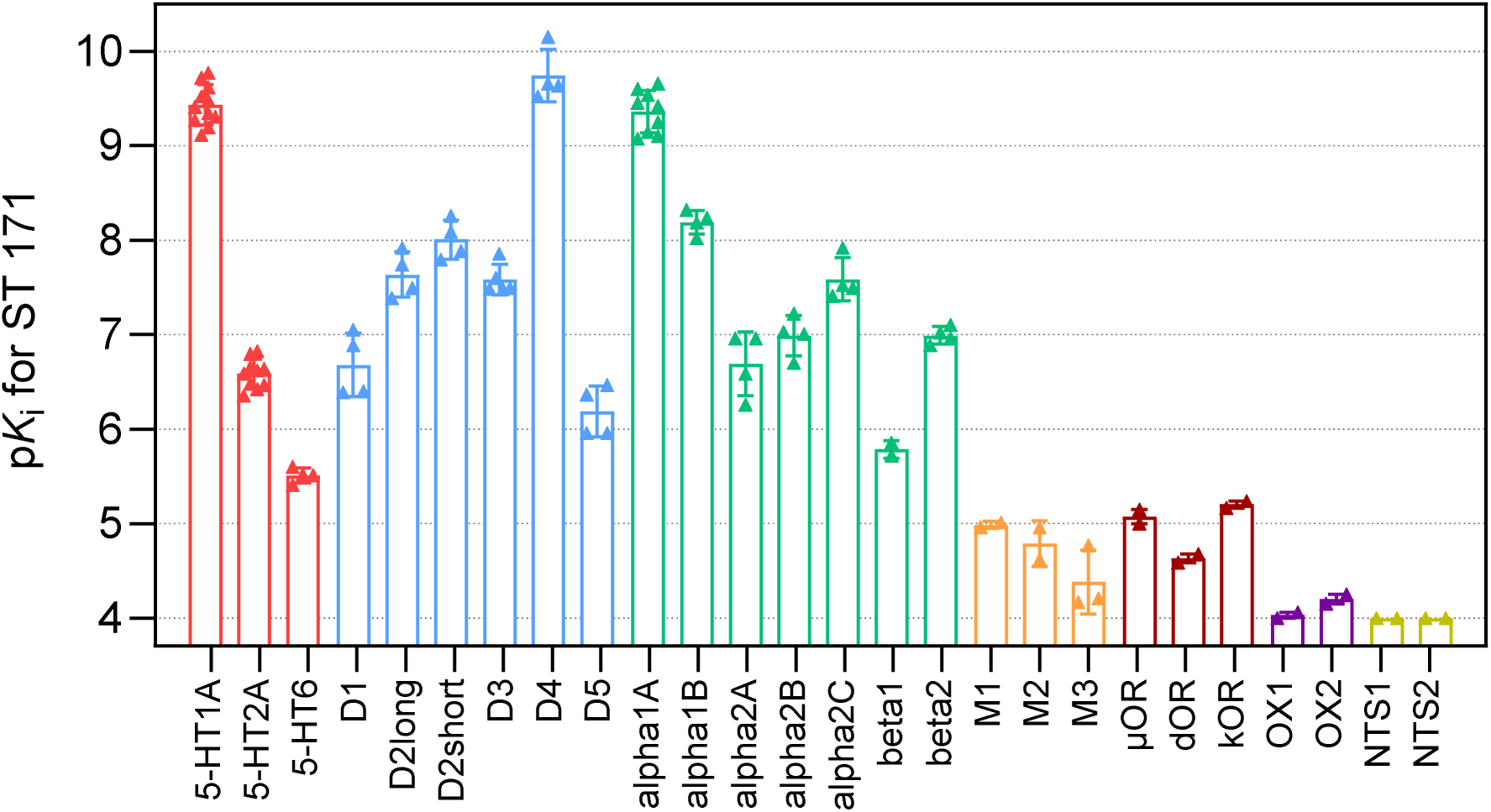
Affinity of ST171 for different GPCRs. pK_i_ values for 26 representative class A GPCRs including serotonergic (red), dopaminergic (blue), adrenergic (green), muscarinic acetylcholine (orange), opioid (brown), orexin (purple), and neurotensin (yellow) receptors were determined by radioligand competition. Bars represent the mean ±SD of 2-11 individual experiments, each performed in triplicates.

**Supplementary Figure 4:**
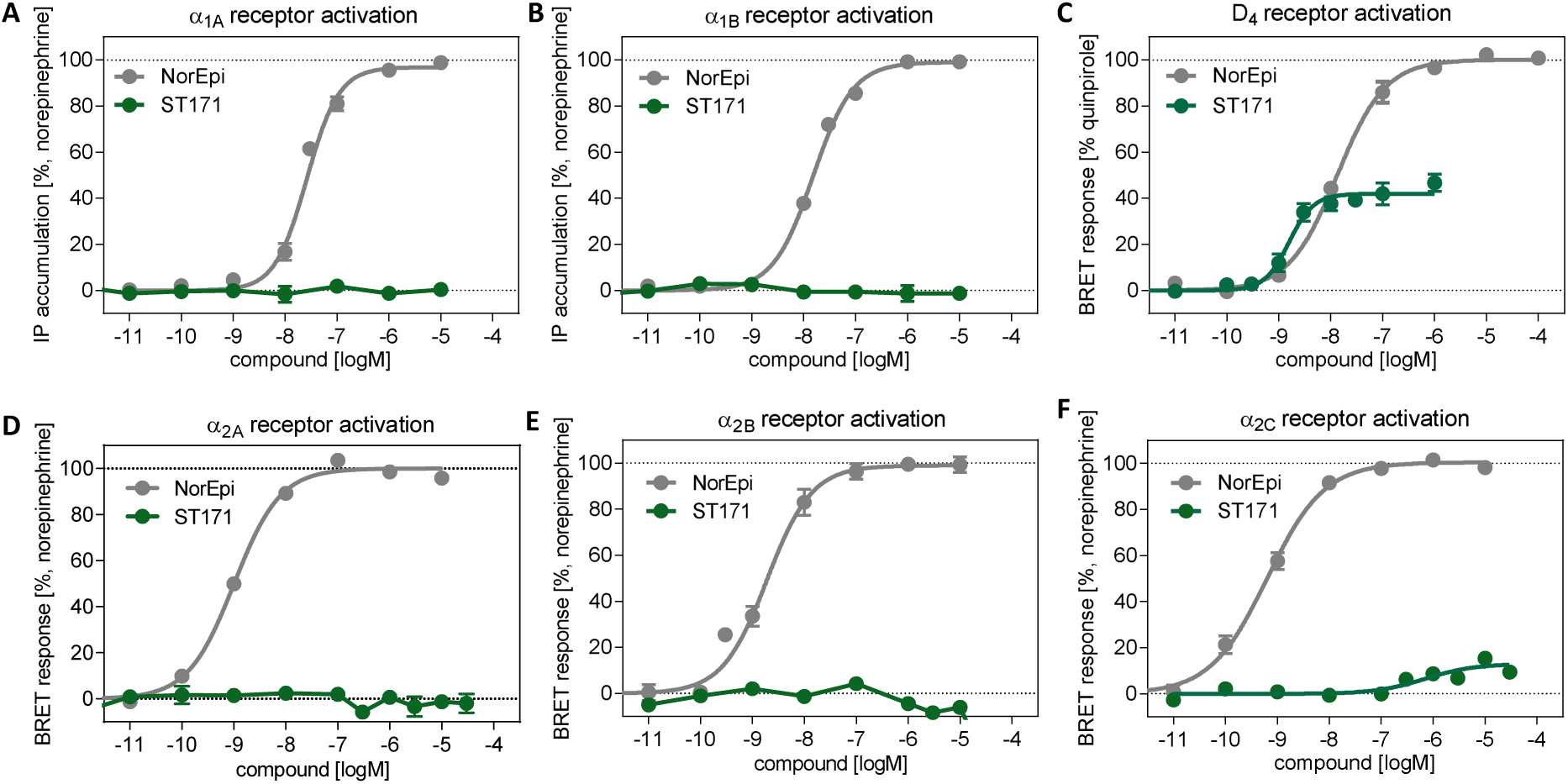
Functional properties of ST171 at selected GPCRs. ST171 showed no activation at the subtypes of adrenergic alpha receptors relative to the reference norepinephrine and at the dopamine receptor subtype D_4_. (A,B) Test on receptor activation at α_1A_AR and α_1B_AR determined in an IP accumulation assay (IP-One, PerkinElmer). (C) Recepotr activation of D_4_ determined in a Gα_i1_ biosensor-based BRET assay. (D,E,F) Functional activity at α_2A_AR, α_2B_AR, and α_2C_AR measured in a Gα_i1_ biosensor-based BRET assay. Graphs show mean curves of individual experiments each performed in duplicates with the number of individual experiments for A) NorEpi (n=5), ST171 (n=4), B) NorEpi (n=7), ST171 (n=3), C) Quinpirole (n=8), ST171 (n=7), D) NorEpi (n=14), ST171 (n=4), E) NorEpi (n=5), ST171 (n=3), F) NorEpi (n=5), ST171 (n=6).

**Supplementary Figure 5:**
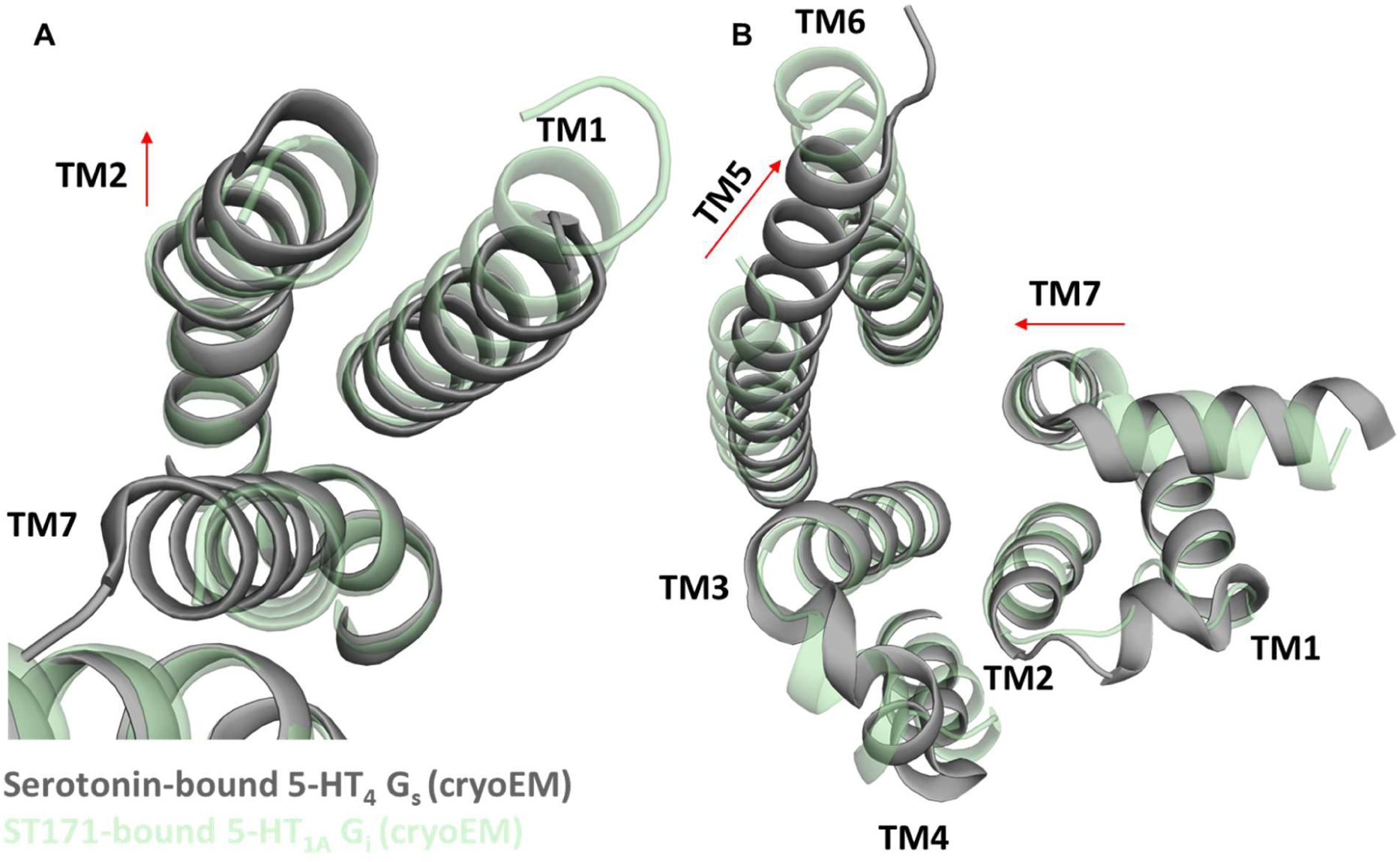
Comparison of the serotonin-bound G_s_-5-HT_4_R complex (grey) and the ST171-bound G_i_-5-HT_1A_R complex (light green) from both extracellular and intracellular perspectives. (A) The extracellular view reveals an outward shift of TM2 by 1.4 Å in the G_s_-coupled complex compared to the G_i_-coupled complex. (B) The intracellular view highlights an inward shift of TM7 by 3.2 Å and a significant kink of TM5 towards TM6 in the G_s_-coupled complex relative to the G_i_-coupled complex.

**Supplementary Figure 6:**
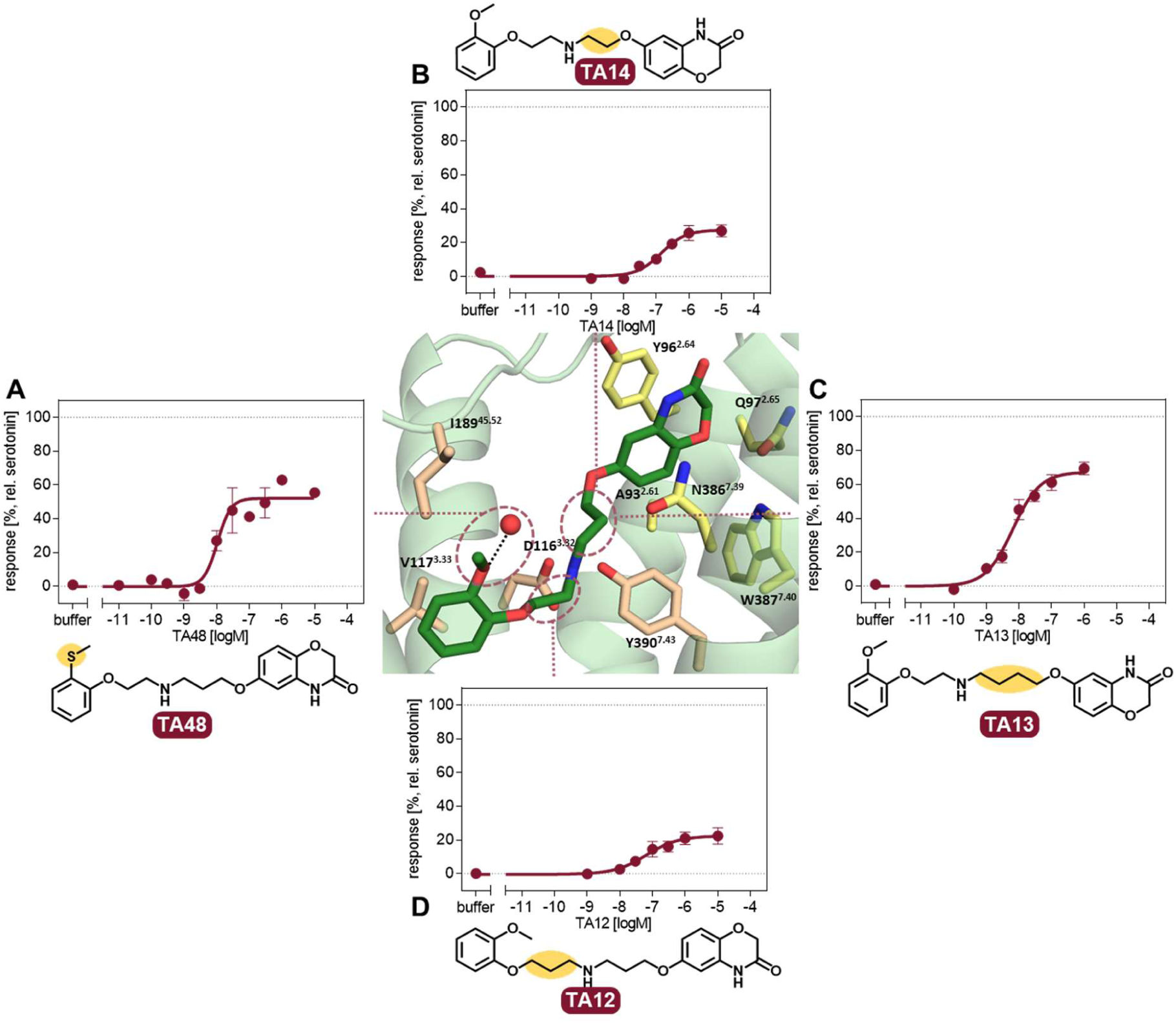
Functional activity of ST171 derivatives. Data obtained for the inhibition of the cAMP accumulation was determined with the CAMYEL assay and normalized relative to the maximal response of serotonin from 2-4 individual experiments, each performed as triplicates. (A) Substitution of the ether moiety with a thioether in TA48 resulted in a decrease in both potency and efficacy when compared to ST171. This reduction in activity can be attributed to the relatively diminished hydrogen-bond accepting capacity of the thioether group in comparison to the ether group. This observation suggests the potential involvement of the methoxy group in ligand binding, particularly in the formation of a water-mediated hydrogen bond. Modifying the positioning of the benzoxazinone moiety through linker unit shortening (B) or extension (C) leads to a substantial reduction in both potency and efficacy, thereby affirming the essential role of the benzoxazinone moiety in ligand binding. (D) Insertion of a CH_2_ unit between the head group and the secondary amine leads to a reduced binding affinity and receptor activation relative to ST171 highlighting the importance of the salt bridge to D116^3.32^.

**Supplementary Figure 7:**
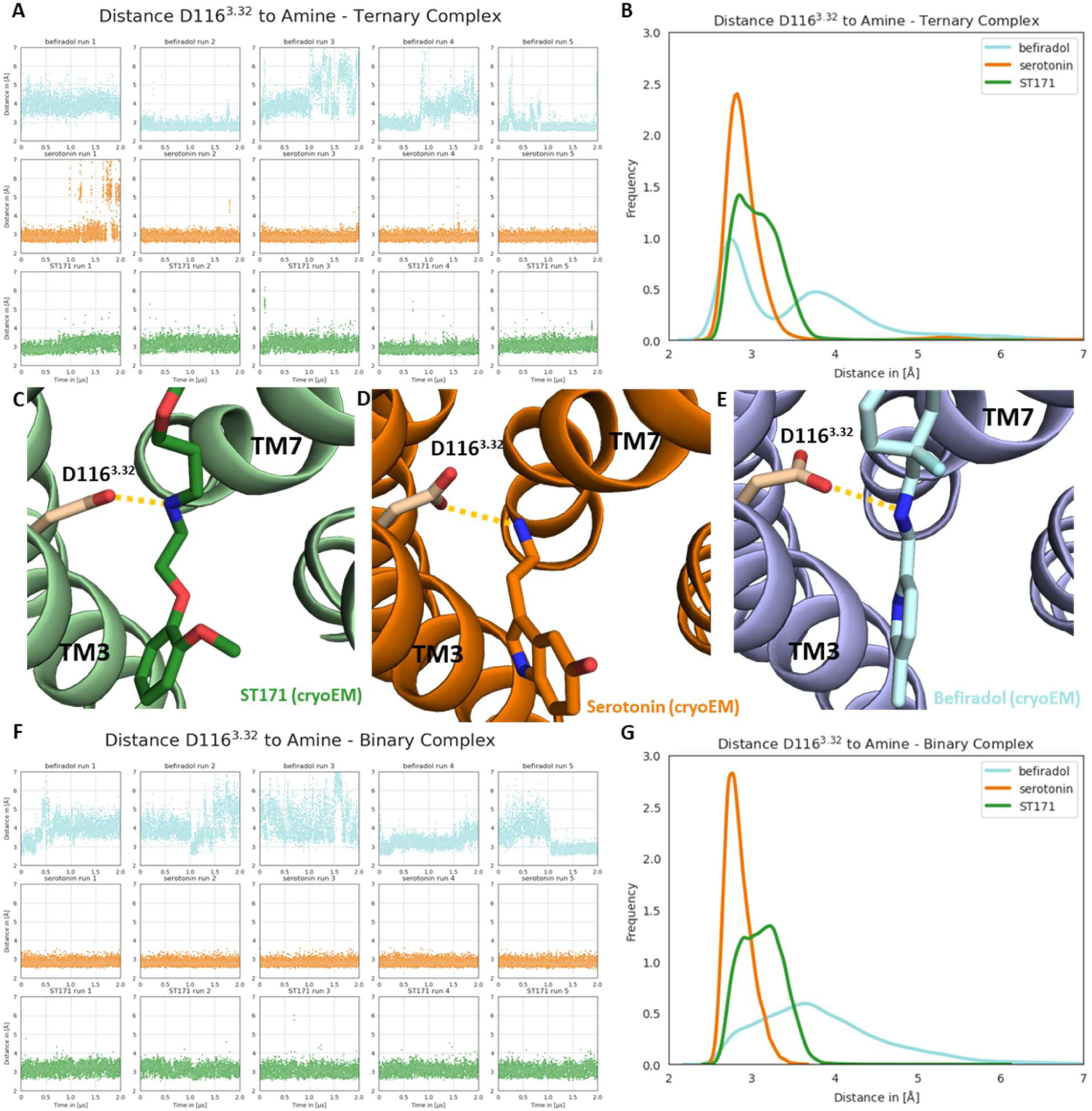
Stability analysis of the salt bridge between D116^3.32^ and basic amine of the ligands. (A) Plots show the progression of the distance between D116^3.32^ and the basic amine of the respective ligand over the course of 2 µs for 5 independent simulations of the ligand-bound 5-HT_1A_R ternary complex. (B) Probability distribution is based on the trajectory frames from all simulations combined of the respective ligand-bound 5-HT_1A_R ternary complex. A Gaussian kernel density estimaton was used. The distance between D116^3.32^ and the basic amine of the ligand in the experimental cryo-EM structures is shown for the ST171-bound (C), the serotonin-bound (D), and the befiradol-bound (E) complex. (F) Plots show the progression of the distance between D116^3.32^ and the basic amine of the respective ligand over the course of 2 µs for 5 independent simulations of the ligand-bound 5-HT_1A_R binary complex. (G) The probability distribution is based on the trajectory frames from all simulations combined of the respective ligand-bound 5-HT_1A_R binary complex. A Gaussian kernel density estimaton was used. For the plots (A, B, F, G), the distance was determined between the center of mass of the carboxy function of D116^3.32^ and the nitrogen atom of the basic amine of the ligand. In the plots (C, D, E), the distance was determined between the nearest oxygen atom of the carboxy function of D116^3.32^ and the nitrogen atom of the basic amine. The plots (B, G) depict the transition of the distance from low to high values in the befiradol-bound ternary complex compared to the respective binary complex. This dynamic behavior of befiradol within the binding pockets of the binary complex potentially suggests the intriguing signaling profile associated with befiradol.

**Supplementary Figure 8:**
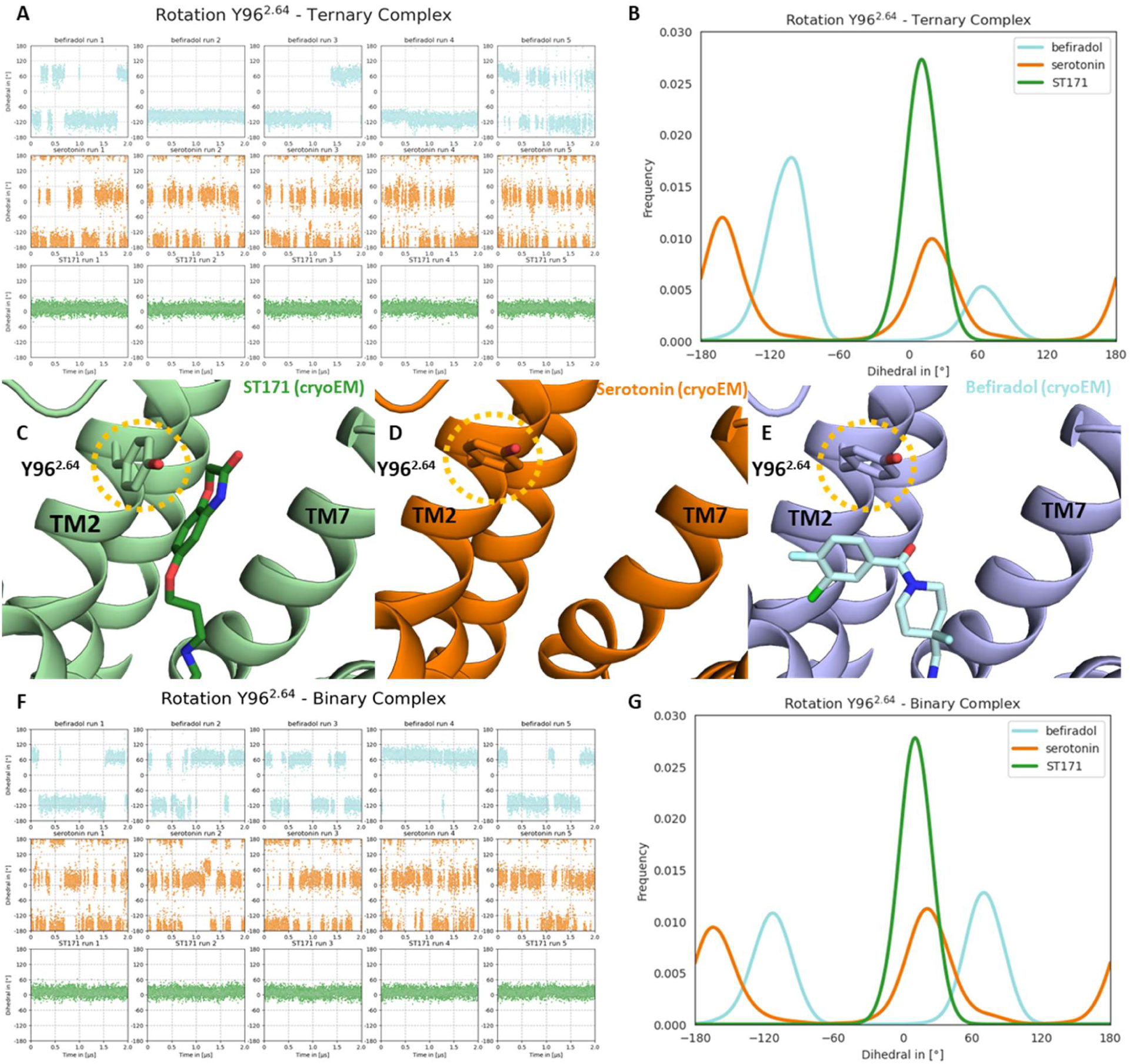
Influence of ligand biding on the torsion angle -C_α_-C_β_-C_γ_-C_δ_- of the Tyr96^2.64^ side chain. (A) Plots show the progression of the torsion angle for Y96^2.64^ over the course of 2 µs for 5 independent simulations of the respective ligand- bound 5-HT_1A_R ternary complex. (B) Probability distribution is based on the trajectory frames from all simulations combined of the respective ligand-bound 5-HT_1A_R ternary complex. The von Mises kernel was utilized as a weighted kernel density estimator. The conformation of Y96^2.64^ in the experimental cryo-EM structures is shown for the ST171-bound (C), the serotonin-bound (D), and the befiradol-bound (E) complex. (F) Plots demonstrate the progression of the dihedral angle of Y96^2.64^ over the course of 2 µs for 5 independent simulations of the respective ligand-bound 5-HT_1A_R binary complexes. (G) Probability distribution is based on the trajectory frames from all simulations combined of the respective ligand-bound 5-HT_1A_R binary complex. The von Mises kernel was utilized as a weighted kernel density estimator. Due to the ambiguity of C_δ_ in tyrosine side chains, the C_δ_ that pointed towards or was closest to the extracellular side was used for this analysis. Following the IUPAC recommendations,(*95*) positive values indicate counterclockwise rotation of the bond between C_γ_ and C_δ_ relative to the C_α_ and C_β_ bond, while negative values indicate clockwise rotation of the bond C_γ_ and C_δ_ relative to the C_α_ and C_β_ bond.

**Supplementary Figure 9:**
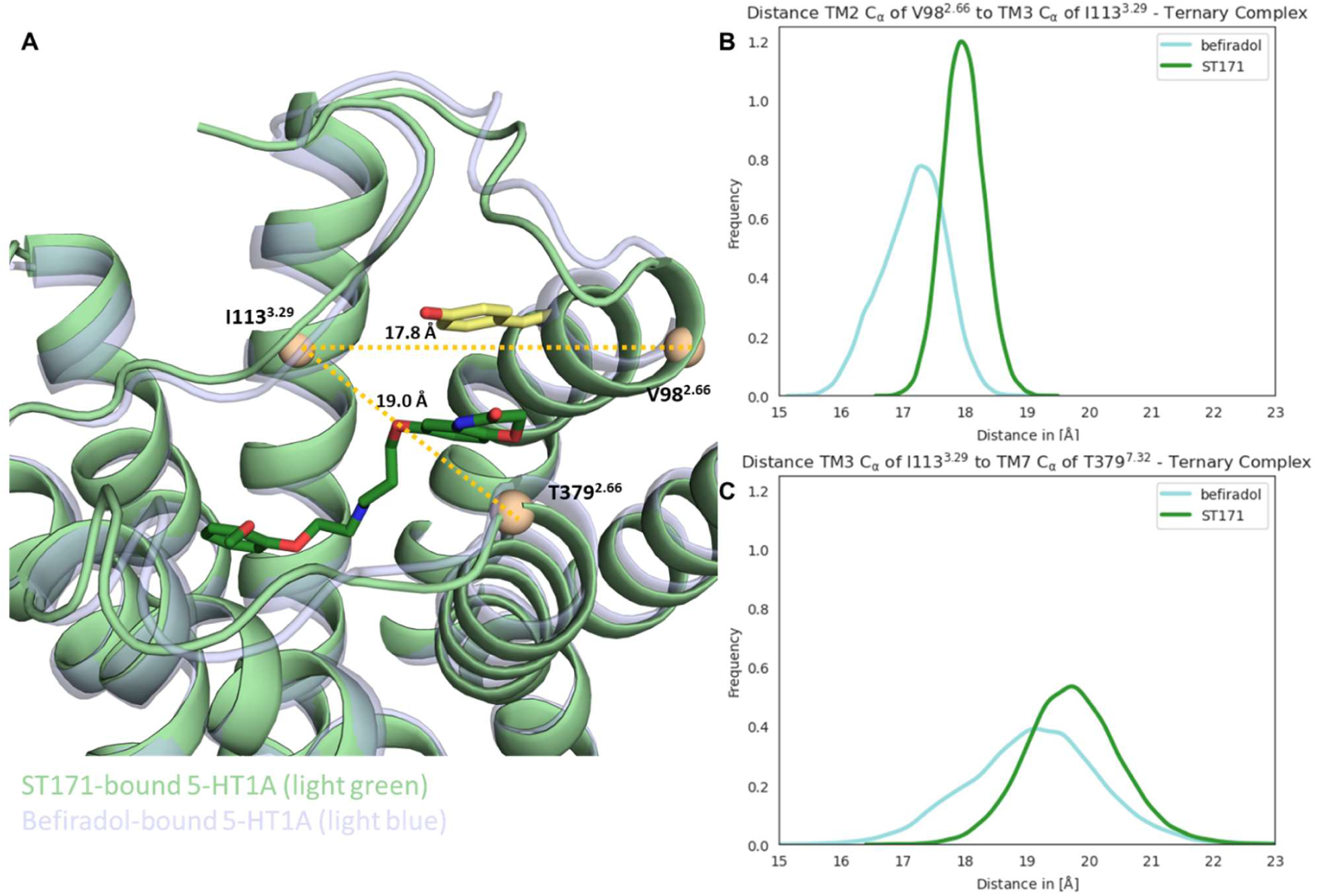
Interhelical distance TM2-TM3 (measured between C_α_ of V98^2.66^ and I113^3.29^) and TM3-TM7 (measured between C_α_ of I113^3.29^ and T379^7.32^) for the ST171- and befiradol-bound complexes. (A) Illustrates the interhelical distances TM2-TM3 and TM3-TM7 within the ST171-bound complex. Additionally, the befiradol-bound complex is depicted to demonstrate the positional changes of TM2 and TM7 in comparison to the ST171-bound complex. (B) Probability distribution for the TM2-TM3 interhelical distance is based on the trajectory frames from all simulations combined of the respective ligand-bound 5-HT_1A_R ternary complex. A Gaussian kernel density estimaton was used. (C) Probability distribution for the TM3-TM7 interhelical distance is based on the trajectory frames from all simulations combined of the respective ligand-bound 5-HT_1A_R ternary complex. A Gaussian kernel density estimaton was used. The subtle distinctions observed in the extracellular regions of TM2 and TM7 between the ST171- and the befiradol-bound complex remained present throughput MD simulations.

**Supplementary Figure 10:**
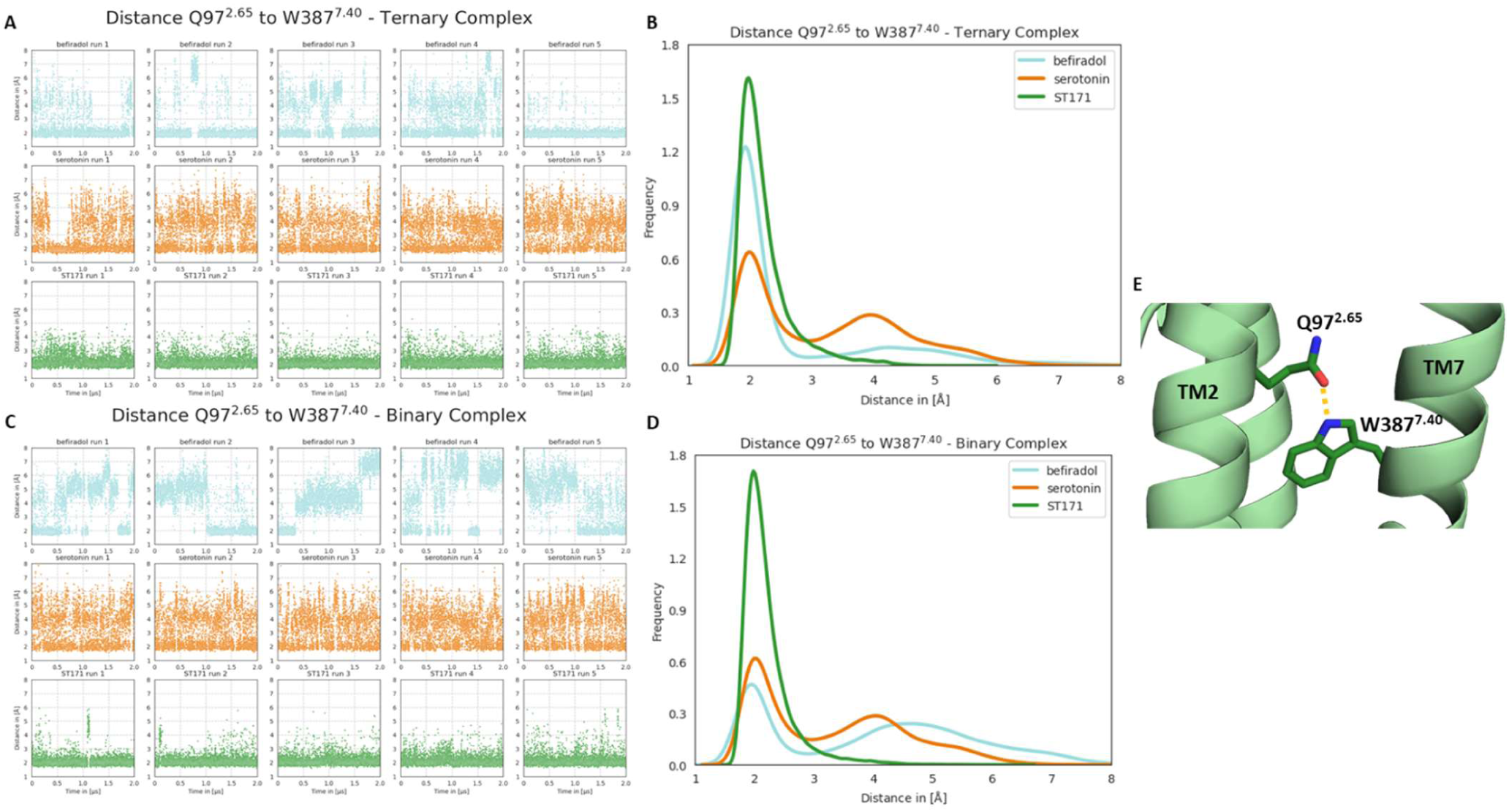
Distance between the amid group of Q97^2.65^ and the NH group W387^7.40^. (A) Plots show the progression of the distance between the amid group of Q97^2.65^ and NH group of W387^7.40^ over the course of 2 µs for 5 independent simulations of the respective ligand-bound 5-HT_1A_R ternary complex. (B) Probability distribution is based on the trajectory frames from all simulations combined of the respective ligand-bound 5-HT_1A_R ternary complex. A Gaussian kernel density estimaton was used. (C) Plots show the progression of the distance between the amid group of Q97^2.65^ and NH group of W387^7.40^ over the course of 2 µs for 5 independent simulations of the respective ligand-bound 5-HT_1A_R binary complex. (D) Probability distribution is based on the trajectory frames from all simulations combined of the respective ligand-bound 5-HT_1A_R binary complex. A Gaussian kernel density estimaton was used. (E) The distance between the amid group of Q97^2.65^ and NH group of W387^7.40^ in the experimental cryo-EM structures is shown for the ST171-bound complex. The comparison of (B, D) shows that the distance between Q97^2.65^ and W387^7.40^ in the befiradol-bound receptor is significantly influenced by whether the receptor is in a binary or ternary complex. While the ternary complex stabilizes the hydrogen bond between Q97^2.65^ and W387^7.40^, this interaction ceases gradually in the binary complex.

**Supplementary Figure 11:**
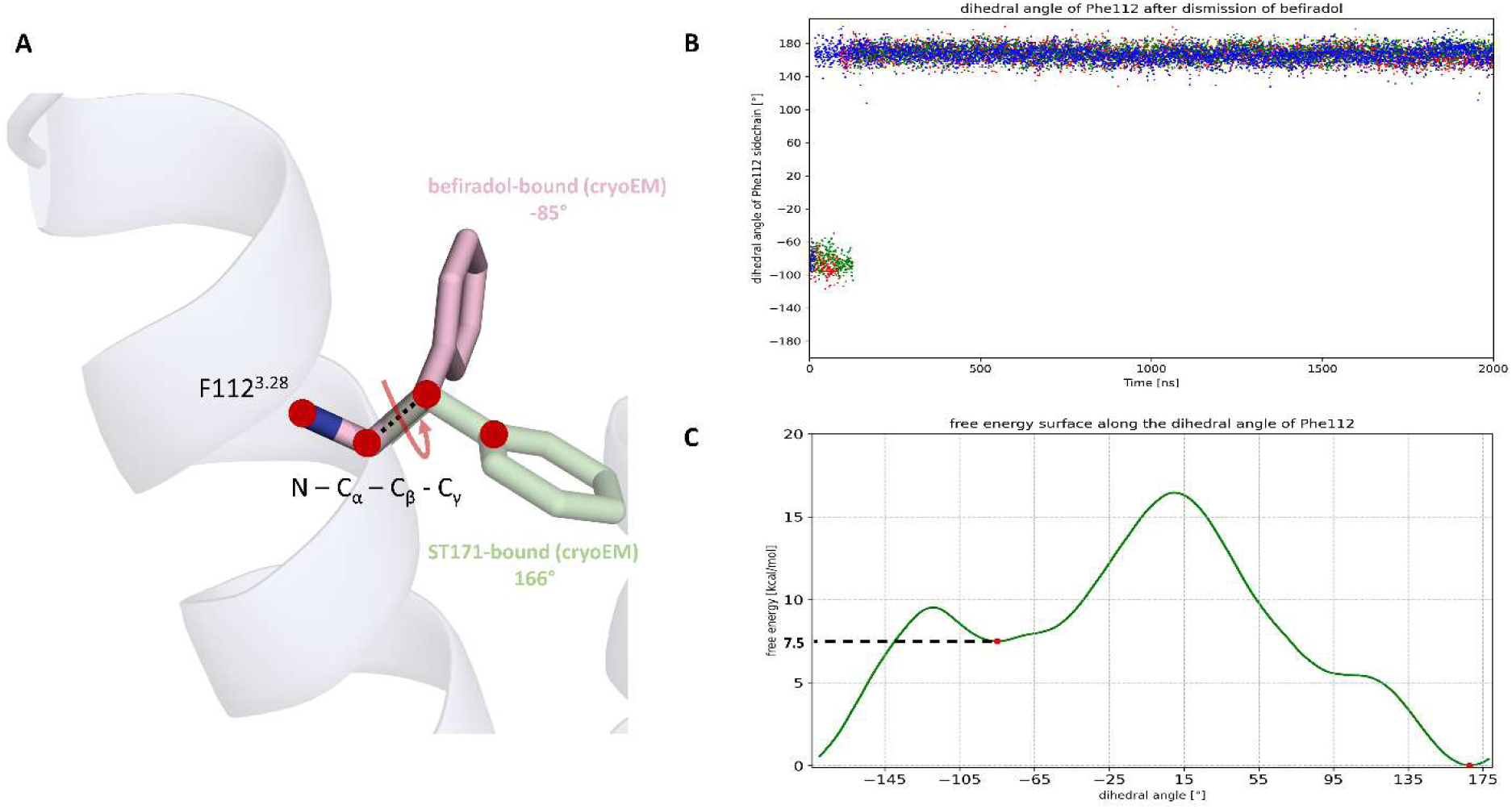
F112^3.28^ rotamers in ST171- and befiradol-bound 5-HT_1A_. (A) Comparison of the dihedral angles of the two different rotamers of F112^3.28^ in the ST171-bound and the befiradol-bound cryo-EM structures. (B) Measurement of the dihedral angle of F112^3.28^ over 3 independent runs in an unbiased molecular dynamics simulation (2 µs) of the befiradol-bound binary complex model after dismission of befiradol. (C) Free energy profile along the dihedral angle of F112^3.28^ obtained by metadynamics simulations (6.7 µs) of the befiradol-bound binary complex model after dismission of befiradol.

**Supplementary Figure 12:**
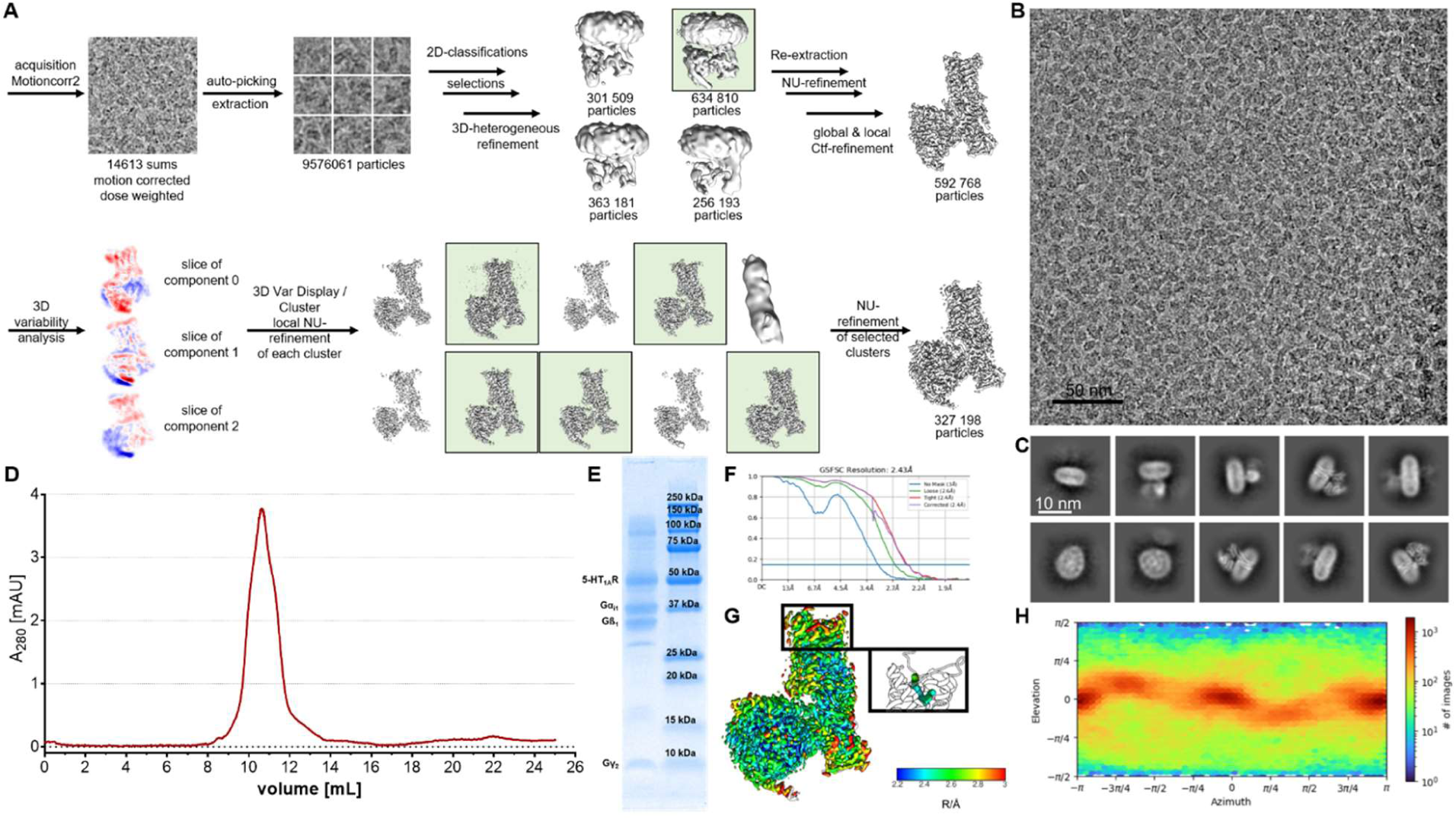
Cryo-EM data processing and complex analytics of the ST171-bound 5-HT_1A_R-Gi1 complex. (A) Cryo-EM single particle analysis workflow, the green subsets were selected for the subsequent steps. (B) Motion corrected and dose weighted average of movie. (C) Selected class averages (10 out of 100). (D) Size-exclusion chromatography profile. (E) SDS- PAGE analysis of the receptor complex. (F) Fourier Shell Correlation. (G) Map filtered and colored according to local resolution with the local resolution of the ligand shown as close up. (H) Distribution of orientation in final map.

**Supplementary Figure 13:**
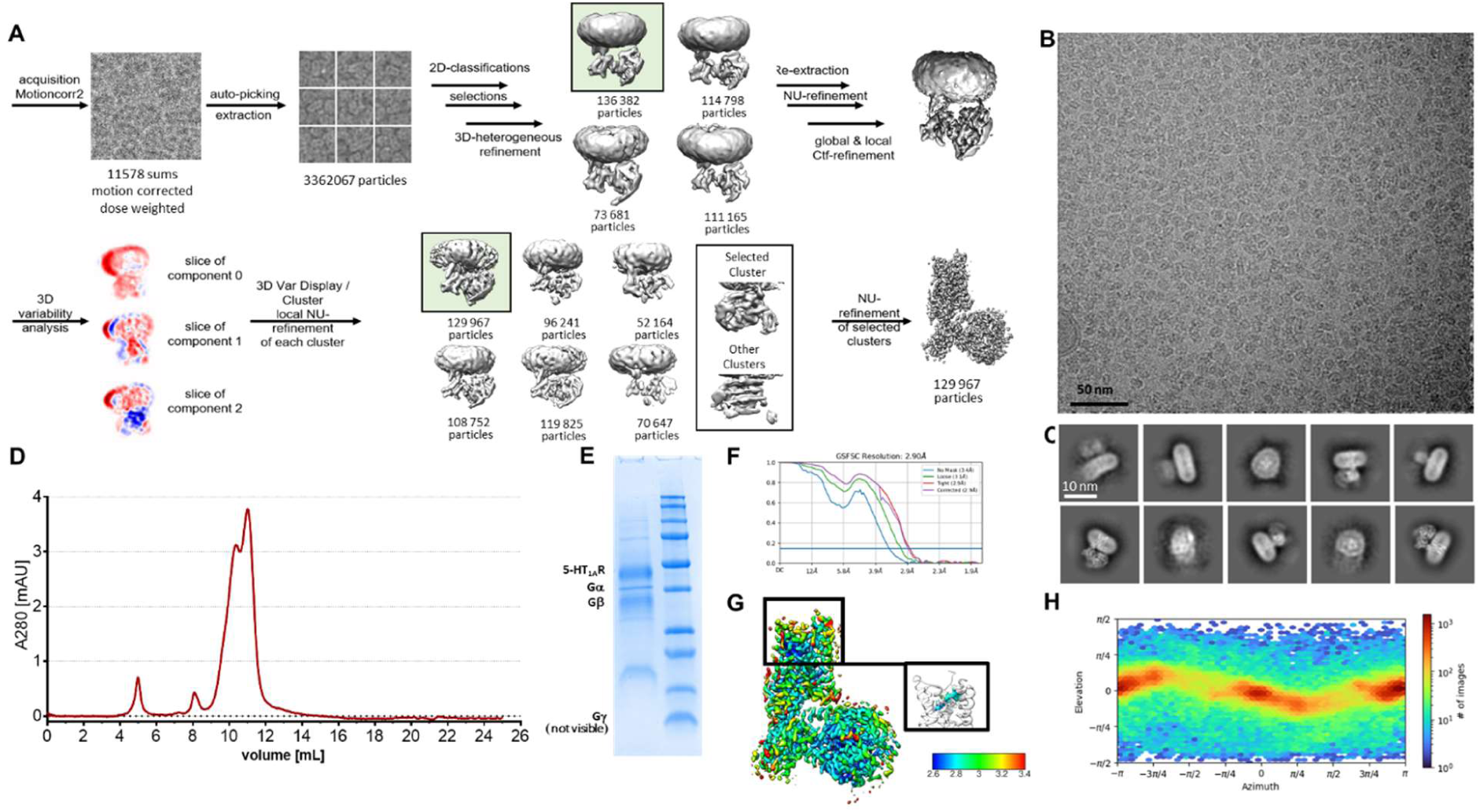
Cryo-EM data processing and complex analytics of the befiradol-bound 5-HT_1A_R-Gi1 complex. (A) Cryo-EM single particle analysis workflow, the green subsets were selected for the subsequent steps. (B) Motion corrected and dose weighted average of movie. (C) Selected class averages (10 out of 100). (D) Size-exclusion chromatography profile. (E) SDS- PAGE analysis of the receptor complex. (F) Fourier Shell Correlation. (G) Map filtered and colored according to local resolution with the local resolution of the ligand shown as close up. (H) Distribution of orientation in final map.

**Supplementary Figure 14:**
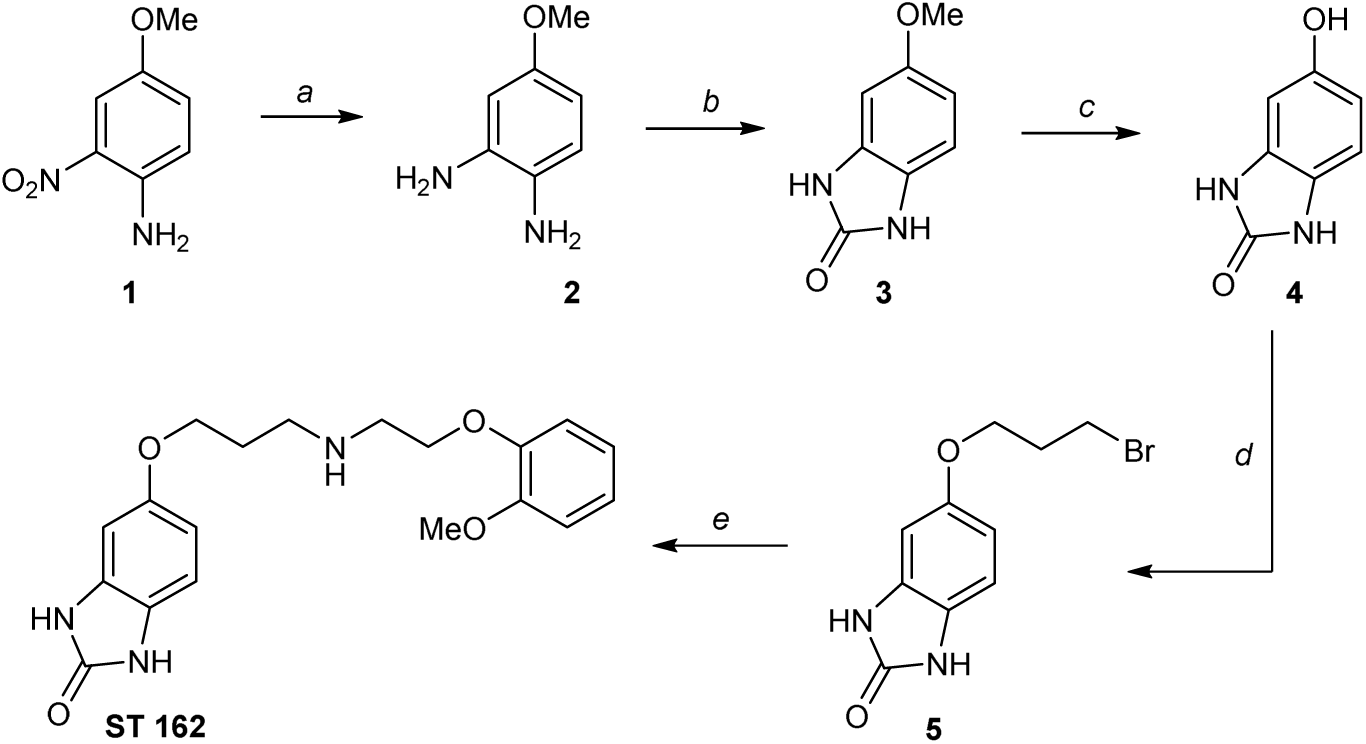
Synthesis of ST162. The synthesis of **ST162** started from 4-methoxy-2-nitroaniline (**1**). Catalytic hydrogenation furnished the respective diamine **2**, which was cyclized CDI to give the benzimidazolone **3**. After cleavage of the methyl ether with BBr_3_, the resulting phenol **4** was alkylated to give the respective bromopropyl ether **5**. In the final step, nucleophilic substitution of **5** with 2-methoxyphenoxyethylamine (*96*) afforded the desired secondary amine **ST 162**. Reagents and conditions: (a) H_2_, Pd/C, methanol, rt, 24 h (78%); (b) 1,1’-carbonyldiimidazole, DMF, Ar, 90°C, 21 h, (68%); (c) 1 M BBr_3_ in CH_2_Cl_2_, CH_2_Cl_2_, Ar, -78°C-rt, 3h (59%); (d) 1,3-dibromopropane, K_2_CO_3_, ethanol, reflux, 3.5 h (30%); (e) 2- methoxyphenoxyethylamine (*96*), KI, CH_3_CN, reflux, 4.5 h (31%).

**Supplementary Figure 15:**
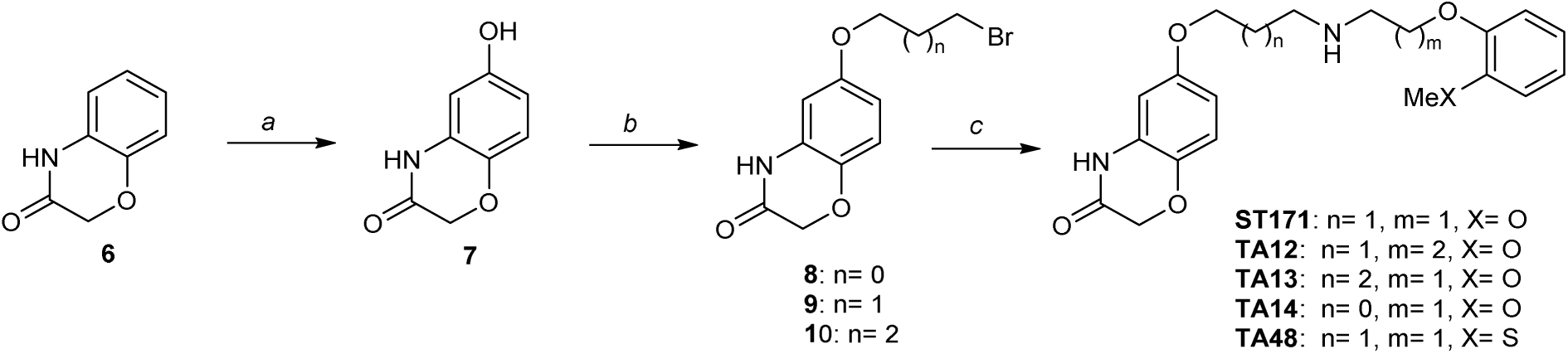
Synthesis of ST171 and analogs thereof. The synthesis of **ST171** and its analogs **TA12**, **TA13**, **TA14** and **TA48** started from 2*H*-benzo[*b*][1,4]oxazin-3(4*H*)-one (**6**), which was oxidized with [bis(trifluoroacetoxy)iodo] benzene to give the respective hydroxyl analogue **7**. Alkylation with different dibromoalkanes afforded the bromoalkyl ethers **8**, **9** and **10**. In the final step, nucleophilic substitution with 2-methoxyphenoxyalkylamine or 2-methylthiophenoxyethylamine afforded the desired secondary amines **ST 171, TA12**, **TA13**, **TA14** and **TA48**. Reagents and conditions: (a) [bis(trifluoroacetoxy)iodo] benzene, trifluoroacetic acid, reflux, 10 min (45%); (b) 1,3-dibromoalkane, K_2_CO_3_, ethanol, reflux, 3.5 h (53-63%); (c) 2-methoxyphenoxyethylamine (*96*) or 3-(2-methoxyphenoxy)propylamine (**11**) or 2-(2-methylthiophenoxy)ethylamine (**12**), KI, CH_3_CN, reflux, 2.5-5 h (39-74%).

**Supplementary Table 1:**
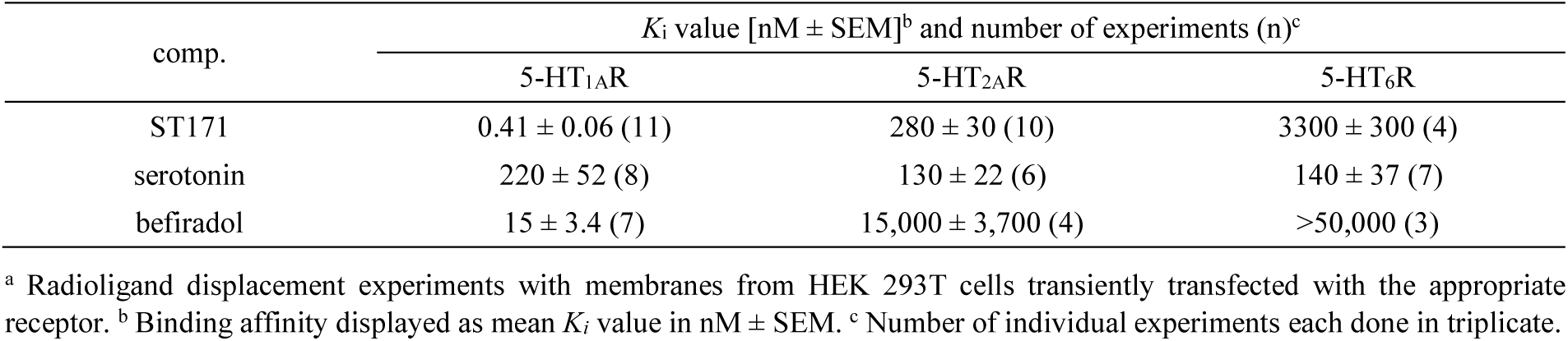
Binding affinity for ST171 and the references serotonin and befiradol to the serotonin receptor subtypes 5-HT_1A_, 5-HT_2A_, and 5-HT_6_.^a^.

**Supplementary Table 2:** Functional activity of ST171 and the references serotonin and befiradol as receptor mediated change of cAMP concentration monitored in different cell systems applying the GloSensor system.

| cell system | comp. | functional activity<sup>a</sup> |  |  |  | n<sup>f</sup> |
| --- | --- | --- | --- | --- | --- | --- |
|  |  | EC<sub>50</sub><sup>left</sup> [nM±SEM]<sup>b</sup> | E<sub>max</sub> [%±SEM]<sup>c</sup> | EC<sub>50</sub><sup>right</sup> [nM±SEM]<sup>d</sup> | E<sub>sat</sub> [%±SEM]<sup>e</sup> |  |
| HEK Δβ-arrestin | ST171 | 3.5 ± 1.0 | 60 ± 3 | --- | --- | 7 |
|  | serotonin | 1.9 ± 0.2 | 100 | 140 ± 44 | 77 ± 3 | 7 |
|  | befiradol | 4.7 ± 0.9 | 89 ± 4 | 330 ± 120 | 73 ± 5 | 7 |
| HEK293A | ST171 | 1.6 ± 0.4 | 78 ± 5 | --- | --- | 5 |
|  | serotonin | 1.4 ± 0.2 | 100 | 360 ± 71 | 82 ± 2 | 7 |
|  | befiradol | 2.9 ± 0.6 | 91 ± 2 | 390 ± 64 | 78 ± 2 | 6 |
| HEK293T | ST171 | 0.90 ± 0.2 | 100 ± 3 | --- | --- | 7 |
|  | serotonin | 0.40 ± 0.1 | 100 | 170 ± 33 | 50 ± 5 | 6 |
|  | befiradol | 1.7 ± 0.6 | 111 ± 6 | 180 ± 57 | 71 ± 6 | 5 |

<sup>a</sup> Functional activity was determined in different HEK cells transiently transfected with 5-HT<sub>1A</sub> and the cAMP Glo-biosensor. <sup>b</sup> Mean EC<sub>50</sub> value representing the potency for the inhibition of cAMP accumulation in [nM±SEM] derived from experiments performed in duplicates (for HEK293T) or triplicates (HEK Δβ-arrestin, HEK293A). <sup>c</sup> Mean value of the maximum activity for the inhibition of cAMP accumulation in [%±SEM] relative to the full effect of serotonin [=100%]. <sup>d</sup> Mean EC<sub>50</sub> value in [nM±SEM] expressing the accumulation of cAMP. <sup>e</sup> Mean E<sub>max</sub> in [%±SEM] relative to the maximum effect of inhibition of cAMP accumulation by serotonin. <sup>f</sup> Number of individual experiments.

**Supplementary Table 3:**
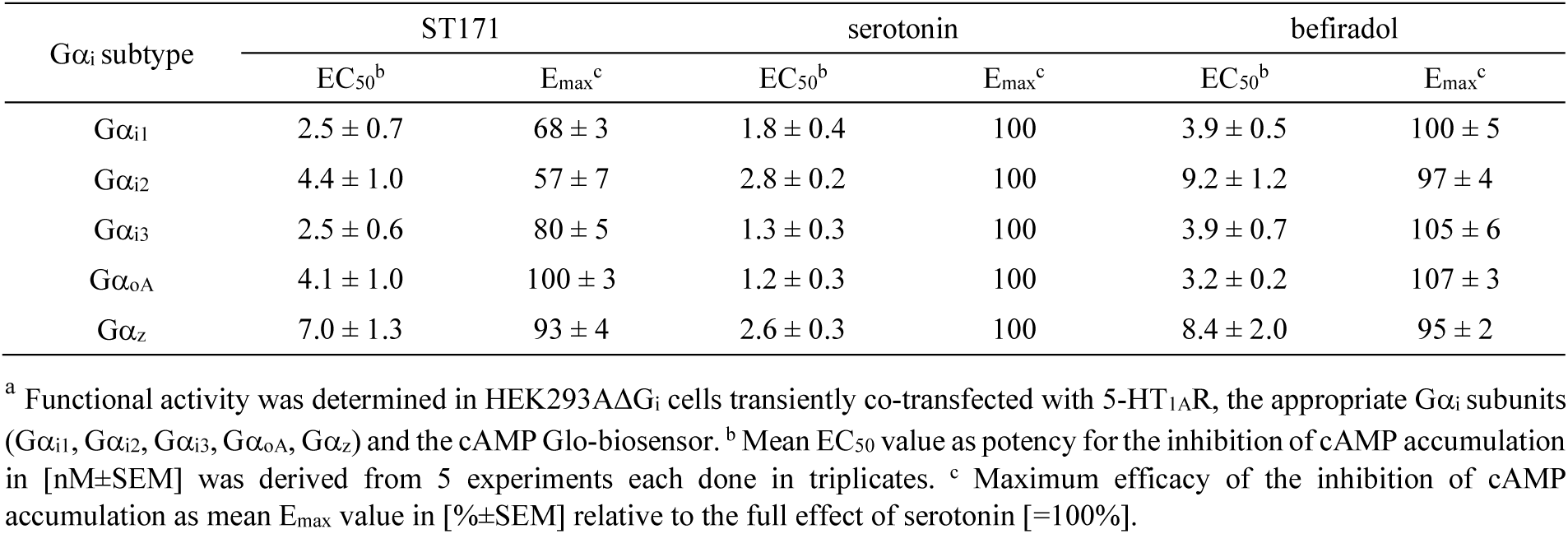
Subtype selective G-protein activation measured as receptor mediated change of cAMP concentration stimulated by serotonin and befiradol.^a^.

**Supplementary Table 4:** Functional activity of ST171, serotonin and befiradol in correlation to the receptor density of 5-HT1A in HEK293T cells.^a^.

| 5-HT<sub>1A</sub>R |  |  | ST171 |  |  | serotonin |  |  | befiradol |  |  |
| --- | --- | --- | --- | --- | --- | --- | --- | --- | --- | --- | --- |
| cDNA<sup>b</sup> | density<sup>c</sup> | n<sup>d</sup> | EC<sub>50</sub><sup>e</sup> | E<sub>max</sub><sup>f</sup> | n<sup>g</sup> | EC<sub>50</sub><sup>e</sup> | E<sub>max</sub><sup>c</sup> | n | EC<sub>50</sub><sup>b</sup> | E<sub>max</sub><sup>c</sup> | n |
| 0.4 μg | 1.5 ± 0.16 | 7 | 1.9 ± 0.31 | 77 ± 3 | 17 | 8.7 ± 1.0 | 100 | 16 | 4.5 ± 0.77 | 102 ± 3 | 12 |
| 0.2 μg | 1.3 ± 0.15 | 4 | 1.6 ± 0.17 | 66 ± 4 | 8 | 7.5 ± 1.5 | 100 | 8 | 7.1 ± 0.90 | 102 ± 2 | 5 |
| 0.1 μg | 0.70 ± 0.094 | 7 | 1.6 ± 0.27 | 62 ± 2 | 15 | 19 ± 2.8 | 100 | 14 | 6.9 ± 1.3 | 101 ± 4 | 10 |
| 0.04 μg | 0.40 ± 0.070 | 8 | 1.2 ± 0.23 | 45 ± 2 | 12 | 27 ± 5.6 | 100 | 12 | 11 ± 2.2 | 103 ± 3 | 10 |
| 0.01 μg | 0.081 ± 0.009 | 5 | 19 ± 10 | 37 ± 5 | 8 | 24 ± 6.3 | 100 | 6 | 11 ± 6.7 | 99 ± 9 | 4 |

<sup>a</sup> Functional activity was determined with the BRET biosensor system applying Gα<sub>i</sub>RLuc and GγGFP. <sup>b</sup> Amount of transfected cDNA. <sup>c</sup> 5-HT<sub>1A</sub>R density expressed as 10<sup>6</sup> receptors per cell was determined in a whole cell saturation binding experiment with the radioligand [<sup>3</sup>H]WAY600,135. <sup>d</sup> Number of individual experiments. <sup>e</sup> Mean EC<sub>50</sub> value for the inhibition of cAMP accumulation in [nM±SEM]. <sup>f</sup> Maximum efficacy of the inhibition of cAMP accumulation in [%±SEM] relative to the full effect of serotonin [=100%]. <sup>g</sup> Number of individual experiments each done in duplicates.

**Supplementary Table 5:** Selectivity profile of binding affinity for ST 171 to a set of class A GPCRs.^a^.

| GPCR | subtype | binding affinity |  |  | experimental conditions |  |  |
| --- | --- | --- | --- | --- | --- | --- | --- |
| | | $K_i$ [nM ± SD]<sup>b</sup> | $pK_i$<sup>c</sup> | n<sup>d</sup> | radioligand | $K_D$<sup>e</sup> | $B_{max}$<sup>f</sup> |
| serotonin | 5-HT<sub>1A</sub> | 0.41 ± 0.19 | 9.433 | 11 | [<sup>3</sup>H]WAY100635 | 0.080 | 3700 |
| | 5-HT<sub>2A</sub> | 280 ± 95 | 6.585 | 10 | [<sup>3</sup>H]ketanserin | 0.29 | 1400 |
| | 5-HT<sub>6</sub> | 3100 ± 5800 | 5.511 | 4 | [<sup>3</sup>H]N-methyl-LSD | 2.3 | 800 |
| dopamine | D<sub>1</sub> | 260 ± 170 | 6.680 | 4 | [<sup>3</sup>H]SCH23390 | 0.29 | 4000 |
| | D<sub>2long</sub><sup>g</sup> | 21 ± 13 | 7.637 | 6 | [<sup>3</sup>H]spiperone | 0.078 | 1300 |
| | D<sub>2long</sub><sup>g</sup> | 8.5 ± 5.0 | 8.007 | 6 | [<sup>3</sup>H]spiperone | 0.080 | 6800 |
| | D<sub>3</sub><sup>g</sup> | 22 ± 11 | 7.583 | 7 | [<sup>3</sup>H]spiperone | 0.16 | 6300 |
| | D<sub>4</sub><sup>g</sup> | 0.17 ± 0.09 | 9.742 | 6 | [<sup>3</sup>H]spiperone | 0.20 | 1200 |
| | D<sub>5</sub> | 740 ± 410 | 6.188 | 4 | [<sup>3</sup>H]SCH23390 | 0.26 | 1400 |
| adrenoceptor | $\alpha_{1A}$ | 0.49 ± 0.24 | 9.361 | 9 | [<sup>3</sup>H]prazosine | 0.35 | 3900 |
| | $\alpha_{1B}$ | 6.7 ± 2.0 | 8.191 | 4 | [<sup>3</sup>H]prazosine | 0.11 | 3600 |
| | $\alpha_{2A}$ | 260 ± 210 | 6.690 | 4 | [<sup>3</sup>H]RX821002 | 0.48 | 1500 |
| | $\alpha_{2B}$ | 110 ± 60 | 6.989 | 4 | [<sup>3</sup>H]RX821002 | 1.75 | 3900 |
| | $\alpha_{2C}$ | 28 ± 12 | 7.587 | 4 | [<sup>3</sup>H]RX821002 | 0.60 | 7000 |
| | $\beta_1$ | 1700 ± 350 | 5.788 | 2 | [<sup>3</sup>H]CGP12177 | 0.080 | 3500 |
| | $\beta_2$ | 100 ± 11 | 6.992 | 4 | [<sup>3</sup>H]CGP12177 | 0.095 | 1500 |
| muscarinic | M<sub>1</sub> | 10000 ± 920 | 4.986 | 2 | [<sup>3</sup>H]NMS | 0.050 | 5600 |
| | M<sub>2</sub> | 18000 ± 9200 | 4.789 | 2 | [<sup>3</sup>H]NMS | 0.18 | 1500 |
| | M<sub>3</sub> | 49000 ± 28000 | 4.382 | 3 | [<sup>3</sup>H]NMS | 0.18 | 3500 |
| opioid | μOR | 8600 ± 2100 | 5.074 | 2 | [<sup>3</sup>H]diprenorphine | 0.080 | 1800 |
| | δOR | 24000 ± 3500 | 4.631 | 2 | [<sup>3</sup>H]diprenorphine | 0.19 | 2500 |
| | κOR | 6200 ± 850 | 5.202 | 2 | [<sup>3</sup>H]diprenorphine | 0.070 | 4500 |
| orexin | OX<sub>1</sub> | >50000 | <4.300 | 2 | [<sup>3</sup>H]SB674042 | 0.62 | 8300 |
|  | OX<sub>2</sub> | >50000 | <4.300 | 2 | [<sup>3</sup>H]EMPA | 1.1 | 7000 |
| neurotensin | NTS<sub>1</sub> | >50000 | <4.300 | 2 | [<sup>3</sup>H]NT(8-13)<sup>i</sup> | 0.70 | 10000 |
|  | NTS<sub>2</sub> | >50000 | <4.300 | 2 | [<sup>3</sup>H]NT(8-13)<sup>i</sup> | 1.8 | 700 |

<sup>a</sup> Radioligand displacement experiments with membranes from HEK 293T cells transiently transfected with the appropriate receptor. <sup>b</sup> Binding affinity displayed as mean $K_i$ value in nM ± SD. <sup>c</sup> Binding affinities displayed as $pK_i$ . <sup>d</sup> Number of individual experiments each done in triplicate. <sup>e</sup> Binding affinity for the radioligand as $K_D$ value in nM. <sup>f</sup> Receptor density as $B_{max}$ in fmol/mg protein. <sup>g</sup> Dopamine D<sub>2long</sub>, D<sub>2short</sub> and D<sub>3</sub> receptors were stably expressed in CHO cells. <sup>h</sup> The receptor isoform D<sub>4.4</sub> was stably expressed in CHO cells. <sup>i</sup> [<sup>3</sup>H]NT8-13 was purchased as a customs synthesis by Novandi, Sönderdälje, Sweden.

**Supplementary Table 6:** Functional activity of ST171 for the adrenergic receptors α_1A_, α_1B_, α_2A_, α_2B_, and α_2C_ in comparison to norepinephrine and for the dopamine receptor subtype D_4_ relative to quinpirole.^a^.

| adrenergic | ST171 |  |  | norepinephrine |  |  |
| --- | --- | --- | --- | --- | --- | --- |
|  | EC<sub>50</sub> [nM±SEM]<sup>b</sup> | E<sub>max</sub> [%]<sup>c</sup> | n<sup>d</sup> | EC<sub>50</sub> [nM±SEM]<sup>b</sup> | E<sub>max</sub> [%]<sup>c</sup> | n<sup>d</sup> |
| $\alpha_{1A}$ | n.d. | <5 | 4 | 29 ± 5.2 | 100 | 5 |
| $\alpha_{1B}$ | n.d. | <5 | 3 | 18 ± 4.0 | 100 | 7 |
| $\alpha_{2A}$ | n.d. | <5 | 4 | 1.0 ± 0.08 | 100 | 14 |
| $\alpha_{2B}$ | n.d. | <5 | 3 | 2.4 ± 0.80 | 100 | 5 |
| $\alpha_{2C}$ | 3300 ± 1500 | 14 ± 1 | 4 | 0.60 ± 0.18 | 100 | 5 |
| dopaminergic | ST171 |  |  | quinpirole |  |  |
|  | EC<sub>50</sub> [nM±SEM]<sup>b</sup> | E<sub>max</sub> [%]<sup>c</sup> | n<sup>d</sup> | EC<sub>50</sub> [nM±SEM]<sup>b</sup> | E<sub>max</sub> [%]<sup>c</sup> | n<sup>d</sup> |
| D<sub>4</sub> | 1.6 ± 0.22 | 42 ± 3 | 7 | 15 ± 1.8 | 100 | 8 |

<sup>a</sup> Functional activity was determined in HEK293T cells measuring G $\alpha_q$ activation (by $\alpha_{1A}$ and $\alpha_{1B}$ ) when applying an IP accumulation assay. G $\alpha_i$ activation (stimulated by $\alpha_{2A}$ , $\alpha_{2B}$ , $\alpha_{2C}$ , and D<sub>4</sub>) was monitored with a BRET biosensor system based on G $\alpha_{i1}$ -G $\gamma_1$ dissociation. <sup>b</sup> Mean EC<sub>50</sub> values in [nM±SEM]. <sup>c</sup> Maximum efficacy as mean E<sub>max</sub> value in [%±SEM] relative to the full effect of norepinephrine (for adrenergic receptors) or quinpirole (D<sub>4</sub>). <sup>d</sup> Number of individual experiments each done in duplicates. n.d.: could not be analyzed.

**Supplementary Table 7:** Binding affinity and functional activity for the ST171 derivatives TA12, TA13, TA14, and TA48 at the 5-HT_1A_R.

| comp. | binding affinity<sup>a</sup> |  | inhibition of cAMP accumulation<sup>b</sup> |  |  | β-arrestin recruitment<sup>c</sup> |  |
| --- | --- | --- | --- | --- | --- | --- | --- |
|  | K<sub>i</sub> [nM ± SEM]<sup>d</sup> | n<sup>e</sup> | EC<sub>50</sub><sup>f</sup> | E<sub>max</sub><sup>g</sup> | n<sup>e</sup> | EC<sub>50</sub><sup>h</sup> | E<sub>max</sub><sup>g</sup> |
| TA12 | 7.8 ± 0.36 | 3 | 130 ± 60 | 23 ± 5 | 4 | n.d. | < 5 |
| TA13 | 0.49 ± 0.089 | 3 | 7.3 ± 1.8 | 68 ± 4 | 4 | n.d. | < 5 |
| TA14 | 20 ± 3.0 | 3 | 140 ± 10 | 27 ± 4 | 4 | n.d. | < 5 |
| TA48 | 0.98 ± 0.079 | 3 | 18 ± 7.0 | 57 ± 4 | 4 | n.d. | < 5 |

<sup>a</sup> Binding affinity was determined in a radioligand displacement assay with membranes from HEK293T cell transiently transfected with 5-HT<sub>1A</sub>R and the radioligand [<sup>3</sup>H]WAY600,135. <sup>b</sup> G-protein signaling evaluated with a cAMP accumulation assay with the biosensor CAMYEL. <sup>c</sup> β-Arrestin 2 recruitment was measured applying the PathHunter assay with 5-HT<sub>1A</sub>R-ARMS2-PK2 in HEK cells stably expressing β-arrestin-2-EA. <sup>d</sup> K<sub>i</sub> in [nM ± SEM] were derived from experiments each done in triplicates. <sup>e</sup> Number of individual experiments. <sup>f</sup> Mean EC<sub>50</sub> values in [nM±SEM]. <sup>g</sup> Maximum efficacy as mean E<sub>max</sub> value in [% ± SEM] relative to the full effect of serotonin. <sup>h</sup> Mean EC<sub>50</sub> values in [nM±SEM] derived from 3 single experiments each done in duplicates. n.d.: could not be analyzed.

## References

1. E. Bruera, E. Del Fabbro, Pain management in the era of the opioid crisis. Am. Soc. Clin. Oncol. Educ. Book, 807–812 (2018).

2. S. E. Nadeau, J. K. Wu, R. A. Lawhern, Opioids and chronic pain: An analytic review of the clinical evidence. Fron. Pain. Res. (Lausanne) 2, 721357 (2021).

3. S. L. Walsh, K. L. Preston, G. E. Bigelow, M. L. Stitzer, Acute administration of buprenorphine in humans: partial agonist and blockade effects. J. Pharmacol. Exp. Ther. 274, 361–372 (1995).

4. A. Manglik et al., Structure-based discovery of opioid analgesics with reduced side effects. Nature 537, 185–190 (2016).

5. H. Wang et al., Structure-based evolution of G protein-biased μ-opioid receptor agonists. Angew. Chem. Int. Ed. 61, e202200269 (2022).

6. P. Celada, A. Bortolozzi, F. Artigas, Serotonin 5-HT1A receptors as targets for agents to treat psychiatric disorders: Rationale and current status of research. CNS Drugs 27, 703–716 (2013).

7. J. Gjerstad, A. Tjølsen, K. Hole, The effect of 5-HT1A receptor stimulation on nociceptive dorsal horn neurones in rats. Eur. J. Pharmacol. 318, 315–321 (1996).

8. R. Bardoni, Serotonergic modulation of nociceptive circuits in spinal cord dorsal horn. Curr. Neuropharmacol. 17, 1133–1145 (2019).

9. X.-J. Xu, F. Colpaert, Z. Wiesenfeld-Hallin, Opioid hyperalgesia and tolerance versus 5-HT1A receptor-mediated inverse tolerance. Trends Pharmacol. Sci. 24, 634–639 (2003).

10. D. J. Haleem, Serotonin-1A receptor dependent modulation of pain and reward for improving therapy of chronic pain. Pharmacol. Res. 134, 212–219 (2018).

11. J. Ren, X. Ding, J. J. Greer, 5-HT1A receptor agonist befiradol reduces fentanyl-induced respiratory depression, analgesia, and sedation in rats. Anesthesiology 122, 424–434 (2015).

12. A. Newman-Tancredi et al., NLX-112, a highly selective 5-HT(1A) receptor agonist, mediates analgesia and antidepressant-like activity in rats via spinal cord and prefrontal cortex 5-HT(1A) receptors, respectively. Brain Res. 1688, 1–7 (2018).

13. K. Deseure, S. Bréand, F. C. Colpaert, Curative-like analgesia in a neuropathic pain model: Parametric analysis of the dose and the duration of treatment with a high-efficacy 5-HT1A receptor agonist. Eur. J. Pharmacol. 568, 134–141 (2007).

14. R. Fisher et al., The selective 5-HT1A receptor agonist, NLX-112, exerts anti-dyskinetic and anti- parkinsonian-like effects in MPTP-treated marmosets. Neuropharmacology 167, 107997 (2020).

15. E. A. Fink et al., Structure-based discovery of nonopioid analgesics acting through the α2A-adrenergic receptor. Science 377, eabn7065 (2022).

16. J. R. Raymond, Y. V. Mukhin, T. W. Gettys, M. N. Garnovskaya, The recombinant 5-HT1A receptor: G protein coupling and signalling pathways. Br. J. Pharmacol. 127, 1751–1764 (1999).

17. M. O’Hayre et al., Genetic evidence that β-arrestins are dispensable for the initiation of β(2)-adrenergic receptor signaling to ERK. Sci. Signaling 10, eaal3395 (2017).

18. M. Ui, Islet-activating protein, pertussis toxin: a probe for functions of the inhibitory guanine nucleotide regulatory component of adenylate cyclase. Trends Pharmacol. Sci. 5, 277–279 (1984).

19. R. E. West, J. Moss, M. Vaughan, T. Liu, T. Y. Liu, Pertussis toxin-catalyzed ADP-ribosylation of transducin. Cysteine 347 is the ADP-ribose acceptor site. J. Biol. Chem. 260, 14428–14430 (1985).

20. W. Stallaert et al., Purinergic receptor transactivation by the β(2)-adrenergic receptor increases intracellular Ca(2+) in nonexcitable cells. Mol. Pharmacol. 91, 533–544 (2017).

21. Y. Ono et al., Generation of Gαi knock-out HEK293 cells illuminates Gαi-coupling diversity of GPCRs. *Commun*. Biol. 6, 112 (2023).

22. Y. Namkung et al., Monitoring G protein-coupled receptor and β-arrestin trafficking in live cells using enhanced bystander BRET. Nat. Commun. 7, 12178 (2016).

23. K. A. Farzam K, Lakhkar AD, Adrenergic drugs. National Library of Medicine (Internet), StatPearls Publishing 2023, (2022).

24. M. J. Millan, Descending control of pain. Prog. Neurobiol. 66, 355–474 (2002).

25. Q. Q. Liu et al., Role of 5-HT receptors in neuropathic pain: potential therapeutic implications. Pharmacol. Res. 159, 104949 (2020).

26. A. Dray, Inflammatory mediators of pain. Br. J. Anaesth. 75, 125–131 (1995).

27. K. Okamoto et al., 5-HT2A receptor subtype in the peripheral branch of sensory fibers is involved in the potentiation of inflammatory pain in rats. Pain 99, 133–143 (2002).

28. N. M. E. Carmichael, M. P. Charlton, J. O. Dostrovsky, Activation of the 5-HT1B/D receptor reduces hindlimb neurogenic inflammation caused by sensory nerve stimulation and capsaicin. Pain 134, 97–105 (2008).

29. K. P. Zeitz et al., The 5-HT3 subtype of serotonin receptor contributes to nociceptive processing via a novel subset of myelinated and unmyelinated nociceptors. J. Neurosci. 22, 1010 (2002).

30. M. Sasaki, H. Obata, K. Kawahara, S. Saito, F. Goto, Peripheral 5-HT2A receptor antagonism attenuates primary thermal hyperalgesia and secondary mechanical allodynia after thermal injury in rats. Pain 122, 130–136 (2006).

31. V. Kayser, B. Aubel, M. Hamon, S. Bourgoin, The antimigraine 5-HT1B/1D receptor agonists, sumatriptan, zolmitriptan and dihydroergotamine, attenuate pain-related behaviour in a rat model of trigeminal neuropathic pain. Br. J. Pharmacol. 137, 1287–1297 (2002).

32. T. Nikai, A. I. Basbaum, A. H. Ahn, Profound reduction of somatic and visceral pain in mice by intrathecal administration of the anti-migraine drug, sumatriptan. Pain 139, 533–540 (2008).

33. A. Dogrul, M. H. Ossipov, F. Porreca, Differential mediation of descending pain facilitation and inhibition by spinal 5HT-3 and 5HT-7 receptors. Brain Res. 1280, 52–59 (2009).

34. D. Hoyer, J. P. Hannon, G. R. Martin, Molecular, pharmacological and functional diversity of 5-HT receptors. Pharmacol. Biochem. Behav. 71, 533–554 (2002).

35. M. J. Millan, L. Seguin, P. Honoré, S. Girardon, K. Bervoets, Pro- and antinociceptive actions of serotonin (5-HT)1A agonists and antagonists in rodents: relationship to algesiometric paradigm. Behav. Brain Res. 73, 69–77 (1995).

36. L. Bardin, F. C. Colpaert, Role of spinal 5-HT1A receptors in morphine analgesia and tolerance in rats. *Eur. J. Pain (Oxford*, U. K.) 8, 253–261 (2004).

37. H.-J. Jeong, V. A. Mitchell, C. W. Vaughan, Role of 5-HT1 receptor subtypes in the modulation of pain and synaptic transmission in rat spinal superficial dorsal horn. Br. J. Pharmacol. 165, 1956–1965 (2012).

38. K. Sałat et al., Antinociceptive, antiallodynic and antihyperalgesic effects of the 5-HT1A receptor selective agonist, NLX-112 in mouse models of pain. Neuropharmacology 125, 181–188 (2017).

39. L. Bardin, M. Bardin, J. Lavarenne, A. Eschalier, Effect of intrathecal serotonin on nociception in rats: influence of the pain test used. Exp. Brain Res. 113, 81–87 (1997).

40. R. Nadeson, C. S. Goodchild, Antinociceptive role of 5-HT1A receptors in rat spinal cord. Br. J. Anaesth. 88, 679–684 (2002).

41. C. Schmauss, D. L. Hammond, J. W. Ochi, T. L. Yaksh, Pharmacological antagonism of the antinociceptive effects of serotonin in the rat spinal cord. Eur. J. Pharmacol. 90, 349–357 (1983).

42. N. Mjellem, A. Lund, P. K. Eide, R. Størkson, A. Tjølsen, The role of 5-HT1A and 5-HT1B receptors in spinal nociceptive transmission and in the modulation of NMDA induced behaviour. Neuroreport 3, 1061–1064 (1992).

43. X. Khawaja, Quantitative autoradiographic characterisation of the binding of [3H]WAY-100635, a selective 5-HT1A receptor antagonist. Brain Res. 673, 217–225 (1995).

44. H. K. Kia et al., Immunocytochemical localization of serotonin1A receptors in the rat central nervous system. J. Comp. Neurol. 365, 289–305 (1996).

45. G. Daval, D. Vergé, A. I. Basbaum, S. Bourgoin, M. Hamon, Autoradiographic evidence of serotonin1 binding sites on primary afferent fibres in the dorsal horn of the rat spinal cord. Neurosci. Lett. 83, 71–76 (1987).

46. A. Levit Kaplan et al., Structure-based design of a chemical probe set for the 5-HT5A serotonin receptor. J. Med. Chem. 65, 4201–4217 (2022).

47. S. D. Shields, W. A. Eckert, A. I. Basbaum, Spared nerve injury model of neuropathic pain in the mouse: a behavioral and anatomic analysis. J. Pain 4, 465–470 (2003).

48. T. J. Pucadyil, A. Chattopadhyay, Cholesterol modulates ligand binding and G-protein coupling to serotonin(1A) receptors from bovine hippocampus. Biochim. Biophys. Acta 1663, 188–200 (2004).

49. W. Yin et al., Crystal structure of the human 5-HT1B serotonin receptor bound to an inverse agonist. Cell Discovery 4, 12 (2018).

50. P. Xu et al., Structural insights into the lipid and ligand regulation of serotonin receptors. Nature 592, 469–473 (2021).

51. S. Huang et al., GPCRs steer Gi and Gs selectivity via TM5-TM6 switches as revealed by structures of serotonin receptors. Mol. Cell 82, 2681–2695.e2686 (2022).

52. C. Cao et al., Signaling snapshots of a serotonin receptor activated by the prototypical psychedelic LSD. Neuron 110, 3154–3167.e3157 (2022).

53. H. S. Biswal, S. Wategaonkar, Nature of the N−H···S Hydrogen Bond. J. Phys. Chem. A 113, 12763–12773 (2009).

54. A. J. Kooistra, S. Kuhne, I. J. P. de Esch, R. Leurs, C. de Graaf, A structural chemogenomics analysis of aminergic GPCRs: lessons for histamine receptor ligand design. Br. J. Pharmacol. 170, 101–126 (2013).

55. Q. Qu et al., Insights into distinct signaling profiles of the µOR activated by diverse agonists. Nat. Chem. Biol. 19, 423–430 (2023).

56. A. Newman-Tancredi, R. Y. Depoortère, M. S. Kleven, M. Kołaczkowski, L. Zimmer, Translating biased agonists from molecules to medications: Serotonin 5-HT1A receptor functional selectivity for CNS disorders. Pharmacol. Ther. 229, 107937 (2022).

57. P. K. Eide, K. Hole, Interactions between serotonin and substance P in the spinal regulation of nociception. Brain Res. 550, 225–230 (1991).

58. A. Brenchat et al., 5-HT7 receptor activation inhibits mechanical hypersensitivity secondary to capsaicin sensitization in mice. Pain 141, 239–247 (2009).

59. H. Hübner, C. Haubmann, W. Utz, P. Gmeiner, Conjugated enynes as nonaromatic catechol bioisosteres: Synthesis, binding experiments, and computational studies of novel dopamine receptor agonists recognizing preferentially the D3 subtype. J. Med. Chem. 43, 756–762 (2000).

60. O. H. Lowry, N. J. Rosebrough, A. L. Farr, R. J. Randall, Protein measurement with the folin phenol reagent. J. Biol. Chem. 193, 265–275 (1951).

61. Y.-C. Cheng, W. H. Prusoff, Relationship between the inhibition constant (KI) and the concentration of inhibitor which causes 50 per cent inhibition (I50) of an enzymatic reaction. Biochem. Pharmacol. 22, 3099–3108 (1973).

62. L. I. Jiang et al., Use of a cAMP BRET sensor to characterize a novel regulation of cAMP by the sphingosine 1-phosphate/G13 pathway. J. Biol. Chem. 282, 10576–10584 (2007).

63. J. Köckenberger et al., Synthesis, characterization, and application of muscarinergic M3 receptor ligands linked to fluorescent dyes. J. Med. Chem. 65, 16494–16509 (2022).

64. C. Galés et al., Probing the activation-promoted structural rearrangements in preassembled receptor– G protein complexes. Nat. Struct. Mol. Biol. 13, 778–786 (2006).

65. D. Möller et al., Discovery of G protein-biased dopaminergics with a pyrazolo[1,5-a]pyridine substructure. J. Med. Chem. 60, 2908–2929 (2017).

66. S. R. Chaplan, F. W. Bach, J. W. Pogrel, J. M. Chung, T. L. Yaksh, Quantitative assessment of tactile allodynia in the rat paw. J. Neurosci. Methods 53, 55–63 (1994).

67. Y.-L. Liang et al., Dominant negative G proteins enhance formation and purification of agonist-GPCR-G protein complexes for structure determination. ACS Pharmacol. Transl. Sci. 1, 12–20 (2018).

68. S. Q. Zheng et al., MotionCor2: anisotropic correction of beam-induced motion for improved cryo-electron microscopy. Nat. Methods 14, 331–332 (2017).

69. A. Punjani, J. L. Rubinstein, D. J. Fleet, M. A. Brubaker, cryoSPARC: algorithms for rapid unsupervised cryo-EM structure determination. Nat. Methods 14, 290–296 (2017).

70. D. Kimanius, L. Dong, G. Sharov, T. Nakane, S. H. W. Scheres, New tools for automated cryo-EM single-particle analysis in RELION-4.0. Biochem. J. 478, 4169–4185 (2021).

71. D. Liebschner et al., Macromolecular structure determination using X-rays, neutrons and electrons: recent developments in Phenix. *Acta Crystallogr.*, Sect. D: Struct. Biol. 75, 861–877 (2019).

72. E. F. Pettersen et al., UCSF Chimera - a visualization system for exploratory research and analysis. J. Comput. Chem. 25, 1605–1612 (2004).

73. P. Emsley, B. Lohkamp, W. G. Scott, K. Cowtan, Features and development of Coot. Acta Crystallogr., Sect. D: Biol. Crystallogr. 66, 486–501 (2010).

74. P. D. Adams et al., PHENIX: a comprehensive Python-based system for macromolecular structure solution. *Acta Crystallogr.*, Sect. D: Biol. Crystallogr. 66, 213–221 (2010).

75. A. Šali, T. L. Blundell, Comparative protein modelling by satisfaction of spatial restraints. J. Mol. Biol. 234, 779–815 (1993).

76. R. Anandakrishnan, B. Aguilar, A. V. Onufriev, H++ 3.0: automating pK prediction and the preparation of biomolecular structures for atomistic molecular modeling and simulations. Nucleic Acids Res. 40, W537–W541 (2012).

77. A. Ranganathan, R. O. Dror, J. Carlsson, Insights into the role of Asp79 2.50 in β2 adrenergic receptor activation from molecular dynamics simulations. Biochemistry 53, 7283–7296 (2014).

78. D. A. Case et al., Amber 2022. University of California, San Francisco, (2022).

79. M. R. Shirts et al., Lessons learned from comparing molecular dynamics engines on the SAMPL5 dataset. J. Comput. Aided Mol. Des. 31, 147–161 (2017).

80. M. A. Lomize, I. D. Pogozheva, H. Joo, H. I. Mosberg, A. L. Lomize, OPM database and PPM web server: resources for positioning of proteins in membranes. Nucleic Acids Res. 40, D370–D376 (2012).

81. M. G. Wolf, M. Hoefling, C. Aponte-Santamaría, H. Grubmüller, G. Groenhof, g_membed: Efficient insertion of a membrane protein into an equilibrated lipid bilayer with minimal perturbation. J. Comput. Chem. 31, 2169–2174 (2010).

82. J. Wang, R. M. Wolf, J. W. Caldwell, P. A. Kollman, D. A. Case, Development and testing of a general amber force field. J. Comput. Chem. 25, 1157–1174 (2004).

83. C. J. Dickson et al., Lipid14: The amber lipid force field. J. Chem. Theory Comput. 10, 865–879 (2014).

84. J. A. Maier et al., ff14SB: Improving the accuracy of protein side chain and backbone parameters from ff99SB. J. Chem. Theory Comput. 11, 3696–3713 (2015).

85. H. J. C. Berendsen, J. R. Grigera, T. P. Straatsma, The missing term in effective pair potentials. J. Phys. Chem. 91, 6269–6271 (1987).

86. C. I. Bayly, P. Cieplak, W. Cornell, P. A. Kollman, A well-behaved electrostatic potential based method using charge restraints for deriving atomic charges: the RESP model. J. Phys. Chem. 97, 10269–10280 (1993).

87. D. Van Der Spoel et al., GROMACS: Fast, flexible, and free. J. Comput. Chem. 26, 1701–1718 (2005).

88. B. Hess, H. Bekker, H. J. C. Berendsen, J. G. E. M. Fraaije, LINCS: A linear constraint solver for molecular simulations. J. Comput. Chem. 18, 1463–1472 (1997).

89. T. Darden, D. York, L. Pedersen, Particle mesh Ewald: An N⋅log(N) method for Ewald sums in large systems. J. Chem. Phys. 98, 10089–10092 (1993).

90. M. Bonomi et al., Promoting transparency and reproducibility in enhanced molecular simulations. Nat. Methods 16, 670–673 (2019).

91. W. Humphrey, A. Dalke, K. Schulten, VMD: Visual molecular dynamics. J. Mol. Graphics 14, 33–38 (1996).

92. D. R. Roe, T. E. Cheatham, III, PTRAJ and CPPTRAJ: Software for Processing and Analysis of Molecular Dynamics Trajectory Data. J. Chem. Theory Comput. 9, 3084–3095 (2013).

93. J. D. Hunter, Matplotlib: A 2D Graphics Environment. Comput. Sci. Eng. 9, 90–95 (2007).

94. M. Waskom, Seaborn: Statistical data visualization. JOSS 6, 3021 (2021).

95. G. P. Moss, Basic terminology of stereochemistry (IUPAC Recommendations 1996). Pure Appl. Chem. 68, 2193–2222 (1996).

96. C. Materne, I. Khan, Method for preparing 2-alkoxyphenoxyethanamines from 2- alkoxyphenoxyethylacetamides, WO 03095416(A1)2003 (2003).

